# Potent Human Broadly SARS-CoV-2 Neutralizing IgA and IgG Antibodies Effective Against Omicron BA.1 and BA.2

**DOI:** 10.1101/2022.04.01.486719

**Authors:** Cyril Planchais, Ignacio Fernández, Timothée Bruel, Guilherme Dias de Melo, Matthieu Prot, Maxime Beretta, Pablo Guardado-Calvo, Jérémy Dufloo, Luis M. Molinos-Albert, Marija Backovic, Jeanne Chiaravalli, Emilie Giraud, Benjamin Vesin, Laurine Conquet, Ludivine Grzelak, Delphine Planas, Isabelle Staropoli, Florence Guivel-Benhassine, Mikaël Boullé, Minerva Cervantes-Gonzalez, French COVID Cohort Study Group, CORSER Study Group, Marie-Noëlle Ungeheuer, Pierre Charneau, Sylvie van der Werf, Fabrice Agou, Jordan D. Dimitrov, Etienne Simon-Lorière, Hervé Bourhy, Xavier Montagutelli, Félix A. Rey, Olivier Schwartz, Hugo Mouquet

**Author notes:** Equal contribution (Timothée Bruel, Guilherme Dias de Melo, Matthieu Prot). These senior authors contributed equally (Etienne Simon-Lorière, Hervé Bourhy, Xavier Montagutelli, Félix A. Rey, Olivier Schwartz). Correspondence (H.M.); (F.A.R.).

## Abstract

Memory B-cell and antibody responses to the SARS-CoV-2 spike protein contribute to long-term immune protection against severe COVID-19, which can also be prevented by antibody-based interventions. Here, wide SARS-CoV-2 immunoprofiling in COVID-19 convalescents combining serological, cellular and monoclonal antibody explorations, revealed humoral immunity coordination. Detailed characterization of a hundred SARS-CoV-2 spike memory B-cell monoclonal antibodies uncovered diversity in their repertoire and antiviral functions. The latter were influenced by the targeted spike region with strong Fc-dependent effectors to the S2 subunit and potent neutralizers to the receptor binding domain. Amongst those, Cv2.1169 and Cv2.3194 antibodies cross-neutralized SARS-CoV-2 variants of concern including Omicron BA.1 and BA.2. Cv2.1169, isolated from a mucosa-derived IgA memory B cell, demonstrated potency boost as IgA dimers and therapeutic efficacy as IgG antibodies in animal models. Structural data provided mechanistic clues to Cv2.1169 potency and breadth. Thus, potent broadly neutralizing IgA antibodies elicited in mucosal tissues can stem SARS-CoV-2 infection, and Cv2.1169 and Cv2.3194 are prime candidates for COVID-19 prevention and treatment.

## Introduction

The coronavirus disease 2019 (COVID-19) is caused by the severe acute respiratory syndrome coronavirus 2 (SARS-CoV-2), and accounts to date for nearly 480 million infection cases and 6 million deaths worldwide (WHO, 2022). SARS-CoV-2 infects host cells through interactions of its surface envelope protein, or spike, with the cellular angiotensin-converting enzyme 2 (ACE2) receptor (Hoffmann et al., 2020; Lan et al., 2020). The SARS-CoV-2 spike (S) is a homo-trimeric glycoprotein with each protomer composed of subunits S1 and S2 (Ke et al., 2020; Walls et al., 2020; Wrapp et al., 2020). S1 contains the N-terminal domain (NTD) and the receptor binding domain (RBD) that interacts with ACE2, while S2 mediates viral fusion (Lan et al., 2020; Yan et al., 2020). Antibodies rapidly develop in response to SARS-CoV-2 infection (Long et al., 2020; Sette and Crotty, 2021), including neutralizing antibodies recognizing distinct S protein regions (Schmidt et al., 2021). The RBD is the primary target of neutralizing antibodies including potent neutralizers, albeit the NTD and S2 stem region also contain neutralizing epitopes (Andreano et al., 2021; Brouwer et al., 2020; Chi et al., 2020; Ju et al., 2020; Liu et al., 2020; Pinto et al., 2021; Rogers et al., 2020; Wec et al., 2020; Zost et al., 2020a). SARS-CoV-2 neutralizing IgA antibodies, detected as early as a week after onset of symptoms, contribute to seroneutralization and can be as potent as IgGs (Sterlin et al., 2021; Wang et al., 2021b). Neutralizing antibodies are the main correlate of protection for COVID-19 vaccines (Krammer, 2021). Still, SARS-CoV-2 spike-specific antibodies, including non-neutralizers, can exert antiviral Fc-dependent effector functions important for *in vivo* protection *i*.*e*., antibody-dependent cellular cytotoxicity (ADCC), and phagocytosis (ADCP) (Chertow et al., 2021; Dufloo et al., 2021; Schäfer et al., 2021). Unprecedented global efforts have been undertaken to develop effective vaccines and prophylactic/therapeutic strategies to fight COVID-19 (Kelley, 2020). Immunotherapies based on SARS-CoV-2 neutralizing antibodies have been rapidly explored, and this led to the clinical use of several monoclonal antibodies (mAbs) alone or in bi-therapies (Corti et al., 2021). Highly potent human SARS-CoV-2 neutralizing mAbs isolated so far, including those tested or used in clinics, all target the RBD and can prevent infection and/or protect animals from severe disease in preclinical models (Andreano et al., 2021; Cao et al., 2020; Corti et al., 2021; Kreye et al., 2020; Noy-Porat et al., 2021; Rogers et al., 2020; Rosenfeld et al., 2021; Shi et al., 2020; Tortorici et al., 2020; Zost et al., 2020b). However, viral variants with spike mutations conferring resistance to antibody neutralization emerged during the pandemics and annihilated some of these therapies (Kumar et al., 2021; Planas et al., 2021b, 2021a; Radvak et al., 2021). The search for broadly neutralizing mAbs is being pursued. Novel antibodies active against all variants of concern (VOCs), including the currently prevalent omicron lineage, have been described (Cameroni et al., 2022; Gruell et al., 2022; Westendorf et al., 2022).

Here, we report on the detailed molecular and functional characterization of 102 human SARS-CoV-2 spike mAbs cloned from IgG and IgA memory B cells of ten convalescent COVID-19 individuals. These antibodies are encoded by a diverse set of immunoglobulin genes, recognize various conformational spike protein epitopes, and predominantly bind the S2 subunit. No anti-S2 mAbs were neutralizing but many harboured Fc-dependent effector functions. A third of the RBD-targeting antibodies potently neutralized SARS-CoV2 *in vitro*. The most potent, Cv2.1169 IgA and Cv2.3194 IgG, were fully active against VOCs Alpha, Beta, Gamma, and Delta, and still strongly blocked Omicron BA.1 and BA.2 infection *in vitro*. J-chain dimerization of Cv2.1169 IgA greatly improved its neutralization potency against BA.1 and BA.2. Cv2.1169 showed therapeutic efficacy in mouse and hamster SARS-CoV-2 infection models. Structural analyses by cryo-EM and X-ray crystallography revealed the mode of binding of Cv2.1169 and its contacts with the RBD at atomic level. Collectively, this study allowed gaining insights into fundamental aspects of the SARS-CoV-2-specific humoral response, and identified potent and broad neutralizers with prophylactic and therapeutic potential.

## Results

### Serological antibody profiling of COVID-19 convalescents

In convalescent COVID-19 individuals, serum antibody levels against the spike and RBD proteins have been correlated to SARS-CoV-2 seroneutralizing activities (Grzelak et al., 2020; Robbiani et al., 2020; Wang et al., 2021b). To select for convalescent donors with high seroneutralization for single B-cell antibody cloning, we first evaluated the IgG and IgA seroreactivity of convalescent individuals infected during the first epidemic wave (n=42 with bio-banked PBMC) to soluble recombinant Wuhan SARS-CoV-2 trimeric spike (tri-S) and RBD proteins by ELISA. Most of them had high titers of anti-tri-S IgGs, mainly IgG1, including cross-reacting antibodies against the Middle East respiratory syndrome-related coronavirus (MERS-CoV) tri-S protein (**Figures 1A, 1B, S1A** and **S1B**). High levels of serum anti-RBD IgGs were also detected (**Figures 1A, 1B, S1A** and **S1B**), and correlated with anti-tri-S antibody titers (**Figure S1C**). Although the SARS-CoV-2 seroreactivity of IgA antibodies was globally weaker than for IgGs, both were correlated (**Figures 1B, S1B** and **S1C**). Serum IgA and IgG antibodies from the ten donors with the highest anti-SARS-CoV-2 tri-S antibody titers (purple dots; **Figure 1A**) were purified, and showed strong ELISA binding to Wuhan nucleocapsid (N), tri-S, S1 and S2 subunits, and RBD, and also cross-reacted against recombinant spike proteins from other β-coronaviruses (SARS-CoV-1, MERS-CoV, HKU1 and OC43) as well as α-coronaviruses (229E and NL63) (**Figures 1C, S1D**, and **S1E**). The neutralizing activity of purified serum IgA and IgG antibodies against the Wuhan SARS-CoV-2 strain was then determined using an *in vitro* pseudoneutralization assay (**Figure 1D**). Fifty percent inhibitory concentrations (IC_50_) of purified IgA antibodies were in average lower as compared to IgGs (70.4 *vs* 115.6 µg/ml for IgAs and IgGs, respectively, p=0.068), ranging from 43 to 133 µg/ml for IgAs, and from 21 to 257 µg/ml for IgGs (**Figure 1D**). IC_50_ values for IgA but not IgG antibodies were negatively correlated to their respective binding levels to SARS-CoV-2 S1 and RBD proteins (**Figure S2A**).

**Figure 1.**
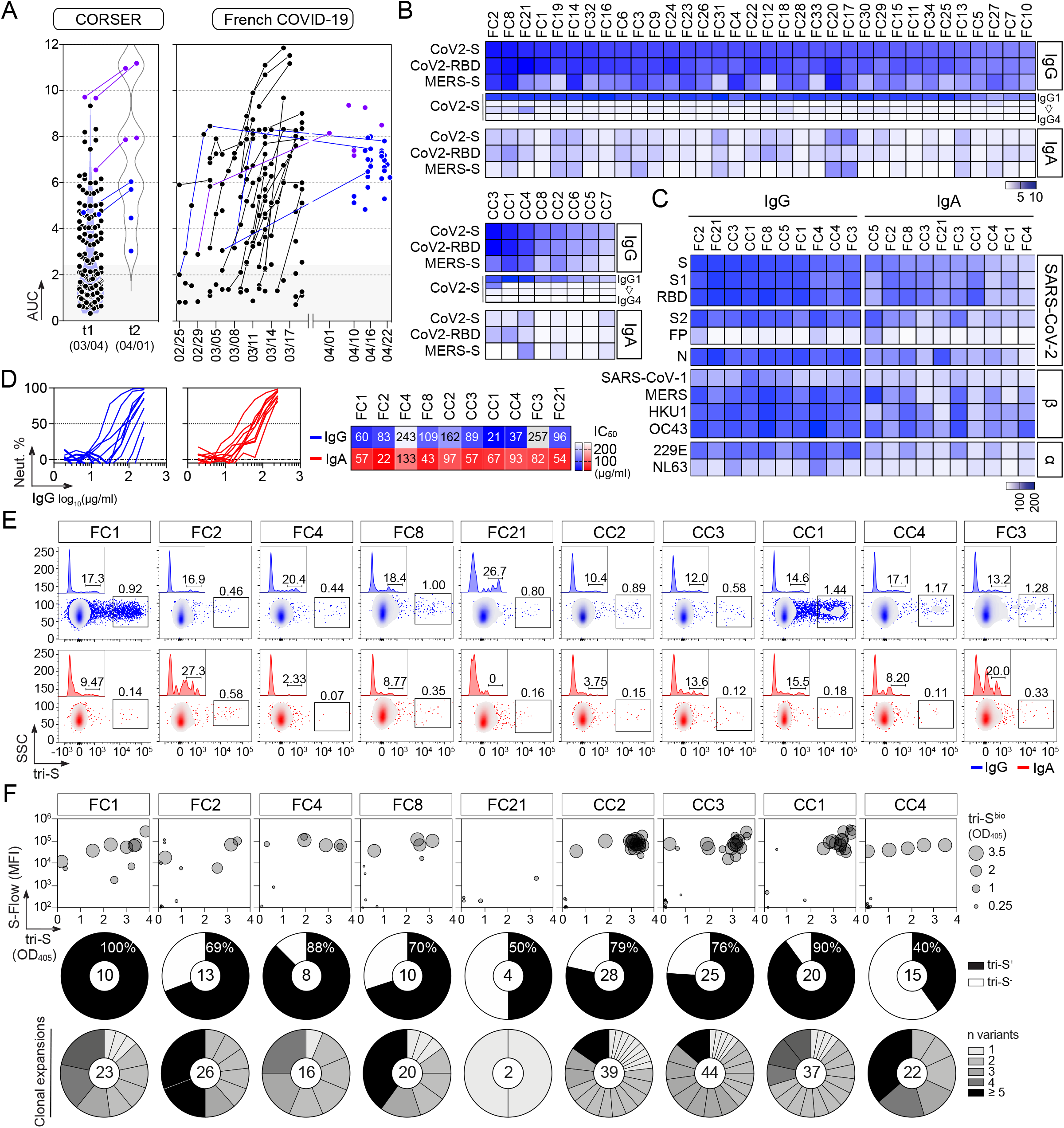
SARS-CoV-2 spike-specific memory antibodies cloned from convalescent COVID-19 individuals. **(A)** Dot plots showing the IgG antibody binding to SARS-CoV-2 tri-S as area under the curve (AUC) values determined by ELISA with serially-diluted sera from convalescent COVID-19 individuals in the CORSER (n=212; two timepoints t1 and t2) and French COVID-19 cohorts (n=159; with a follow-up overtime for some samples). Colored dots (blue and purple) show selected samples tested in (B). Purple dots indicate samples tested in (C). **(B)** Heatmap showing the IgG, IgG subclass and IgA seroreactivity of selected convalescent COVID-19 individuals from the CORSER (n=8) and French COVID-19 (n=34) cohorts against SARS-CoV-2 tri-S and RBD proteins as measured in Figure S1B. Samples were also tested against MERS tri-S to assay for cross-reactivity against another β-coronavirus. **(C)** Heatmap showing the antibody binding of serum IgG and IgA antibodies purified from selected convalescent donors against SARS-CoV-2 antigens and trimeric spike proteins from other coronaviruses (α, α-coronaviruses; β, β-coronaviruses) as measured in Figures S1D and S1E. RBD, receptor binding domain; FP, fusion peptide. **(D)** Graph showing the *in vitro* SARS-CoV-2 neutralizing activity of purified serum IgG and IgA antibodies from selected COVID-19 convalescents (left). Calculated IC_50_ values are presented in the heatmap on the right. **(E)** Flow-cytometric plots showing the SARS-CoV-2 S-binding IgG^+^ and IgA^+^ memory B cells in the blood from convalescent donors. Flow-cytometric histograms in the upper left-hand corner show the proportion of RBD^+^ cells among SARS-CoV-2 S-binding IgG^+^ and IgA^+^ memory B lymphocytes. **(F)** Bubble plots showing the reactivity of human IgG mAbs cloned from SARS-CoV-2 S-binding IgG^+^ and IgA^+^ memory B cells of convalescent donors against SARS-CoV-2 S protein as measured by S-Flow (Y axis), tri-S ELISA (X axis) and tri-S-capture ELISA (bubble size). For each donor, the pie chart shows the proportion of SARS-CoV-2 S-specific antibodies from cloned antibodies (top; total number indicated in the pie chart center) and the number (n) of variants in each SARS-CoV-2 S-specific B-cell clonal family. See also **Table S1** and **Figure S1**.

### Human SARS-CoV-2 spike-specific memory B-cell antibodies from COVID-19 convalescents

Next, peripheral blood IgA^+^ and IgG^+^ memory B cells from the selected convalescent individuals were stained with fluorescently-labeled RBD and tri-S, the latter being used as a bait to capture single SARS-CoV-2-reactive B cells by flow cytometric sorting (**Figure 1E**). From the 2870 SARS-CoV-2 tri-S^+^ IgA^+^/G^+^ memory B cells isolated, we produced a total of 133 unique human mAbs by recombinant expression cloning (Tiller et al., 2008), with most of them being part of B-cell clonal expansions (**Figure 1F**). ELISA and flow cytometry-based (S-Flow) binding analyses showed that 101 purified mAbs specifically bind to SARS-CoV-2 S protein (76% [40-100%]; **Figures 1F** and **S1F**). RBD-binding cells represented 11% and 17% of the tri-S^+^ IgA^+^ and IgG^+^ B cells, respectively (**Figure 2A**). Anti-RBD IgA titers were correlated with blood RBD^+^ IgA^+^ B-cell frequencies, and inversely correlated with neutralization IC_50_ values of IgAs (**Figure S2A**). Both total and SARS-CoV-2 tri-S-specific class-switched memory B cells showed a resting memory B-cell phenotype (RM, CD19^+^CD27^+^CD21^+^) (**Figures 2B**-**2D**). The frequency of circulating blood follicular helper T cell (cTfh) subsets was also determined. We found that cTfh2 (CD4^+^CXCR5^+^CCR6^-^CXCR3^-^), with a high proportion being activated (PD1^+/high^ and/or ICOS^+^), were predominant (**Figures 2E** and **2F**), and correlated with tri-S^+^ IgG^+^ RM B cells (r=0.83; p=0.0098) (**Figures 2G** and **S2B**), illustrating their capacity to promote class switching and affinity maturation of B cells as previously shown (Locci et al., 2013; Morita et al., 2011). Comparison of immunoglobulin gene features with IgG^+^ memory B cells from healthy controls (Prigent et al., 2016) revealed an increased usage in the SARS-CoV-2 spike-specific B-cell repertoire of rearranged V_H_3V_λ_3 (p=0.0047) and V_λ_3/J_λ_2 (p=0.0019), J_H_4 (p=0.0312), and J_κ_4 (p=0.0387) genes, as well as IgG1 subclass (p=0.0001) (**Figures 2H** and **S2**; **Table S1**). Anti-spike antibodies were also enriched in V_H_1-24/-69 and V_H_3-30/-33 genes (**Figure S2J**) as previously observed (Brouwer et al., 2020; Kreer et al., 2020; Vanshylla et al., 2022), and had reduced CDR_H_3 positive charges (p=0.0001) and somatic mutations in IgH (9.5 *vs* 19.2, p<0.0001) and Igλ (6.8 *vs* 12.4, p<0.0001) (**Figures 2H, 2I**, and **S2H**; **Table S1**). Certain antibody clones were shared among several of the COVID-19 convalescents (**Figure 2J**), demonstrating further the inter-individual convergence of antibody responses to SARS-CoV-2 as observed by others (Brouwer et al., 2020; Chen et al., 2021; Galson et al., 2020; Kreye et al., 2020; Nielsen et al., 2020; Robbiani et al., 2020; Vanshylla et al., 2022).

**Figure 2.**
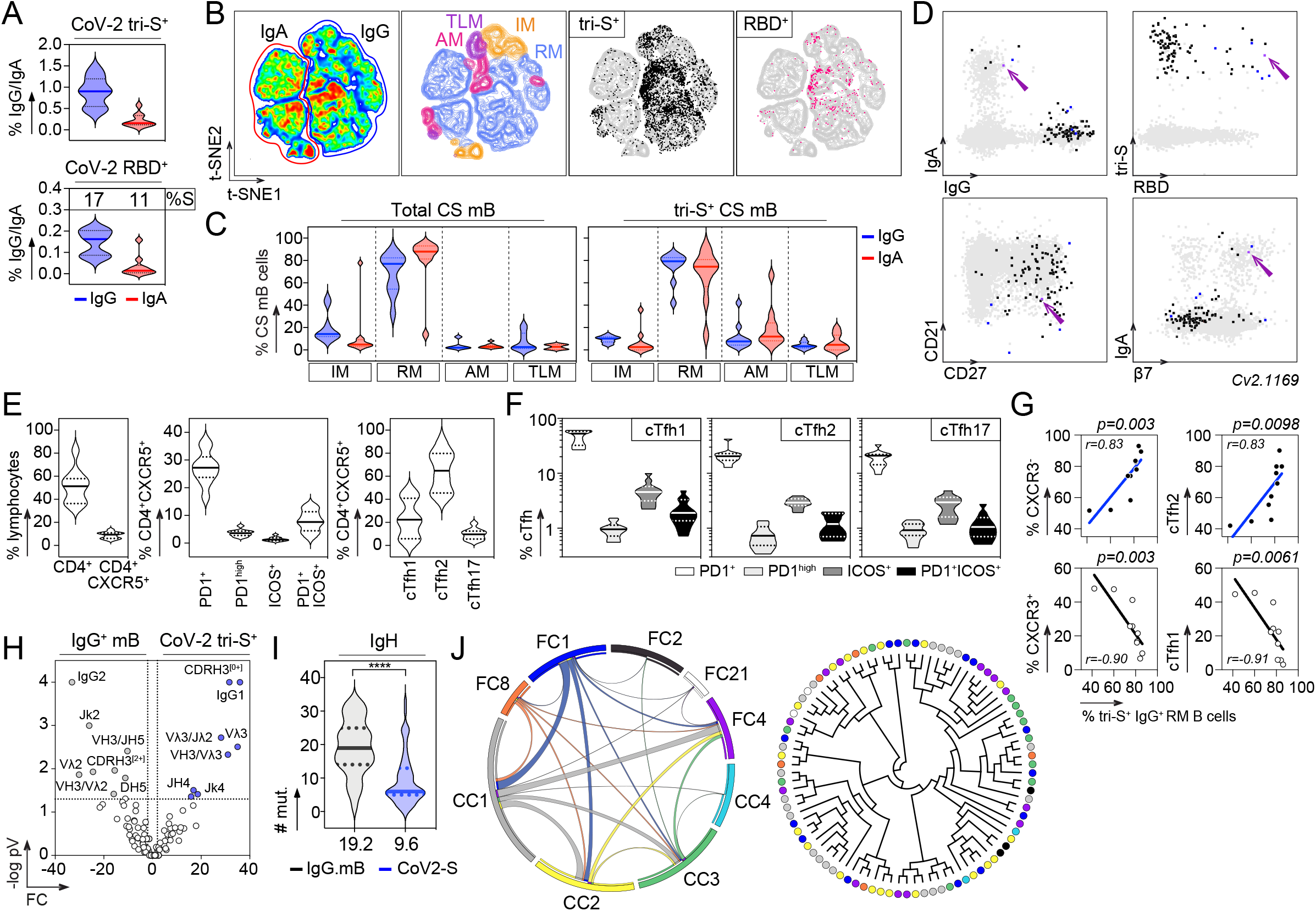
Immunophenotyping and antibody gene repertoire of SARS-CoV-2 spike-specific memory B cells. **(A)** Violin plots showing the percentage of SARS-CoV-2 tri-S^+^ cells among total IgG^+^ and IgA^+^ memory B cells (top) and of SARS-CoV-2 RBD^+^ cells among tri-S^+^ IgG^+^ and IgA^+^ memory B cells (bottom) in the blood of convalescent COVID-19 individuals (n=10). **(B)** Pseudocolor plots showing the t-SNE analysis of concatenated CD19^+^CD10^−^ B cells in convalescent COVID-19 individuals (n=10). Density maps presenting the staining intensity of CD27 and CD21 markers used to define memory B-cell subsets. IM (Intermediate memory, CD27^-^CD21^+^), RM (resting memory CD27^+^CD21^+^), AM (activated memory, CD27^+^CD21^-^), TLM (tissue-like memory CD27^-^CD21^-^). Black and pink dots indicate tri-S^+^ and RBD^+^ IgG^+^ and IgA^+^ B memory cells in the density map. **(C)** Violin plots showing the distribution of total and SARS-CoV-2 tri-S^+^ IgG^+^ and IgA^+^ memory B-cell subset frequencies as depicted in (B). CS mB, class-switched memory B cells in convalescent COVID-19 individuals (n=10). **(D)** Immunophenotyping flow cytometric plots showing the expression of B-cell surface markers on sorted SARS-CoV-2 tri-S-specific B cells (n=101, black, blue and purple dots). Blue dots indicate potent neutralizing antibodies while the purple dot is the ultra-potent neutralizer Cv2.1169 (purple arrow). **(E)** Violin plots showing the frequency of total CD4^+^, CD4^+^CXCR5^+^ lymphocytes and circulating follicular helper T cell (cTfh) subsets in the blood of convalescent COVID-19 individuals (n=10). **(F)** Violin plots comparing the frequency of PD1^+^, PD1^hi^, ICOS^+^ and ICOS^+^PD1^+^ cells among cTh1, cTfh2 and cTh17 subsets in the blood of convalescent COVID-19 individuals (n=10). **(G)** Correlation plots showing the frequency of SARS-CoV-2 tri-S^+^ IgG^+^ RM B cells *vs* CXCR3^+^ cTfh, CXCR3^-^ cTfh, cTfh1 and cTfh2. Spearman correlation coefficients with the corresponding p-values are indicated. **(H)** Volcano plot analysis comparing the immunoglobulin (Ig) gene repertoire of SARS-CoV-2 S-specific IgG^+^ / IgA^+^ B cells from convalescent donors and IgG^+^ memory B cells from healthy individuals (IgG.mB, unexposed to SARS-CoV-2). Grey and blue dots indicate statistically significant differences between both Ig gene repertoires. pV, p-value; FC, fold changes. **(I)** Violin plots comparing the number of mutations in V_H_ genes of SARS-CoV-2 S-specific and control IgG^+^ memory B cells from unexposed healthy individuals (n=72). The average number of mutations is indicated below. Numbers of mutations were compared across groups of antibodies using unpaired student t-test with Welch’s correction. **(J)** Circos plot (left) showing the clonal variants shared between distinct donors with the size of the links proportional to the number of clones sharing 75 % CDR_H_3 amino acid identity. Cladogram (right) showing the distribution of individual shared clones between donors. See also **Table S1** and **Figure S2**.

### Binding and antiviral properties of human anti-SARS-CoV-2 spike antibodies

Epitope mapping analyses showed that 59% of the anti-S mAbs (n=101) bind to the S2 subunit, 16% the RBD, 17% the NTD, 1% the S1 connecting domain (CD), and 7% to other regions of the SARS-CoV-2 spike (**Figures 3A** and **3B**; **Table S1**). Only one anti-S antibody (0.99% of the total) targeting S2 recognized the denatured tri-S protein by immunoblotting, but did not bind S-covering linear peptides (**Figures S3A-S3C**), indicating that most SARS-CoV-2-S memory antibodies target conformational epitopes. To determine whether anti-spike memory antibodies neutralize the Wuhan strain, we measured their inhibitory activity using three different *in vitro* functional assays: a competition ELISA measuring the blockage of soluble tri-S or RBD binding to ACE2 ectodomain, a pseudoneutralization assay and a neutralizing assay using live virus called S-Fuse (Sterlin et al., 2021) (**Figure 3C**). Overall, ∼ 15% of the anti-S mAbs showed inhibitory activities > 50% in the S-Fuse assay, many of which also neutralized pseudotyped SARS-CoV-2 virions and blocked tri-S-ACE2 interactions (**Figure 3C**; **Table S1**). Potent neutralizers targeted the RBD (**Table S1**), but only 50% of all anti-RBD mAbs blocked SARS-CoV-2 infection with IC_50_ values < 10 µg/ml (**Figures 3C, 3F** and **3G**; **Table S1**).

**Figure 3.**
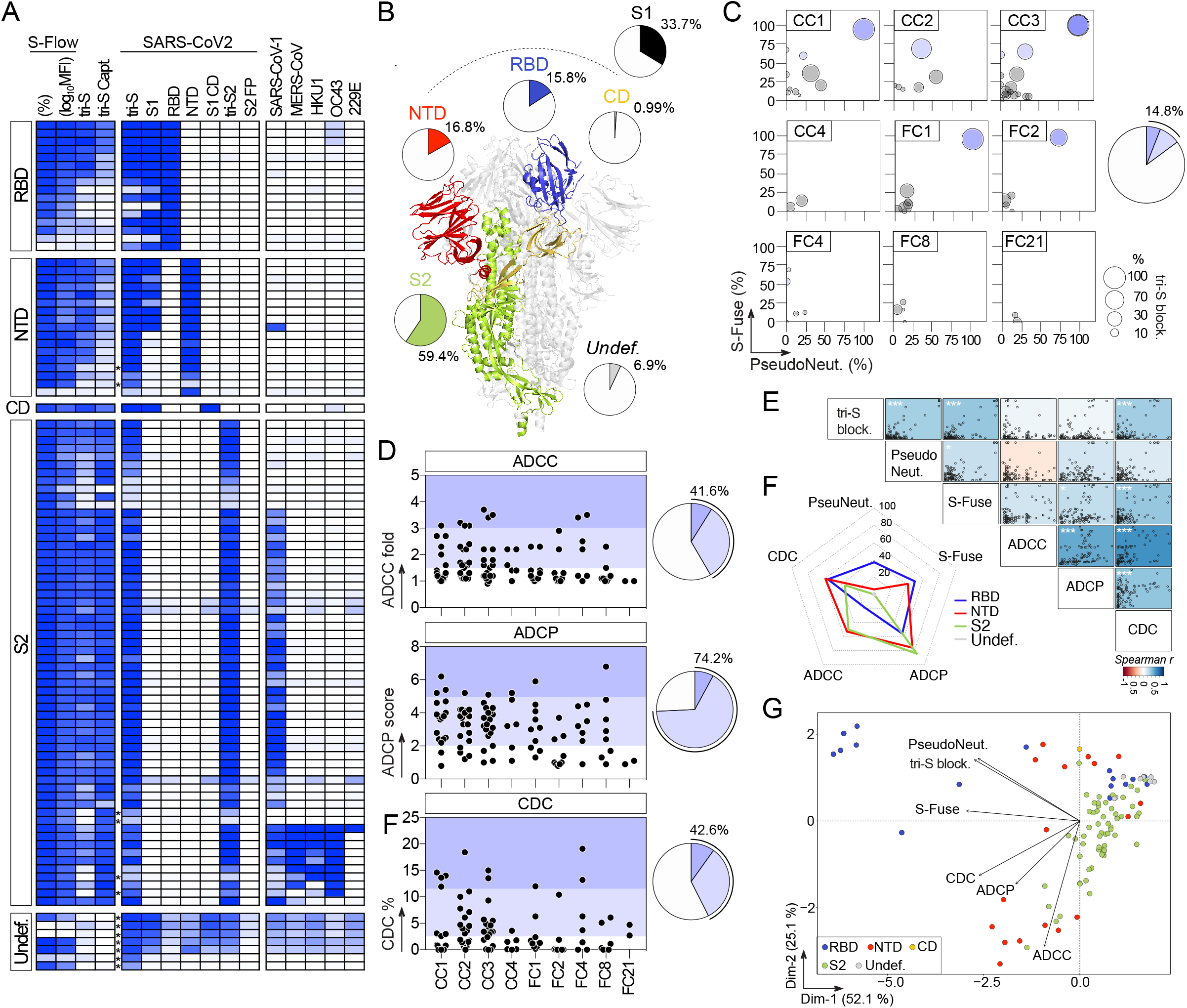
Reactivity and antiviral activities of SARS-CoV-2 S-specific memory B-cell antibodies. **(A)** Heatmap showing the reactivity of human anti-S mAbs (n=101) against SARS-CoV-2 antigens and trimeric spike proteins from other coronaviruses (α-coronaviruses: SARS-CoV-1, MERS-CoV, HKU1, and β-coronaviruses: OC43, 229E). RBD, receptor binding domain; NTD, N-terminal domain; CD, connecting domain; FP, fusion peptide. Asterisks indicate the antibodies tested at a higher IgG concentration. **(B)** Schematic diagram showing the distribution of specificities of anti-S antibodies on the highlighted regions of the SARS-CoV-2 spike as determined in (A) (ribbon representation of the PDB: 6VXX structure). **(C)** Bubble plots showing the neutralization activity of human SARS-CoV-2 S-specific antibodies (n=101) tested at a concentration of 10 µg/ml in the S-Fuse (Y axis), and pseudoneutralization (X axis, PseudoNeut.) assays against SARS-CoV-2. The bubble size corresponds to the blocking capacity of SARS-CoV-2 S-ACE2 interactions by the antibodies as measured by ELISA. Pie chart (right) show the distribution of non-active (white) *vs* neutralizing (shades of blue) antibodies according to neutralization % measured with the S-Fuse assay. **(D)** Dot plot showing the *in vitro* Fc-dependent effector activities of anti-S IgG antibodies (n=101). Pie charts (right) show for each measured effector function the distribution of non-active (white) *vs* active (shades of blue) antibodies. ADCC, antibody-dependent cell-mediated cytotoxicity; ADCP, antibody-dependent cellular phagocytosis; CDC, complement-dependent cytotoxicity. (D) Matrix showing the correlation analyses between neutralization activities and Fc-dependent effector functions measured for SARS-CoV-2 S-specific IgG antibodies. Spearman correlation coefficients (color coded) with their corresponding p values are shown. ***p<0.0001, *p<0.05. **(F)** Radar plots comparing the *in vitro* neutralizing and Fc-dependent effector activities of anti-S IgG antibodies according to their targeted spike domains. Percent of antibodies *per* specificity group mediating a given antiviral activity as determined in (D) is shown. **(G)** Principal component analysis 2D-plot showing the antiviral-related variables discriminating anti-S mAbs color-coded by specificities. The two dimensions account for 77.2% of the variability. The location of the variables is associated with the distribution of the antibodies. See also **Table S1**.

SARS-CoV-2 antibodies can be armed with Fc-dependent effector functions allowing the elimination of virions and infected cells (Dufloo et al., 2021), which can alter the course of infection *in vivo* (Schäfer et al., 2021; Winkler et al., 2021). We evaluated the *in vitro* capacity of anti-S mAbs to promote antibody dependent cellular cytotoxicity (ADCC), antibody dependent cellular phagocytosis (ADCP) and complement dependent cytotoxicity (CDC). On average, 41.6%, 74.2% and 42.6% of the IgG antibodies displayed ADCC, ADCP and CDC activity, respectively (**Figure 3D**). Effector activities of SARS-CoV-2 antibodies were globally correlated (**Figure 3E**). ADCC- and ADCP-inducing antibodies were directed principally against S2 (50% and 85%, respectively) and the NTD (53% and 76%, respectively) (**Figure 3F**; **Table S1**). Conversely, anti-RBD antibodies as a group were less efficient at performing ADCC, and to a lesser extent ADCP (**Figure 3F**; **Table S1**). SARS-CoV-2 mAbs with CDC potential targeted mainly the NTD (59% of anti-NTD) and the RBD (56% of anti-RBD) (**Figure 3F**; **Table S1**). Accordingly, CDC and tri-S-ACE2 blocking activities were correlated (**Figure 3E**). Principal-component analyses (PCA) showed that neutralizing and Fc-dependent effector functions segregated into two separate clusters in the PCA of antiviral functions, with 77% of the variance reached when combining the two first principal components (**Figure 3G**). The “neutralization” cluster included mainly anti-RBD antibodies, while the “effector” cluster comprised both NTD- and S2-specific IgGs (**Figure 3G**).

### Antibody features of potent SARS-CoV-2 neutralizers

In the collection of 101 anti-S mAbs, 5 potent SARS-CoV-2 neutralizing antibodies were identified (**Table S1**). They bound to the recombinant tri-S, S1 and RBD proteins with high affinity as measured by surface plasmon resonance (**Figure 4A**). They targeted similar or spatially-close epitopes on the RBD as shown by their cross-competition for ligand binding by ELISA (**Figures 4B** and **S3D**). They efficiently blocked the interaction of tri-S to the soluble ACE2 ectodomain (**Figure 4C**), suggesting that they recognize the receptor binding motif (RBM). IC_50_ values for SARS-CoV-2 neutralization, determined using the pseudoneutralization and S-Fuse assays, ranged from 3 to 37 ng/ml and from 0.95 to 11.5 ng/ml, respectively (**Figure 4D**). The most potent antibody, Cv2.1169, was encoded by V_H_1-58/D_H_2-15/J_H_3 and Vκ3-20/Jκ1 immunoglobulin gene rearrangements, and exhibited low levels of somatic mutation (3.1% V_H_ and 2.1% V_K_ at the amino acid level) (**Table S1**). The potential of the SARS-CoV-2 neutralizers to bind with low-affinity unrelated ligands (polyreactivity), and to cross-react with self-antigens was then evaluated in different complementary binding assays (**Figure S4**). None of the antibodies displayed self-reactivity, while only Cv2.3235 and Cv2.3194 showed polyreactivity (**Figure S4**). None of the potent neutralizers had ADCC potential, but showed moderate CDC and robust ADCP activities (**Figures S5A-S5C**). Remarkably, Cv2.1169 expressed as IgG1 antibod was one of the strongest ADCP-inducer among all the SARS-CoV-2 Spike mAbs (top 2%; **Figure S5C**; **Table S1**).

**Figure 4.**
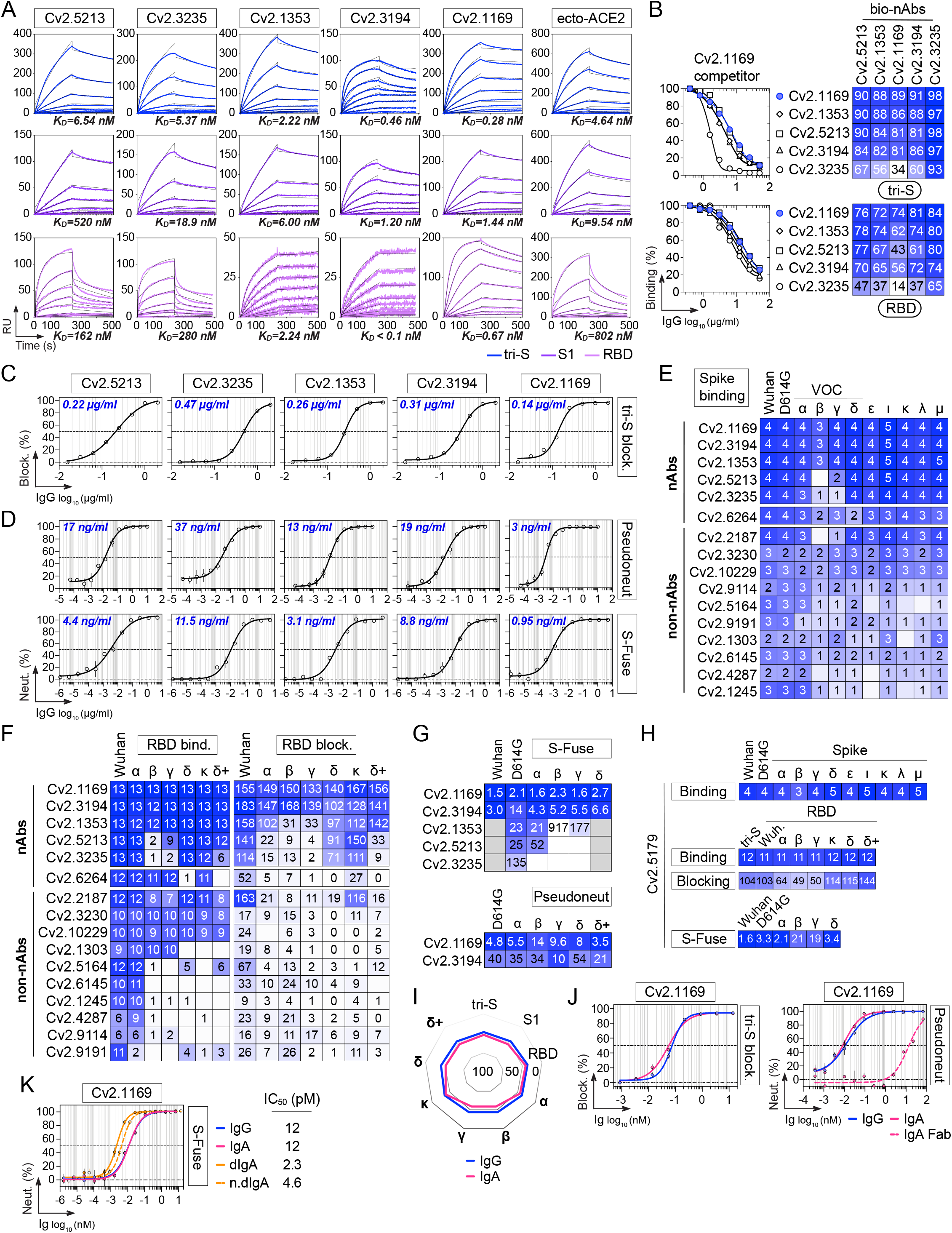
Binding and neutralizing properties of potent anti-RBD neutralizers. **(A)** SPR sensorgrams comparing the relative affinity of neutralizing anti-RBD IgG antibodies for the binding to SARS-CoV-2 S trimers (blue), S1 (purple) and RBD (pink) proteins. Calculated K_D_ values are indicated at the bottom. **(B)** Competition ELISA graphs (left) comparing the IgG binding to SARS-CoV-2 tri-S (top) and RBD (bottom) of selected biotinylated anti-RBD antibodies in presence of Cv2.1169 as potential competitor. Means ± SD of duplicate values are shown. Heatmaps (right) showing the competition of selected anti-RBD nAbs for tri-S and RBD binding as measured in Figure S4D. Dark blue indicates stronger inhibition; lighter colors indicate weaker competition, and white, no competition. **(C)** Competition ELISA graphs showing the binding of biotinylated SARS-CoV-2 tri-S protein to the immobilized soluble ACE2 ectodomain in presence of anti-RBD antibodies as competitors. Means ± SD of duplicate values are shown. **(D)** Graphs showing the neutralization curves of SARS-CoV-2 by selected anti-RBD IgG antibodies as determined with the pseudo-neutralization (top) and S-Fuse neutralization (bottom) assays. Error bars indicate the SD of assay triplicates. IC_50_ values are indicated in the top left-hand corner (in blue). **(E)** Heatmap comparing the binding of RBD-specific IgG antibodies to the cell-expressed spike proteins of SARS-CoV-2 and selected viral variants as measured by flow cytometry. Geometric means of duplicate log_10_ ΔMFI values are shown in each cell. **(F)** Heatmaps comparing the binding (left) and RBD-ACE2 blocking capacity (right) of RBD-specific IgG antibodies for the RBD proteins of SARS-CoV-2 and selected viral variants as measured in Figures S4E-4H. Darker blue colors indicate high binding or competition while light colors show moderate binding or competition (white = no binding or competition). AUC values are shown in each cell. **(G)** Heatmaps comparing the IC_50_ neutralizing values of the selected anti-RBD antibodies against SARS-CoV-2 and selected VOCs with the pseudo-neutralization (top) and S-Fuse neutralization (bottom) assays as measured in Figures S5A and S5B. **(H)** Heatmap showing binding to spike and RBD proteins (top), RBD-ACE2 blocking capacity (middle), and neutralizing activity (bottom) but for Cv2.5179 antibody as measured in Fig S6. **(I)** Radar plot comparing the binding of monomeric Cv2.1169 IgG and IgA antibodies to SARS-CoV-2 tri-S, S1 and RBD proteins, and to RBD from selected viral variants (in bold) as measured in Figure S4I. **(J)** Competition ELISA graphs (left) comparing the binding of biotinylated SARS-CoV-2 tri-S protein to the immobilized soluble ACE2 ectodomain in presence of Cv2.1169 IgG or IgA as a competitor. Means ± SD of duplicate values are shown. Graphs (right) comparing the SARS-CoV-2 neutralizing activity of Cv2.1169 IgG, IgA and IgA Fab as determined with the pseudo-neutralization assay. Error bars indicate the SD of duplicate values. **(K)** Graphs comparing the SARS-CoV-2 neutralizing activity of monomeric and dimeric IgA (dIgA) Cv2.1169 antibodies as determined with the S-Fuse neutralization assay. Error bars indicate the SD of triplicate values. n.dIgA, normalized values according to the number of binding sites. See also **Tables S1** and **S2**, and **Figures S3, S4, S5, S6** and **S7**.

### Neutralization spectrum of potent SARS-CoV-2 neutralizers

Several SARS-CoV-2 variants of concern (VOCs), *i*.*e*., Alpha (α, B.1.1.7), Beta (β, B.1.351), Gamma (γ, P.1) and Delta (δ, B.1.617.2), and variants of interest (VOIs) have emerged during the pandemics (WHO, 2022). We next evaluated the cross-reactive potential of the 16 anti-RBD antibodies against VOCs and VOIs. Binding analyses by flow cytometry showed that 3 out of the 5 potent neutralizers bound to cells expressing the spike proteins from VOCs (α, β, γ, δ) and VOIs (ε, ι, κ, λ, µ), while most non-neutralizing antibodies had narrowed cross-reactivity spectra (**Figure 4E**). Only neutralizers Cv2.1169, Cv2.3194 and Cv2.1353, as well as a third of the non-neutralizing antibodies, displayed unaltered ELISA binding to RBD proteins from the VOCs α, β, γ, δ and VOIs κ, δ^+^ (**Figures 4F, S3E** and **S3G**). Cv2.1169 and Cv2.3194 were the sole anti-RBD antibodies uniformly blocking the interaction of the ACE2 ectodomain with RBD proteins from the viral variants tested (**Figure 4F**; **Table S3**). Three potent neutralizers encoded by V_H_3-53/-66 immunoglobulin genes (Cv2.1353, Cv2.5213 and Cv2.3235), sensitive to the RBD mutations at positions 417 and 501 (Dejnirattisai et al., 2021a; Wibmer et al., 2021), lost binding and/or blocking activity against SARS-CoV-2 variants α, β, γ, δ (**Figures 4F** and **S3E-S3H**). Both S-Fuse and pseudo-neutralization assays showed that Cv2.1169 and Cv2.3194 neutralized SARS-CoV-2 VOCs α, β, γ, δ (**Figures 4G, S5D** and **S5E**). V_H_3-53 gene-expressing antibody Cv2.3194 efficiently bound and neutralized all the variants, most likely due to the usage of rearranged Vκ3-20/Jκ4 light chain genes as previously reported (Dejnirattisai et al., 2021a). Among these cross-neutralizers, Cv2.1169 was the most potent with IC_50_ values ranging from 1.5 to 2.7 ng/ml against Wuhan, D614G variant, α, β, γ, and δ strains in the S-Fuse assay, and from 3.5 to 14 ng/ml against D614G variant, α, β, γ, δ and δ^+^ strains in the pseudoneutralization assay (**Figures 4D, 4G, S5D** and **S5E**; **Table S3**). Cv2.1169 ranked among the strongest cross-neutralizers when compared to the parental versions of benchmarked antibodies used in clinics or in development (**Figures S6A-S6C**). In addition, we produced a Cv2.1169 IgG homolog (V_H_1-58/D_H_2/J_H_3 and Vκ3-20/Jκ1) from a different convalescent donor based on interindividual clonal convergence analyses (**Figures S7A** and **S7B**), Cv2.5179, which also exhibited a potent and broad SARS-CoV-2 neutralizing activity (**Figures 4H** and **S6C-S6E**).

Immunophenotyping of sorted B cells indicated that Cv2.1169 was originally produced by a Spike^+^RBD^+^ IgA^+^ B cell with an activated memory phenotype (CD27^+^CD21^-^), and a surface-expression of the mucosa-homing integrin β7 (**Figure 2D**). We thus also expressed Cv2.1169 as monomeric IgA antibody, which showed equivalent binding and neutralization activities compared to its IgG counterpart (**Figures 4I, 4J** and **S3I**). In contrast, purified J-chain containing IgA dimers demonstrated a higher neutralizing capacity against the Wuhan strain (**Figure 4K**), suggesting an enhanced neutralization by binding avidity effects as previously reported (Barnes et al., 2020a; Rujas et al., 2021). Accordingly, the neutralizing activity of Cv2.1169 IgA Fab against SARS-CoV-2 was strongly impaired as compared to the bivalent immunoglobulins (**Figure 4J**).

SARS-CoV-2 Omicron variant B.1.1.529 or BA.1 became dominant worldwide in January 2021, followed by Omicron BA.2 in March 2022 (WHO, 2022). Omicron BA.1 contains 15 RBD-amino acid substitutions, which conferred resistance to numerous potent anti-RBD neutralizers including those in clinical use (Cameroni et al., 2022; Cao et al., 2022a; Planas et al., 2022). BA.2 has 7 amino acids differing from BA.1 in the RBD, and is also less sensitive to antibody neutralization (Bruel et al., 2022). Cv2.1169 and Cv2.3194, but not the other anti-RBD antibodies, bound well to cell-expressed and soluble BA.1 spike proteins as well as to the BA.1 RBD (**Figure 5A**). Both antibodies blocked BA.1 tri-S binding to ACE2, although less efficiently than for the Wuhan viral spike (**Figure 5B**). Cv2.1169 and Cv2.3194 also had the highest binding and spike-ACE2-blocking capacity to BA.1 viral proteins by ELISA as compared to benchmarked antibodies (**Figures 5C** and **5D**). Cv2.1169 and Cv2.3194, but not Cv2.5179, neutralized BA.1 in the S-Fuse assay with IC_50_ of 253 ng/ml and 24.2 ng/ml, respectively (**Figure 5E**; **Table S3**). Thus, Cv2.1169 and Cv2.3194 presented, respectively, a 79- and 2.2-fold decreased neutralization efficacy on BA.1 omicron as compared to Delta (**Figure 5E**). In contrast, Cv2.1169 and Cv2.3194 showed a slightly stronger RBD-binding against Omicron BA.2 as compared to BA.1 (**Figure 5D**). Consistently, both antibodies blocked more efficiently the binding of the RBD BA.2 to soluble ACE2 (**Figure 5F**). Nonetheless, Cv2.1169 and Cv2.3194 showed comparable neutralizing activities against BA.1 and BA.2 in the S-Fuse assay (**Figure 5G**). As compared to their monomeric counterpart, dimeric Cv2.1169 IgA antibodies had enhanced RBD-binding and spike-ACE2 blocking activities to Omicron variants especially BA.1 (**Figure 5H** and **5I**). This translated into an increased neutralizing potency of Cv2.1169 IgA dimers against BA.1 and BA.2 by a 13- and 20-fold, respectively when normalized for the number of binding sites (**Figure 5J**).

**Figure 5.**
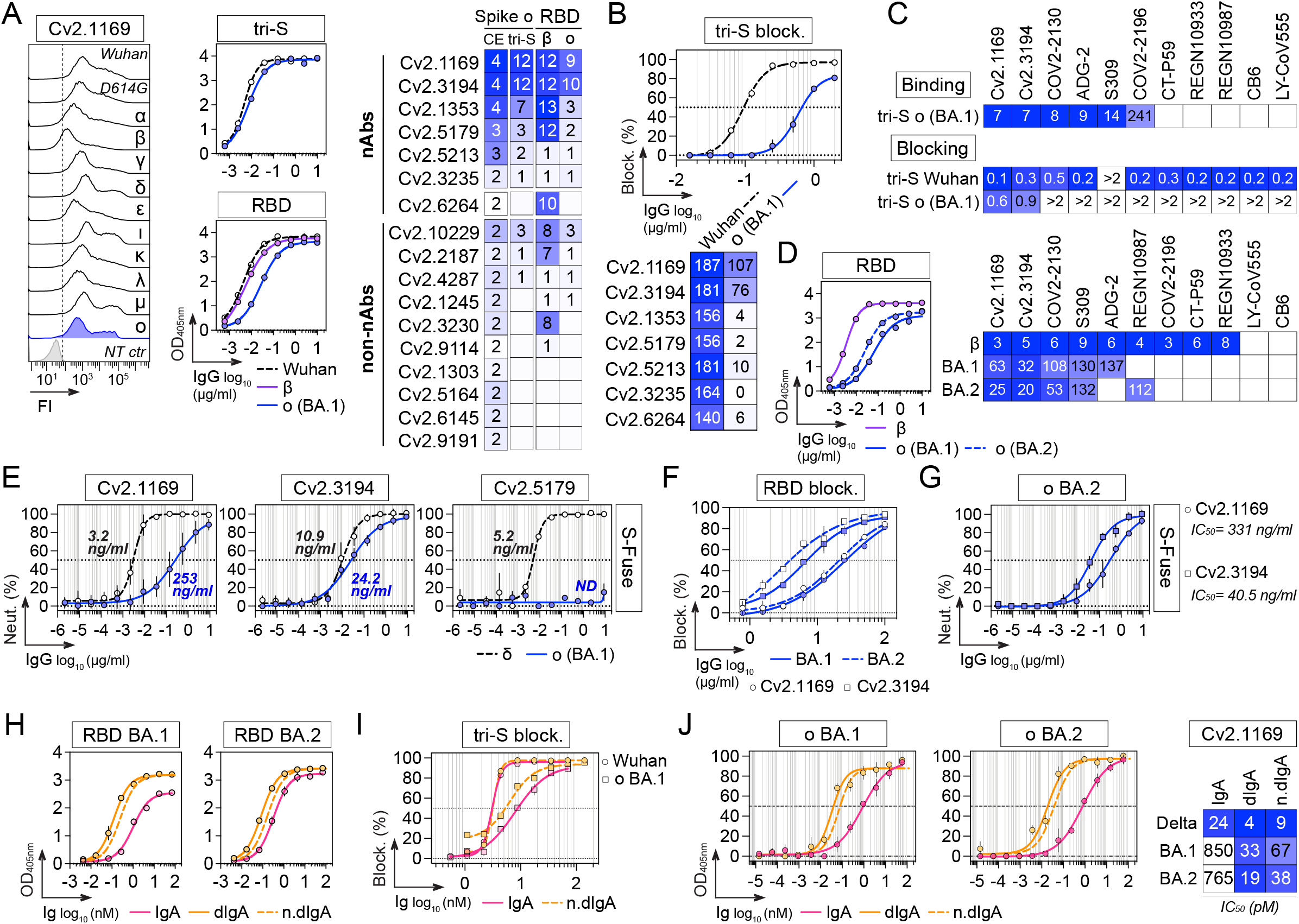
Activity of Cv2.1169 against SARS-CoV-2 Omicron variants. **(A)** Heatmap (right) comparing the binding of RBD-specific IgG antibodies to the cell-expressed (CE) and soluble (tri-S) Omicron (ο) SARS-CoV-2 spike proteins as measured by flow cytometry (mean log_10_ ΔMFI from duplicate values) and ELISA (mean AUC from duplicate values), respectively, as shown on the left for Cv2.1169. NT ctr, non-transfected cell control. The heatmap also presents the comparative antibody reactivity (AUC values) against β and ο RBD proteins. White indicates no binding. **(B)** Heatmap (bottom) comparing the RBD-ACE2 blocking capacity of neutralizing anti-RBD antibodies for the RBD proteins of SARS-CoV-2 and ο variant BA.1 as shown for Cv2.1169 (top). Darker blue colors indicate high competition while light colors show moderate competition (white = no binding or competition). Mean AUC from duplicate values are shown in each cell. **(C)** Heatmaps comparing the tri-S binding (top) and tri-S-ACE2 blocking capacity (bottom) of Cv2.1169 with benchmarked SARS-CoV-2 neutralizers RBD-specific IgG antibodies to the SARS-CoV-2 proteins of the ο variant BA.1. Darker blue colors indicate high binding or competition while light colors show moderate binding or competition (white = no binding or competition). Mean EC_50_ from duplicate values are shown in each cell. **(D)** Heatmap (right) comparing the binding of Cv2.1169 and Cv2.3194 with benchmarked SARS-CoV-2 neutralizers for the RBD proteins of the ο variant BA.1 and BA.2 as measured ELISA (means of duplicate AUC values) as shown on the left for Cv2.1169. Darker blue colors indicate high binding while light colors show moderate binding (white = no binding). Mean EC_50_ from duplicate values are shown in each cell. **(E)** Graphs showing the neutralization curves of SARS-CoV-2 δ and ο BA.1 by potent anti-RBD IgG antibodies as determined with the S-Fuse neutralization assay. Error bars indicate the SD of duplicate values from 2 (Cv2.5179) or 5 (Cv2.1169 and Cv2.3194) independent experiments. IC_50_ values are indicated (in blue for ο BA.1). ND, not determined. **(F)** Competition ELISA graphs showing the binding of biotinylated RBD proteins from SARS-CoV-2 o BA.1 and BA.2 variants to soluble ACE2 ectodomain in presence of Cv2.1169 and Cv2.3194 antibodies as competitors. Means ± SD of duplicate values are shown. **(G)** Same as in (F) but for Cv2.1169 and Cv2.3194 against BA.2. Error bars indicate the SD of duplicate values. **(H)** Graphs comparing the ELISA binding of monomeric and dimeric Cv2.1169 IgA antibodies to the RBD proteins of SARS-CoV-2 o BA.1 and BA.2 variants. Means ± SD of duplicate values are shown. n.dIgA, normalized values according to the number of binding sites. **(I)** Same as in (E) but for Wuhan and o BA.1 tri-S proteins with monomeric and dimeric Cv2.1169 IgA antibodies. Means ± SD of duplicate values are shown. n.dIgA, normalized values according to the number of binding sites. **(J)** Same as in (F) but for Cv2.1169 IgA monomers and J-chain dimers (dIgA) against BA.1 and BA.2. Error bars indicate the SD of duplicate values. Heatmap (right) presents the IC_50_ values calculated from the curves (left). n.dIgA, normalized values according to the number of binding sites.

### Structural characterization of the epitopes

To define the epitopes and neutralization mechanisms of the most potent mAbs, we co-crystallized the corresponding Fab in complex with the Wuhan RBD. The structures of the Cv2.3235 Fab/RBD and the Cv2.6264 Fab/RBD complexes were determined to 2.3 Å and 2.8 Å resolution, respectively (**Figure S9**; **Table S3**). The Cv2.1169 Fab/RBD binary complex did not crystallize, but the Cv2.1169 IgA Fab/CR3022 IgG1 Fab/RBD ternary complex produced crystals that allowed us to determine the X-ray structure to 2.9 Å. The electron density maps for the ternary complex were of poor quality and uninterpretable for the constant domain of Cv2.1169 Fab, indicating their intrinsic mobility. The Cv2.1169 variable domains and the paratope/epitope region were however well resolved (**Table S3**). The structure revealed that Cv2.1169 binds the RBM and straddles the RBD ridge leaning toward the face that is occluded in the “down” conformation of the RBD on a “closed” spike (**Figure 6A**). This binding mode is similar to other V_H_1-58/V_K_3-20-derived neutralizing antibodies (Dejnirattisai et al., 2021b; Starr et al., 2021; Tortorici et al., 2020; Wang et al., 2021a), as shown in the superposition of the RBD complexed with A23-58.1, COVOX-253 and S2E12 mAbs (**Figure S8**). Superposing the structures of the RBD/Cv2.1169 and RBD/ACE2 complexes showed extensive clashes between the antibody and the receptor (**Figure 6B**), providing the structural basis for its neutralization mechanism, and agreeing with its RBD-ACE2 blocking capacity (**Figures 4C, 4F, S3F** and **S3H**). Cv2.1169, Cv2.3235 and Cv2.6264 bound differently to the RBD, with Cv2.1169 having the lowest total buried surface area (BSA) (∼1400 Å^2^, ∼2620 Å^2^ and ∼1610 Å^2^, for Cv2.1169, Cv2.3235 and Cv2.6264, respectively) (**Table S4**), despite being the only mAb that contacts the RBD with all its CDRs. Cv2.1169 also has the highest heavy chain contribution to the interaction surface (∼80% of the paratope’s BSA), mainly through the CDR_H_3 (**Table S4**). The Cv2.1169 CDR_H_3 (14 amino acid length by Kabat definition) bends at P99 and at F110, delimiting a tongue-like loop that is stabilized by a disulfide bond between C101^CDRH3^ and C106^CDRH3^ (**Figure 6C**). This particular shape allows residues between G103 and F110, which are on one side of the CDR_H_3 tongue, to recognize the RBD tip and to form hydrogen bonds through their main-chain atoms (**Figure 6C**; **Table S5**). The interface is further stabilized by polar interactions between the side chains of D108 in the CDR_H_3 and Y33 in the CDR_L_1 (**Figure 6C**; **Table S4**).

**Figure 6.**
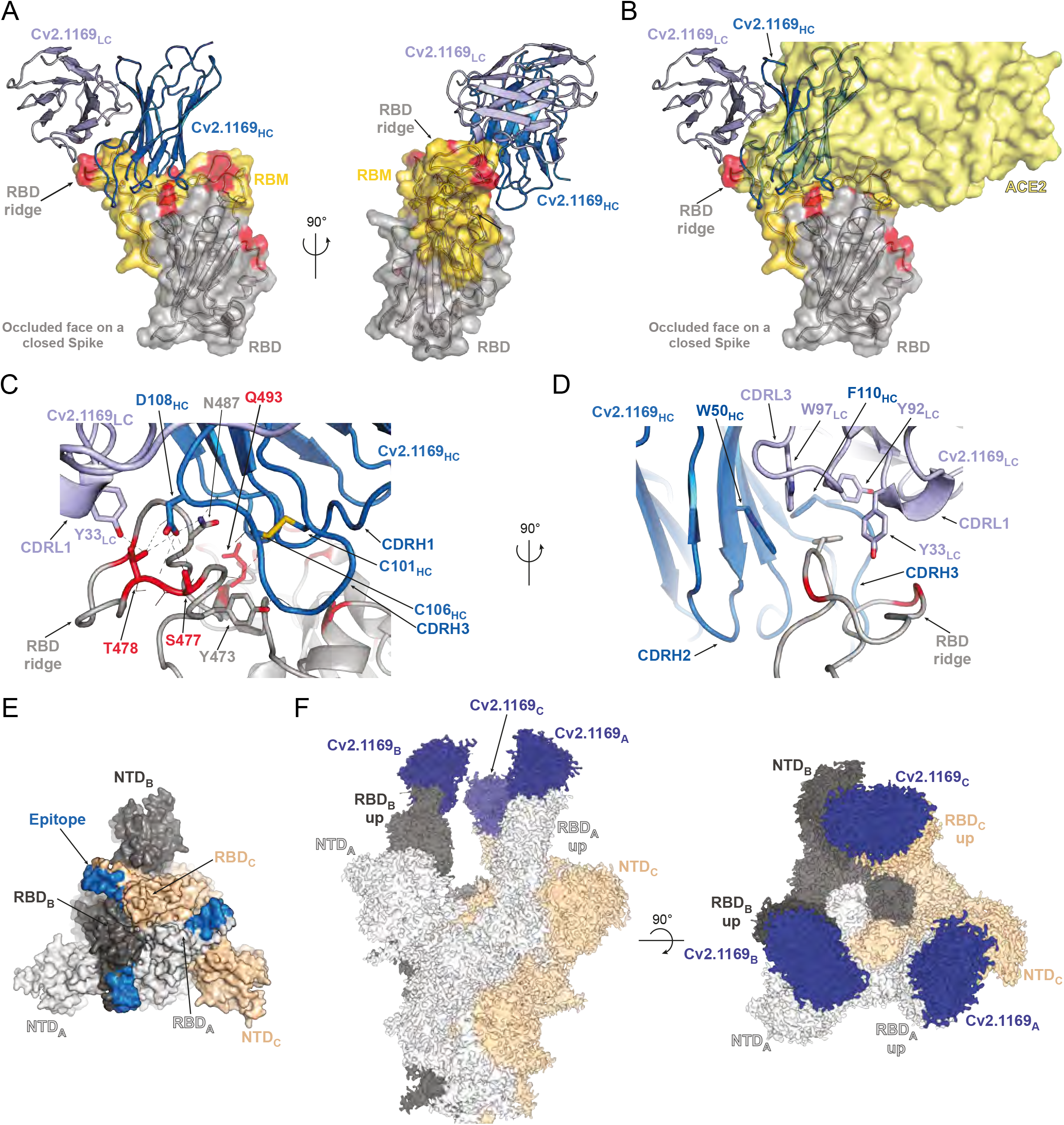
Structural analyses of the Cv2.1169 epitope. **(A)** Crystal structure of the complex formed by the Receptor Binding Domain (RBD) and Cv2.1169. The RBD is represented in cartoon with a transparent surface, highlighting the Receptor Binding Motif (RBM, yellow) and residues that are mutated in the Variants of Concern (VOCs, red). The constant domain from Cv2.1169 could not be built on the residual electron density and the variable domains are indicated in different shades of blue (IgH, dark blue; IgL, light blue). **(B)** Superposition of the RBD-Cv2.1169 and RBD-ACE2 (PDB: 6M0J) structures, showing the receptor on surface representation (light yellow) and its clashes with the antibody. **(C)** Close-up at the RBD-Cv2.1169 interface. For clarity, only the side chains from residues forming hydrogen bonds (dashed lines) are shown as sticks. Residues mutated in the VOCs are in red and the CDR_H_3 disulfide bond is indicated with yellow sticks. **(D)** Details of the hydrophobic residues that anchor F486 at the interface between the light and heavy chains of Cv2.1169. **(E)** Identification of the Cv2.1169 epitope (blue) on the structure of a closed spike (PDB: 6VXX). The different protomers are identified with a subscript letter and colored in light grey (protomer A), dark grey (protomer B) and wheat (protomer C). **(F)** Cryo-EM map from the trimeric spike ectodomain in complex with Cv2.1169. See also **Tables S3**-**S5**, and **Figures S8, S9** and **S10**.

Cv2.1169 epitope comprises the RBD segments 417-421, 455-458, 473-478 and 484-493 (**Figures 6A** and **6C**; **Table S5**). Apart from T478, all the mutated RBD residues present in the SARS-CoV-2 VOCs prior to Omicron are at the rim of the contact area (K417, E484) or outside (L452, N501) (**Figures 6A** and **6C**). Conversely, Cv2.3235 interacts with several residues mutated in several VOCs, *e*.*g*., K417 and N501 (**Figures S9A** and **S9C**), explaining its reduced capacity to bind and to neutralize α, β, γ and δ^+^ variants (**Figures 4E, 4F** and **S5A**). The RBD residue T478 forms hydrogen bonds with Cv2.1169 heavy and light chains, and is mutated in the δ and δ^+^ variants (T478K) (**Figure 6C**; **Table S5**). Despite this substitution, Cv2.1169 is still able to efficiently bind and neutralize both variants (**Figures 4E, 4F, S5A** and **S5B**). This indicates that the interface integrity does not depend on the hydrogen bonds formed with the T478 side chain and that there is enough space for the lysine residue to adopt a rotamer with reduced clashes with the antibody. Unlike the Cv2.6264 antibody, which also straddles the RBD ridge but lost reactivity against the δ and δ^+^ variants (**Figures 4E, S9B** and **S9D**), Cv2.1169 buries the RBD F486 within a hydrophobic cavity. This pocket is formed by aromatic residues of the FWR_H_2 (W50), the CDR_H_3 (F110), the CDR_L_1 (Y33) and the CDR_L_3 (Y92 and W97) (**Figure 6D**), and mimics the environment encountered when interacting with ACE2 (Lan et al., 2020). Thus, the F486 residue likely acts as an anchor for Cv2.1169, strengthening its interaction with the RBM allowing to tolerate the T478K mutation in the δ and δ^+^ variants. Four of the Cv2.1169-RBD contacting residues are mutated in BA.1 and BA.2 variants, including the substitution K417N already present in β and γ, and T478K in δ, as well as two Omicron-specific mutations S477N and Q493R (**Tables S5**). Although all of them are at the periphery of Cv2.1169 binding site (**Figures 6A-6C**), their combination explains the decreased binding and neutralization of SARS-CoV-2 BA.1 and BA.2 compared to the other VOCs (**Figure 5**).

As afore-mentioned, Cv2.1169 leans towards the RBD’s occluded face, making the epitope inaccessible on the ‘down’ conformation (**Figure 6E**), which implies that the antibody binds only to the RBD in its ‘up’ conformation. This was confirmed by the 2.8Å cryo-EM reconstruction of the SARS-CoV-2 S_6P protein trimer in complex with Cv2.1169 IgA Fab (See **Figure S10** for the cryo-EM processing strategy). The map showed that the spike is in the open form with each protomer bound by a Cv2.1169 Fab (**Figure 6F**). Considering that Cv2.1169 blocked SARS-CoV-2 tri-S binding to soluble ACE2 receptor, and that its binding site is only accessible in the up-RBD conformation, our data suggest that the antibody belongs to the class 1 category (or Ia) (Barnes et al., 2020b), with an epitope in the RBD-B group (Yuan et al., 2021). Accordingly, Cv2.1169 cross-competed for binding to spike and RBD proteins with class 1 benchmarked SARS-CoV-2 neutralizers (CT-P59, COV2-2196, REGN10933, and CB6), but also moderately with class 2 antibody LY-CoV555 (**Figure S6D**).

### *In vivo* therapeutic activity of Cv2.1169 against SARS-CoV-2 infection

We evaluated the *in vivo* therapeutic potential of neutralizing antibody Cv2.1169 using first the K18-hACE2 transgenic mouse model for SARS-CoV-2 (Wuhan strain) infection. Mice intranasally infected with 10^4^ PFU of SARS-CoV-2 were treated 6 h later with a single intraperitoneal (i.p.) injection of Cv2.1169 IgG antibody (0.25 mg, ∼10 mg/kg and 0.5 mg, ∼20 mg/kg) or control IgG antibody (0.5 mg, ∼20 mg/kg) (**Figure 7A**). Infected mice from the control group lost up to 25% of their body weight within the first 6 days post-infection (dpi) before reaching humane endpoints at 7-8 dpi (**Figure 7A**). In contrast, all animals treated with Cv2.1169 IgG survived and recovered their initial body weight after experiencing a transient loss during the first week (**Figure 7A**). Even when infected with a higher viral inoculum (10^5^ PFU SARS-CoV-2), and treated 22 h post-infection with Cv2.1169 IgG (∼ 40 mg/kg *i*.*p. plus i*.*n*.), half of the mice survived compared to those in the control group (p=0.029) (**Figure 7B**). Next, to evaluate the *in vivo* efficacy of Cv2.1169 IgA antibodies, a single low dose of either Cv2.1169 IgA or IgG antibodies (0.125 mg *i*.*p*., ∼ 5 mg/kg) was administered to SARS-CoV-2-infected mice (10^4^ PFU challenge dose). Despite a significant and comparable reduction of viral loads in the oral swabs of Cv2.1169 IgA- and IgG-treated mice compared to control animals at 4 dpi (2.6×10^4^ eqPFU/ml *vs* 5.7×10^3^ eqPFU/ml for Cv2.1169 IgA [p=0.008], and 4.7×10^3^ eqPFU/ml for Cv2.1169 IgG [p=0.029]) (**Figure S11A**), all mice treated with the SARS-CoV-2 IgAs were euthanized at 7-8 dpi, whereas 75% of the Cv2-1169 IgG-treated mice lost weight and developed symptoms but recovered their initial body weight after 2 weeks (**Figure 7C**). This can be explained by the rapid decay of circulating human IgA as compared to IgG antibodies in mice (**Figure S11C**).

**Figure 7.**
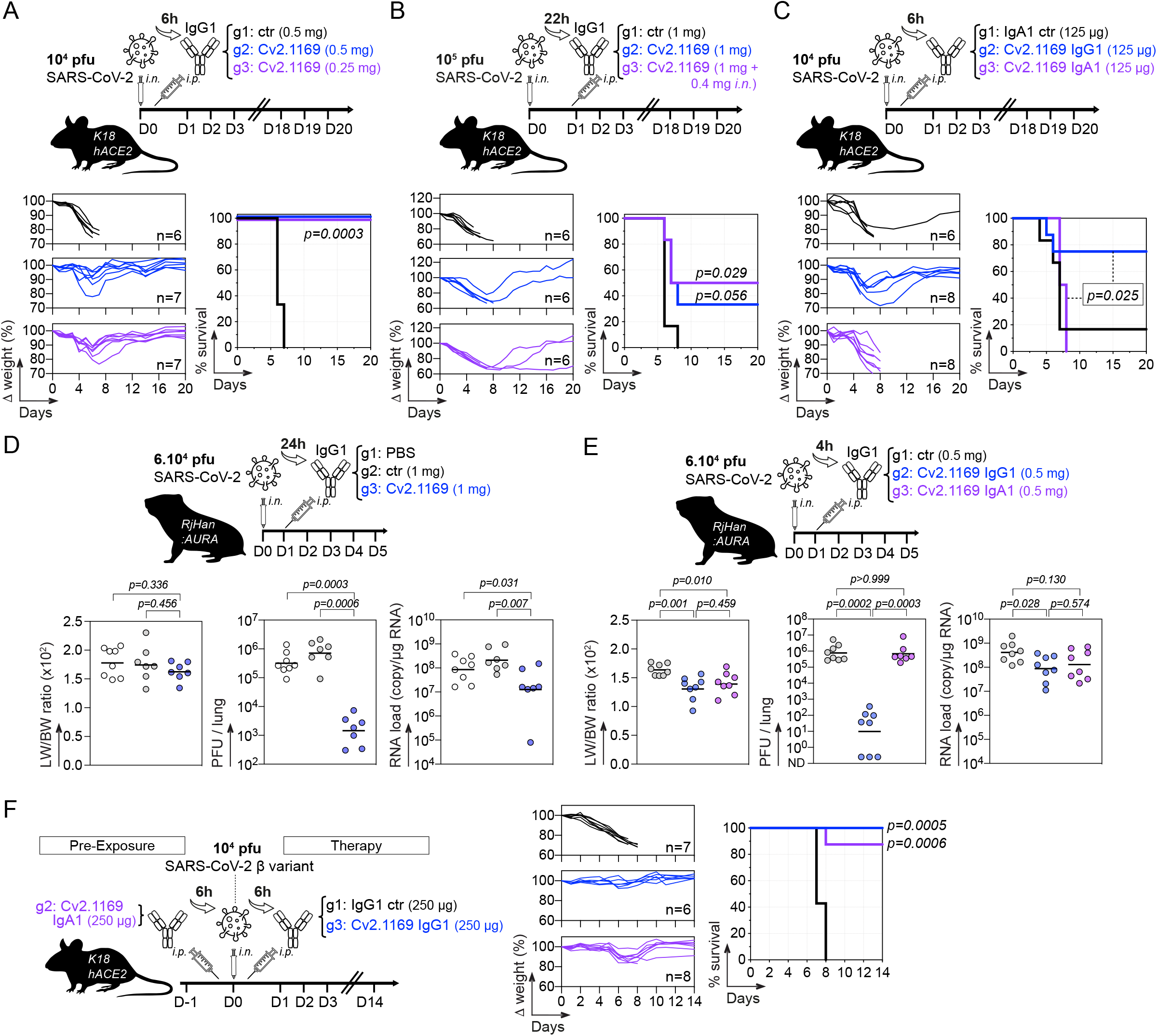
*In vivo* therapeutic activity of potent SARS-CoV-2 neutralizer Cv2.1169. **(A)** Schematic diagram showing the experimental design of Cv2.1169 antibody therapy in SARS-CoV-2-infected K18-hACE2 mice (top). Animals were infected intranasally (i.n.) with 10^4^ plaque forming units (PFU) of SARS-CoV-2 and received 6 h later an intraperitoneal (i.p.) injection of Cv2.1169 or isotypic control IgG antibody at ∼ 10 mg/kg (0.25 mg) and ∼ 20 mg/kg (0.5 mg). Graphs showing the evolution of initial body weight (% Δ weight, bottom left) and survival rate (bottom right) in animal groups. Groups of mice were compared in the Kaplan-Meier analysis using Log-rank Mantel-Cox test. **(B)** Same as in (A) but with K18-hACE2 mice infected with 10^5^ PFU and treated 22 h later with 1 mg i.p. of Cv2.1169 IgG antibody (∼ 40 mg/kg). **(C)** Same as in (A) but with infected mice treated with Cv2.1169 IgG and IgA antibodies at ∼ 5 mg/kg (0.125 mg). **(D)** Schematic diagram shows the experimental design of Cv2.1169 antibody therapy in SARS-CoV-2-infected golden Syrian hamsters (top). Animals were infected intranasally (i.n.) with 6×10^4^ plaque forming units (PFU) of SARS-CoV-2 and received 24 h later an intraperitoneal (i.p.) injection of PBS, Cv2.1169 or isotypic control IgG antibody at ∼ 10 mg/kg (1 mg). Dot plots showing the lung weight / body weight ratio (LW/BW) x 100 (left), infectivity (center) and RNA load (right) measured in animal groups at 5 dpi. Groups of hamsters were compared using two-tailed Mann-Whitney test. **(E)** Same as in (D) but with infected animals treated 4 h later with Cv2.1169 IgG and IgA antibodies at ∼ 5 mg/kg (0.5 mg). **(F)** Same as in (A) but with K18-hACE2 mice infected with 10^4^ PFU of the SARS-CoV-2 variant β (B.1.351), and either pre-treated 6h before infection with ∼ 5 mg/kg (0.5 mg) of Cv2.1169 IgA or treated 6h post-infection with ∼ 5 mg/kg (0.5 mg) of Cv2.1169 IgG or isotype control (ctr). See also **Figures S11**.

SARS-CoV-2-related pathogenesis in infected Golden Syrian hamsters resemble mild-to-moderate COVID-19 disease in humans (Imai et al., 2020; Sia et al., 2020). To further evaluate the *in vivo* efficacy of Cv2.1169 IgG neutralizer, hamsters infected *i*.*n*. with 6.10^4^ PFU of SARS-CoV-2 were treated 24 h later with a single injection of Cv2.1169 IgG or control antibodies (1 mg *i*.*p*., ∼10 mg/kg) (**Figure 7D**). Lung weight to body weight (LW/BW) ratio, intra-lung viral infectivity and RNA load were measured at 5 dpi. Both pulmonary viral infectivity and RNA levels in hamsters treated with Cv2.1169 were significantly reduced compared to control animals (2.44×10^3^ *vs* 10×10^5^ PFU/lung, p=0.0005 and 4.3×10^7^ *vs* 3.4×10^8^ copies/µg RNA, p=0.013, respectively) (**Figure 7D**). We next compared the *in vivo* activity of Cv2.1169 IgG and IgA antibodies at a dose ∼5 mg/kg in hamsters 4h post-infection. IgA- and IgG-treated hamsters showed a reduction in LW/BW ratio compared to control animals (1.64 *vs* 1.4 for IgA [p=0.03] and 1.32 for IgG [p=0.004]) (**Figure 7E**). As expected from the rapid disappearance of circulating human IgA antibodies in treated animals (**Figure S11E**), the intra-lung viral infectivity and RNA loads were comparable between SARS-CoV-2 neutralizing IgA-treated and control hamsters (**Figure 7D**). In contrast, the administration of Cv2.1169 IgG antibodies reduced both SARS-CoV-2 infectivity and RNA levels in the lungs of treated hamsters (1.39×10^6^ *vs* 80 PFU/lung, p=0.0002; 6.14×10^8^ *vs* 1.51×10^8^ copies/µg RNA, p=0.028) (**Figure 7D**). Cv2.1169 IgA and IgG-treated animals showed similar endogenous anti-spike IgG titers, which were reduced as compared to the control group (p<0.0001 and p=0.0003, respectively), suggesting potential early antiviral effects of Cv2.1169 IgA antibodies against SARS-CoV-2 infection (**Figure S11F**).

To determine whether Cv2.1169 is active *in vivo* against infection with SARS-CoV-2 VOCs, we tested the prophylactic activity of Cv2.1169 IgA antibodies and the therapeutic activity of Cv2.1169 IgG antibodies against SARS-CoV-2 VOC Beta in K18-hACE2 transgenic mice. A single administration of Cv2.1169 IgA antibodies at ∼10 mg/kg (0.25 mg *i*.*p*.) 6h prior to infection with 10^4^ PFU of SARS-CoV-2 Beta (β) protected 87.5% of the animals from death (**Figure 7F**). Despite the fact that human SARS-CoV-2 IgA antibodies did not persist in the mouse circulation (**Figure S11C**), Cv2.1169 IgA-treated mice also recovered their initial body weight during the follow-up (**Figure 7F**). Likewise, treating once SARS-CoV-2 Beta-infected mice with Cv2.1169 IgG antibodies (0.25 mg *i*.*p*., ∼10 mg/kg) 6h post-infection led to 100% survival, while all animals receiving the control antibodies were euthanized at 7-8 dpi (**Figure 7F**). Of note, human Cv2.1169 IgG antibodies were still detectable in mouse sera at the end of the follow-up (**Figures S11B** and **S11C**). In addition, mice pre-treated with Cv2.1169 IgAs developed higher anti-spike IgG antibody titers as compared to those treated with Cv2.1169 IgG antibodies, suggesting a weaker viral control in the former group (**Figure S11D**).

## Discussion

SARS-CoV-2 infection triggers the production of high-affinity IgGs and IgAs to the viral spike, including neutralizing antibodies, released in mucosal secretions and circulating in the blood (Smith et al., 2021; Sterlin et al., 2021). Class-switched IgG and IgA memory B cells are also elicited during COVID-19, persist for months post-infection, and can continue to mature and expand upon antigenic challenges (Gaebler et al., 2021; Sokal et al., 2021; Wang et al., 2021c). In line with previous reports (Sterlin et al., 2021; Zhou et al., 2021b), we found that serum IgA antibodies from COVID-19 convalescents neutralize SARS-CoV-2, often more efficiently than their IgG counterparts despite their lower representativeness in the blood. IgA neutralizing titers were correlated to anti-S1/-RBD antibody levels and spike^+^ memory IgA B-cell frequencies, suggesting coordinated serological and cellular humoral responses in these individuals as previously reported (Juno et al., 2020). We also document an association between spike-reactive resting memory IgG B cells and Th2-like cTfh cells, which likely encompass spike-specific cTfh2 cells (Juno et al., 2020). In this study, we characterized SARS-CoV-2 spike-specific IgG^+^ and IgA^+^ memory B-cell antibodies from COVID-19 convalescent individuals with high seroneutralization titers. Surprisingly, only a minority (∼7%) of the antibodies - all targeting the RBD - efficiently neutralized SARS-CoV-2 *in vitro*. Other less potent anti-RBD and several anti-NTD antibodies neutralizing SARS-CoV-2 were also isolated as previously reported (Andreano et al., 2021; Brouwer et al., 2020; Chi et al., 2020; Liu et al., 2020; Robbiani et al., 2020; Wec et al., 2020; Zost et al., 2020b).

Besides neutralization, SARS-CoV-2 IgGs can exert antiviral effector functions dependent or not on their binding to FcγR (*i*.*e*., ADCC/ADCP and CDC, respectively), playing a role in the therapeutic protection against SARS-CoV-2 infection *in vivo* (Schäfer et al., 2021; Winkler et al., 2021). Here, we found that despite lacking high neutralization potential, anti-S2 and anti-NTD IgGs harbor strong Fc-dependent effector functions less frequently observed with anti-RBD antibodies. This tendency suggests a dichotomy of antiviral functions based on epitope specificity, with antibodies to the spike head (RBD) being neutralizers and those to the stalk (S2) being effectors, while anti-NTD displayed mixed activities. Of note, one neutralizing antibody termed S2P6 targeting the S2 stem helix peptide also mediates a strong ADCC activity (Pinto et al., 2021).

Among the 102 SARS-CoV-2 antibodies described in this study, Cv2.1169 and Cv2.3194 were the sole potent neutralizers with a sustained activity against all SARS-CoV-2 variants, including Omicron BA.1 and BA.2 subtypes. Comparably to typical class 1 anti-RBD antibodies, Cv2.3194 uses V_H_3-53 variable genes and displays a short CDR_H_3 (Yuan et al., 2020, 2021), but differs from the others by its resistance to escape mutations in the VOCs. Indeed, V_H_3-53-encoded anti-RBD antibodies usually lose their capacity to neutralize SARS-CoV-2 viruses with mutations in position K417 and N501 including the VOCs α, β, γ, and δ (Yuan et al., 2021; Zhou et al., 2021a). A rare mutation in the CDRκ1 of Vκ3-20-expressing class 1 anti-RBD antibodies (P30S) has been proposed to rescue VOC neutralization (Dejnirattisai et al., 2021a), but is absent in Cv2.3194. As the Cv2.3194 Fab/ RBD complex did not crystallize, the molecular basis for its unaltered potent cross-neutralizing capacity against all VOCs remain to be solved. The other potent SARS-CoV-2 cross-neutralizing antibody, Cv2.1169, is a class 1 neutralizer binding to RBD with a modest total buried surface area. Except for Omicron BA.1 and BA.2, all mutated RBD residues in the SARS-CoV-2 VOCs had a negligeable impact on the SARS-CoV-2 binding and neutralizing capacity of Cv2.1169. Based on structural data analysis, we identified the RBM residues in position F486 and N487 as critical for Cv2.1169 binding, acting as anchors that can accommodate the T478K mutation present in several VOCs. Importantly, as previously shown for V_H_1-58-class antibody S2E12, substitutions in position F486 and N487 are unlikely to occur in potential future VOCs because of their deleterious effects in reducing RBD-binding to ACE2 and viral replicative fitness (Greaney et al., 2021; Han et al., 2021). Hence, Cv2.1169 belongs to a class of broad SARS-CoV-2 neutralizers (*i*.*e*., S2E12, A23.58.1, AZD8895 [COV2-2196]) with a high barrier to viral escape and one of the lowest escapability (Dong et al., 2021; Greaney et al., 2021; Han et al., 2021; Wang et al., 2021a). Also, the diminished potency of Cv2.1169 against SARS-CoV-2 Omicron appears moderate when compared to other neutralizing antibodies to the RBD “V_H_1-58 supersite” that drastically reduced or lost their activity against BA.1 and BA.2 (Cameroni et al., 2022; Cao et al., 2022a; Cao et al., 2022b).

SARS-CoV-2 animal models using rodents and non-human primates have been pivotal in demonstrating the *in vivo* prophylactic and therapeutic capacity of human neutralizing anti-spike antibodies (Noy-Porat et al., 2021; Rogers et al., 2020; Rosenfeld et al., 2021). We show that Cv2.1169 IgG efficiently prevents and/or protects animals from infection with SARS-CoV-2 and its VOC Beta. Cv2.1169 was originally expressed by circulating blood IgA-expressing activated memory B cells likely developing in mucosal tissues, and we established that Cv2.1169 IgA antibodies can protect mice from SARS-CoV-2 VOC Beta. Hence, one can assume that such antibodies if locally present at mucosal surfaces, particularly as dimeric IgAs, could efficiently neutralize and/or eliminate virions and therefore, potentially diminish the risk of infection by SARS-CoV-2 variants. In this regard, longer hinge region and multivalency of IgA1 antibody dimers allow enhancing SARS-CoV-2 neutralization *in vitro* as compared to their IgG1 counterparts (Sun et al., 2021; Wang et al., 2021b). In line with this, we found that the loss of neutralization activity of Cv2.1169 against BA.1 and BA.2 was greatly rescued by avidity effects of the antibody produced in its dimeric IgA form.

Several escape mutations in the spike of SARS-CoV-2 variants caused resistance to antibody neutralization, compromising vaccine and therapeutic antibody efficacy (Cameroni et al., 2022; Pinto et al., 2021; Planas et al., 2021b, 2021a). Remarkably, Cv2.1169 and Cv2.3194 demonstrated a broad activity, neutralizing not only VOCs Alpha, Beta, Gamma, Delta and Delta+ but also BA.1 and BA.2, and ranked as the most potent cross-neutralizer when compared to benchmarked antibodies used in clinics. Adjunct to its neutralizing activity, the strong ADCP potential of Cv2.1169 IgG antibodies could contribute to eliminating cell-free and cell-associated virions and stimulating adaptive immunity *via* vaccinal effects (Corti et al., 2021). Taking into account healthcare benefits afforded by antibody therapies to fight COVID-19 (Corti et al., 2021; Singh et al., 2022), and considering the excellent antiviral attributes of Cv2.1169 and Cv2.3194, these two antibodies represent promising candidates for prophylactic and/or therapeutic strategies against COVID-19. Long-acting versions of these broadly SARS-CoV-2 neutralizing antibodies with extended half-life could be used to provide protective immunity in immunocompromised populations (Gentile and Schiano Moriello, 2022).

## Methods

### Human samples

Blood samples from COVID-19 convalescent donors were obtained as part of the CORSER and REACTing French COVID-19 cohorts in accordance with and after ethical approval from all the French legislation and regulation authorities. The CORSER study was registered with ClinicalTrials.gov (NCT04325646), and received ethical approval by the Comité de Protection des Personnes Ile de France III. The REACTing French Covid-19 study was approved by the regional investigational review board (IRB; Comité de Protection des Personnes Ile-de-France VII, Paris, France), and performed according to the European guidelines and the Declaration of Helsinki. All participants gave written consent to participate in this study, and data were collected under pseudo-anonymized conditions using subject coding.

### Serum IgG and IgA purification

All human sera were heat-inactivated at 56°C for 60 min. Human IgG and IgA antibodies were purified from donors’ sera by affinity chromatography using Protein G Sepharose® 4 Fast Flow (GE Healthcare) and peptide M-coupled agarose beads (Invivogen), respectively. Purified serum antibodies were dialyzed against PBS using Slide-A-Lyzer® Cassettes (10K MWCO, Thermo Fisher Scientific).

### Viruses

SARS-CoV-2 BetaCoV/France/IDF0372/2020 (GISAID ID: EPIISL_406596) and D614G (hCoV-19/France/GE1973/2020; GISAID ID: EPI_ISL_414631) strains were supplied by the National Reference Centre for Respiratory Viruses (Institut Pasteur, France) (Grzelak et al., 2020; Planas et al., 2021a). α (B.1.1.7; GISAID ID: EPI_ISL_735391), β (B.1.351; GISAID ID: EPI_ISL_964916), δ (B.1.617.2; GISAID ID: EPI_ISL_2029113), ο BA.1 (GISAID ID: EPI_ISL_6794907) and BA.2 strains were provided by the Virus and Immunity Unit (Institut Pasteur) (Planas et al., 2021b, 2021a, 2022; Bruel et al., 2022). γ variant (P.1.; hCoV-19/Japan/TY7-501/2021; GISAID ID: EPI_ISL_833366) was obtained from Global Health security action group Laboratory Network (Betton et al., 2021). The Beta strain (β, B.1.351; hcoV-19/France/IDF-IPP00078/2021) used for mouse experiments was supplied by the National Reference Centre for Respiratory Viruses (Institut Pasteur, France). Hamsters were infected with the BetaCoV/France/IDF00372/2020 strain (EVAg collection, Ref-SKU: 014V-03890). Viruses were amplified by one or two passages in Vero E6 cell cultures and titrated. The sequence of the viral stocks was verified by RNAseq. All work with infectious virus was performed in biosafety level 3 containment laboratories at Institut Pasteur.

### Expression and purification of viral proteins

Codon-optimized nucleotide fragments encoding stabilized versions of SARS-CoV-2, SARS-CoV-1, MERS-CoV, OC43-CoV, HKU1-CoV, 229E-CoV, NL63-CoV (2P) and BA.1 spike (HexaPro) (S) ectodomains, and SARS-CoV-2 S2 domain, followed by a foldon trimerization motif and C-terminal tags (Hisx8-tag, Strep-tag, and AviTag) were synthesized and cloned into pcDNA3.1/Zeo(+) expression vector (Thermo Fisher Scientific). For competition ELISA experiments, a SARS-CoV-2 S ectodomain DNA sequence without the StrepTag was also cloned into pcDNA3.1/Zeo(+) vector. Synthetic nucleotide fragments coding for Wuhan SARS-CoV-2 RBD, S1 subunit, S1 N-terminal domain (NTD), S1 connecting domain (CD), nucleocapsid protein (N), BA.1 and BA.2 RBDs followed by C-terminal tags (Hisx8-tag, Strep-tag, and AviTag), as well as human angiotensin-converting enzyme 2 (ACE2) (*plus* Hisx8- and Strep-tags), were cloned into pcDNA3.1/Zeo(+) vector. For SARS-CoV-2 RBD variant proteins, mutations (N501Y for the α variant; K417N, E484K and N501Y for the β variant; K471T, E484K and N501Y for the γ variant; L452R and T478K for the δ variant, K417N, L452R and T478K for the δ+ variant; L452R and E484Q for the κ variant) were introduced using the QuickChange Site-Directed Mutagenesis kit (Agilent Technologies) following the manufacturer’s instructions. Glycoproteins were produced by transient transfection of exponentially growing Freestyle 293-F suspension cells (Thermo Fisher Scientific, Waltham, MA) using polyethylenimine (PEI) precipitation method as previously described (Lorin and Mouquet, 2015). Proteins were purified from culture supernatants by high-performance chromatography using the Ni Sepharose® Excel Resin according to manufacturer’s instructions (GE Healthcare), dialyzed against PBS using Slide-A-Lyzer® dialysis cassettes (Thermo Fisher Scientific), quantified using NanoDrop 2000 instrument (Thermo Fisher Scientific), and controlled for purity by SDS-PAGE using NuPAGE 3-8% Tris-acetate gels (Life Technologies), as previously described (Lorin and Mouquet, 2015). AviTagged tri-S and RBD proteins were biotinylated using BirA biotin-protein ligase bulk reaction kit (Avidity, LLC) or Enzymatic Protein Biotinylation Kit (Sigma-Aldrich). SARS-CoV-2 RDB protein was also coupled to DyLight 650 using the DyLight® Amine-Reactive Dyes kit (Thermo Fisher scientific).

For crystallographic experiments, a codon-optimized nucleotide fragment encoding the SARS-CoV-2 RBD protein (residues 331-528), followed by an enterokinase cleavage site and a C-terminal double strep-tag was cloned into a modified pMT/BiP expression vector (pT350, Invitrogen). Drosophila S2 cells were stably co-transfected with pT350 and pCoPuro (for puromycin selection) plasmids. The cell line was selected and maintained in serum-free insect cell medium (HyClone, Cytiva) supplemented with 7 µg/ml puromycin and 1% penicillin/streptomycin antibiotics. Cells were grown to reach a density of 1 × 10^7^ cells/ml, and protein expression was then induced with 4 µM CdCl_2_. After 6 days of culture, the supernatant was collected, concentrated and proteins were purified by high-performance chromatography using a Streptactin column (IBA). The eluate was buffer-exchanged into 10 mM Tris-HCl (pH 8.0), 100 mM NaCl, 2 mM CaCl_2_ using a HiPrep 26/10 Desalting column (GE Healthcare) and subsequently treated with enterokinase overnight at room temperature to remove the strep-tag. Undigested tagged proteins were removed using a Streptactin column, and monomeric untagged protein was purified by size-exclusion chromatography (SEC) using a Superdex 75 column (Cytiva) equilibrated with 10 mM Tris-HCl (pH 8.0), 100 mM NaCl. Purified monomeric untagged protein was concentrated and stored at -80 ºC until used.

For Cryo-EM experiments, a codon-optimized nucleotide fragment encoding the SARS-CoV-2 spike (S) protein (residues 1-1208) was cloned with its endogenous signal peptide in pcDNA3.1(+) vector, and expressed as a stabilized trimeric prefusion construct with six proline substitutions (F817P, A892P, A899P, A942P, K986P, V987P), along with a GSAS substitution at the furin cleavage site (residues 682–685), followed by a Foldon trimerization motif (Hsieh et al., 2020), and C-terminal tags (Hisx8-tag, Strep-tag and AviTag). The recombinant protein, S_6P, was produced by transient transfection of Expi293F™ cells (Thermo Fisher Scientific, Waltham, MA) using FectroPRO® DNA transfection reagent (Polyplus), according to the manufacturer’s instructions. After 5 days of culture, recombinant proteins were purified from the concentrated supernatant by affinity chromatography using a SrepTactin column (IBA), followed by a SEC using a Superose 6 10/300 column (Cytiva) equilibrated in 10 mM Tris-HCl, 100 mM NaCl (pH 8.0). The peak corresponding to the trimeric protein was concentrated and stored at -80 ºC until used.

### Flow cytometry immunophenotyping

Peripheral blood mononuclear cells (PBMC) were isolated from donors’ blood using Ficoll Plaque Plus (GE Healthcare). Human blood B cells and circulating T follicular helper T cells (cTfh) were analyzed using two different fluorescently-labeled antibody cocktails. For B-cell phenotyping, B cells were first isolated from donors’ PBMC by MACS using human CD19 MicroBeads (Miltenyi Biotec). CD19^+^ B cells were then stained using LIVE/DEAD aqua fixable dead cell stain kit (Molecular Probes, Thermo Fisher Scientific) to exclude dead cells. B cells were incubated for 30 min at 4°C with biotinylated tri-S and DyLight 650-coupled RBD, washed once with 1% FBS-PBS (FACS buffer), and incubated for 30 min at 4°C with a cocktail of mouse anti-human antibodies: CD19 Alexa 700 (HIB19, BD Biosciences, San Jose, CA), CD21 BV421 (B-ly4, BD Biosciences), CD27 PE-CF594 (M-T271, BD Biosciences), IgG BV786 (G18-145, BD Biosciences), IgA FITC (IS11-8E10, Miltenyi Biotec, Bergisch Gladbach, Germany), Integrin β7 BUV395 (FIB504, BD Biosciences) and streptavidin R-PE conjugate (Invitrogen, Thermo Fisher Scientific). Cells were then washed and resuspended in FACS buffer. Following a lymphocyte and single cell gating, live cells were gated on CD19^+^ B cells. FACS analyses were performed using a FACS Aria Fusion Cell Sorter (Becton Dickinson, Franklin Lakes, NJ) and FlowJo software (v10.3, FlowJo LLC, Ashland, OR). Immunophenotyping of cTfh subsets was performed on negative fractions from the CD19 MACS. The cTfh antibody panel included: CD3 BV605 (SK7), CD4 PE-CF594 (RPA-T4), CD185/CXCR5 AF-488 (RF8B2), CD183/CXCR3 PE-Cy™5 (1C6/CXCR3), CD196/CCR6 PE-Cy™7 (11A9), CD197/CCR7 AF647 (3D12) (BD Biosciences), CD279/PD1 BV421 (EH12.2H7, BioLegend), and CD278/ICOS PE (ISA-3, Thermo Fisher Scientific). Cells were stained as described above, washed and fixed in 1% paraformaldehyde-PBS. Following a lymphocyte and single cell gating, dead cells were excluded. Flow cytometric analyses of stained cells were performed using a BD LSR Fortessa™ instrument (BD Biosciences), and the FlowJo software (v10.6, FlowJo LLC).

### Single B-cell FACS sorting and expression-cloning of antibodies

Peripheral blood human B cells were isolated and stained as describe above. Single SARS-CoV-2 S^+^ IgG^+^ and IgA^+^ B cells were sorted into 96-well PCR plates using a FACS Aria Fusion Cell Sorter (Becton Dickinson, Franklin Lakes, NJ) as previously described (Tiller et al., 2008). Single-cell cDNA synthesis using SuperScript IV reverse transcriptase (Thermo Fisher Scientific) followed by nested-PCR amplifications of IgH, Igκ and Igλ genes, and sequences analyses for Ig gene features were performed as previously described (Prigent et al., 2016; Tiller et al., 2008). Purified digested PCR products were cloned into human Igγ1-, Igκ- or Igλ-expressing vectors (GenBank# LT615368.1, LT615369.1 and LT615370.1, respectively) as previously described (Tiller et al., 2008). Cv2.1169 were also cloned into human Igγ1^NA^, Igγ1^LALA^ [N297A and L234A/L235A mutations introduced by Site-Directed Mutagenesis (QuickChange, Agilent Technologies)], Igα1 and Fab-Igα1-expressing vectors (Lorin and Mouquet, 2015; Lorin et al., 2022). Cv2.3235, and Cv2.6264 IgH were also cloned into a human Fab-Igγ1-expressing vector (Mouquet et al., 2012). Recombinant antibodies were produced by transient co-transfection of Freestyle^™^ 293-F suspension cells (Thermo Fisher Scientific) using PEI-precipitation method as previously described (Lorin and Mouquet, 2015). The dimeric form of Cv2.1169 IgA1 was produced by co-transfection of Freestyle^™^ 293-F cells with a human J chain pcDNA™3.1/Zeo(+) vector as previously described (Lorin and Mouquet, 2015). Recombinant human IgG, IgA antibodies and Fab were purified by affinity chromatography using Protein G Sepharose® 4 Fast Flow (GE Healthcare), peptide M-coupled agarose beads (Invivogen) and Ni Sepharose® Excel Resin (GE Healthcare), respectively. Monomeric and dimeric Cv2.1169 IgA1 antibodies were separated by SEC using a Superose 6 Increase 10/300 column (Cytiva). After equilibration of the column with PBS, purified IgA antibodies were injected into the column at a flow rate of 0.3 ml/min. Monomers, dimers and multimers were separated upon an isocratic elution with 1.2 CV of PBS. The quality/purity of the different purified fractions was evaluated by SDS-PAGE using 3–8% Tris– Acetate gels (Life Technologies) under non-reducing conditions followed by silver staining (Silver Stain kit, Thermo Scientific). Purified antibodies were dialyzed against PBS. The purified parental IgG1 antibody versions of benchmarked mAbs [REGN10933, REGN10987 (Hansen et al., 2020), CB6 (Shi et al., 2020), LY-CoV555 (Jones et al., 2021), CT-P59 (Kim et al., 2021), COV2-2196, COV2-2130 (Zost et al., 2020b), ADG-2 (Garrett Rappazzo et al., 2021) and S309 (Pinto et al., 2020)] were prepared as described above after cloning of synthetic DNA fragments (GeneArt, Thermo Fisher Scientific) coding for the immunoglobulin variable domains. Antibody preparations for *in vivo* infusions were micro-filtered (Ultrafree®-CL devices - 0.1 µm PVDF membrane, Merck-Millipore, Darmstadt, Germany), and checked for endotoxins levels using the ToxinSensor™ Chromogenic LAL Endotoxin Assay Kit (GenScript).

### ELISAs

ELISAs were performed as previously described (Mouquet et al., 2011, 2012). Briefly, high-binding 96-well ELISA plates (Costar, Corning) were coated overnight with 250 ng/well of purified recombinant Coronavirus proteins and 500 ng/well of a SARS-CoV-2 fusion sequence-containing peptide (KRSFI<u>EDLLFNKVTLAD</u>AGFIK, GenScript Biotech). After washings with 0.05% Tween 20-PBS (washing buffer), plates were blocked 2 h with 2% BSA, 1 mM EDTA, 0.05% Tween 20-PBS (Blocking buffer), washed, and incubated with serially diluted human and rodent sera, purified serum IgA/IgG or recombinant mAbs in PBS. Total sera were diluted 1:100 (for humans and golden hamsters) or 1:10 (for K18-hACE2 mice) following by 7 consecutive 1:4 dilutions in PBS. Purified serum IgG and IgA antibodies were tested at 50 µg/ml and 7 consecutive 1:3 dilutions in PBS. Recombinant IgG1 mAbs were tested at 4 or 10 µg/ml, and 4 to 7 consecutive 1:4 dilutions in PBS. Comparative ELISA binding of Cv2.1169 IgG1 and IgA1 antibodies was performed at a concentration of 70 nM, and 7 consecutive dilutions in PBS. To quantify blood-circulating human Cv2.1169 IgA1 and IgG1 in treated K18-hACE2 mice and golden hamsters, high-binding 96-well ELISA plates (Costar, Corning) were coated overnight with 250 ng/well of purified goat anti-human IgA or IgG antibody (Jackson ImmunoResearch, 0.8 µg/ml final). After washings, plates were blocked, washed, and incubated for 2 h with 1:100 diluted sera from K18-hACE2 mice and golden hamster and seven consecutive 1:3 dilutions in PBS. Cv2.1169 IgA1 or IgG1 antibody at 12 µg/ml and seven consecutive 1:3 dilutions in PBS were used as standards. After washings, the plates were revealed by incubation for 1 h with goat HRP-conjugated anti-mice IgG, anti-golden hamster IgG, anti-human IgG or anti-human IgA antibodies (Jackson ImmunoReseach, 0.8 µg/ml final) and by adding 100 µl of HRP chromogenic substrate (ABTS solution, Euromedex) after washing steps. Optical densities were measured at 405nm (OD_405nm_), and background values given by incubation of PBS alone in coated wells were subtracted. Experiments were performed using HydroSpeed™ microplate washer and Sunrise™ microplate absorbance reader (Tecan Männedorf, Switzerland). For peptide-ELISA, binding of SARS-CoV2 and control IgG antibodies (at 1 µg/ml) to 15-mer S2 overlapping 5-amino acid peptides (n=52, GenScript Biotech, 500 ng/well) was tested using the same procedure as previously described (Wardemann, 2003). For competition ELISAs, 250 ng/well of StrepTag-free tri-S and RBD proteins were coated on ELISA plates (Costar, Corning), which were then blocked, washed, and incubated for 2 h with biotinylated antibodies (at a concentration of 100 ng/ml for tri-S competition and 25 ng/ml for RBD competition) in 1:2 serially diluted solutions of antibody competitors in PBS (IgG concentration ranging from 0.39 to 50 µg/ml). Plates were developed using HRP-conjugated streptavidin (BD Biosciences) as described above. For the competition experiments of tri-S- and RBD-binding to ACE2, ELISA plates (Costar, Corning) were coated overnight with 250 ng/well of purified ACE2 ectodomain. After washings, plates were blocked 2 h with Blocking buffer, PBST-washed, and incubated with recombinant IgG1 mAbs at 2 µg/ml and 7 consecutive 1:2 dilutions in presence of biotinylated tri-S protein at 1 µg/ml in PBS, and at 10 or 100 µg/ml and 7 consecutive 1:2 dilutions in PBS in presence of biotinylated RBD at 0.5 µg/ml. After washings, the plates were revealed by incubation for 30 min with streptavidin HRP-conjugated (BD Biosciences) as described above.Polyreactivity ELISA was performed as previously described (Planchais et al., 2019). Briefly, high-binding 96-well ELISA plates were coated overnight with 500 ng/well of purified double stranded (ds)-DNA, KLH, LPS, Lysozyme, Thyroglobulin, Peptidoglycan from *B. subtilis*, 250 ng/well of insulin (Sigma-Aldrich, Saint-Louis, MO), flagellin from *B. subtilis* (Invivogen), MAPK14 (Planchais et al., 2019), and 125 ng/well of YU2 HIV-1 Env gp140 protein in PBS. After blocking and washing steps, recombinant IgG mAbs were tested at 4 µg/ml and 7 consecutive 1:4 dilutions in PBS. Control antibodies, mGO53 (negative) (Wardemann, 2003), and ED38 (high positive) (Meffre et al., 2004) were included in each experiment. ELISA binding was developed as described above. Serum levels of human IL6, IP10, CXCL13 and BAFF were measured using DuoSet ELISA kits (R&D Systems) with undiluted plasma samples.

### Flow cytometry binding assays

SARS-CoV-2 specificity validation of cloned human IgG antibodies was performed using the S-Flow assay as previously described (Grzelak et al., 2020). To evaluate spike cross-reactivity, Freestyle™ 293-F were transfected with pUNO1-Spike-dfur expression vectors (Spike and SpikeV1 to V11 plasmids, Invivogen) (1.2 µg plasmid DNA *per* 10^6^ cells) using PEI-precipitation method. Forty-eight hours post-transfection, 0.5×10^6^ transfected and non-transfected control cells were incubated with IgG antibodies for 30 min at 4°C (1 µg/ml). After washings, cells were incubated 20 min at 4°C with AF647-conjugated goat anti-human IgG antibodies (1:1000 dilution; Thermo Fisher Scientific) and LIVE/DEAD Fixable Viability dye Aqua (1:1000 dilution; Thermo Fisher Scientific), washed and resuspended in PBS-Paraformaldehyde 1% (Electron Microscopy Sciences). Data were acquired using a CytoFLEX flow cytometer (Beckman Coulter), and analyzed using FlowJo software (v10.7.1; FlowJo LLC). Antibodies were tested in duplicate.

### *HEp-2 IFA* assay

Recombinant SARS-CoV-2 S-specific and control IgG antibodies (mGO53 and ED38) at 100 µg/ml were analyzed by indirect immuno-fluorescence assay (IFA) on HEp-2 cells sections (ANA HEp-2 AeskuSlides®, Aesku.Diagnostics, Wendelsheim, Germany) using the kit’s controls and FITC-conjugated anti-human IgG antibodies as the tracer according to the manufacturer’ instructions. HEp-2 sections were examined using the fluorescence microscope Axio Imager 2 (Zeiss, Jena, Germany), and pictures were taken at magnification x 40 with 5000 ms-acquisition using ZEN imaging software (Zen 2.0 blue version, Zeiss) at the Imagopole platform (Institut Pasteur).

### Infrared immunoblotting

Recombinant tri-S protein was heat-denatured at 100°C for 3 min in loading buffer (Invitrogen) containing 1X sample reducing agent (Invitrogen). Denatured tri-S protein (50 µg total) was separated by SDS-PAGE with a NuPAGE® 4-12% Bis-Tris Gel (1-well, Invitrogen), electro-transferred onto nitrocellulose membranes, and saturated in PBS-0.05% Tween 20 (PBST)-5% dry milk overnight at 4°C. Membranes were inserted into a Miniblot apparatus (Immunetics) and then incubated with human mAbs (at a concentration of 1 µg/ml) and mouse anti-Hisx6 antibody (1 µg/ml, BD Biosciences) in PBS-T 5% dry milk in each channel for 2 h. For dot blotting experiments, denatured tri-S (ranging from 0.125 to 2 µg) was immobilized on dry nitrocellulose membranes for 2 h at room temperature and saturated in PBS-0.05% Tween 20 (PBST)-5% dry milk overnight at 4°C. The membranes were then incubated with human mAbs (at a concentration of 1 µg/ml) and mouse anti-Hisx6 antibody (1 µg/ml, BD Biosciences) in PBS-T 5% dry milk for 2 h. After washing with PBST, membranes were incubated for 1h with 1/25,000-diluted Alexa Fluor 680-conjugated donkey anti-human IgG (Jackson ImmunoResearch) and 1/25,000-diluted IR Dye® 800CW-conjugated goat anti-mouse IgG (LI-COR Biosciences) in PBST-5% dry milk. Finally, membranes were washed, and examined with the Odyssey Infrared Imaging system (LI-COR Biosciences).

### Protein microarray binding analyses

All experiments were performed at 4°C using ProtoArray Human Protein Microarrays (Thermo Fisher Scientific). Microarrays were blocked for 1 h in blocking solution (Thermo Fisher), washed and incubated for 1h30 with IgG antibodies at 2.5 µg/ml as previously described (Grzelak et al., 2020). After washings, arrays were incubated for 1h30 with AF647-conjugated goat anti-human IgG antibodies (at 1 µg/ml in PBS; Thermo Fisher Scientific), and revealed using GenePix 4000B microarray scanner (Molecular Devices) and GenePix Pro 6.0 software (Molecular Devices) as previously described (Planchais et al., 2019). Fluorescence intensities were quantified using Spotxel® software (SICASYS Software GmbH, Germany), and mean fluorescence intensity (MFI) signals for each antibody (from duplicate protein spots) was plotted against the reference antibody mGO53 (non-polyreactive isotype control) using GraphPad Prism software (v8.1.2, GraphPad Prism Inc.). For each antibody, Z-scores were calculated using ProtoArray® Prospector software (v5.2.3, Thermo Fisher Scientific), and deviation (σ) to the diagonal, and polyreactivity index (PI) values were calculated as previously described (Planchais et al., 2019). Antibodies were defined as polyreactive when PI > 0.21.

### Surface plasmon resonance

Surface plasmon resonance (SPR)-based technology (Biacore 2000, Biacore, Uppsala, Sweden) was used to assess kinetics of interaction of mAbs with SARS CoV2 proteins – trimer S, S1 and RBD. Antibodies (Cv2.1169, Cv2.1353, Cv2.3194, Cv2.3235 and Cv2.5213) and ACE2 ectodomain were covalently coupled to CM5 sensor chips (Biacore) using amino-coupling kit (Biacore) according to the manufacturer’s procedure. In brief, IgG antibodies and ACE2 protein were diluted in 5 mM maleic acid solution, pH 4 to a final concentration of 10 μg/ml and injected over sensor surfaces pre-activated by a mixture of 1-Ethyl-3-(3-dimethylaminopropyl) carbodiimide and N-hydroxysuccinimide. Uncoupled carboxyl groups were blocked by exposure to 1M solution of ethanolamine.HCl (Biacore). Immobilization densities were 500 RU and 1000 RU for IgG antibodies and ACE2, respectively. All analyses were performed using HBS-EP buffer (10 mM HEPES pH 7.2; 150 mM NaCl; 3 mM EDTA, and 0.005 % Tween 20). The flow rate of buffer during all real-time interaction measurements was set at 30 µl/min. All interactions were performed at temperature of 25 °C. SARS CoV-2 tri-S and S1 proteins were serially diluted (two-fold step) in HBS-EP in the range of 40 – 0.156 nM. Same range of concentrations was used for RBD with exception of low affinity interactions where the concentration range 1280 – 10 nM was applied. The association and dissociation phases of the binding of viral proteins to the immobilized antibodies and ACE2 were monitored for 3 and 4 minutes, respectively. The binding of the proteins to reference channel containing carboxymethylated dextran only was used as negative control and was subtracted from the binding during data processing. The sensor chip surfaces were regenerated by 30 s exposure to 4M solution of guanidine-HCl (Sigma-Aldrich). The evaluation kinetic parameters of the studied interactions were performed by using BIAevaluation version 4.1.1 Software (Biacore).

### SARS-CoV-2 S-Fuse neutralization assay

S-Fuse cells (U2OS-ACE2 GFP1-10 or GFP 11 cells) were mixed (ratio 1:1) and plated at a density of 8 × 10^3^ per well in a μClear 96-well plate (Greiner Bio-One) as previously described (Buchrieser et al., 2020). SARS-CoV-2 and VOC viruses (MOI 0.1) were incubated with recombinant IgG1, monomeric and dimeric IgA1 mAbs at 35 nM or 7 nM, and 11 consecutive 1:4 dilutions in culture medium for 30 min at room temperature and added to S-Fuse cells. The cells were fixed, 18 h later, in 2% paraformaldehyde, washed and stained with Hoechst stain (dilution 1:1000; Invitrogen). Images were acquired with an Opera Phenix high-content confocal microscope (Perkin Elmer). The area displaying GFP expression and the number of nuclei were quantified with Harmony software 4.8 (Perkin Elmer). The percentage neutralization was calculated from the GFP-positive area as follows: 100 × (1 – (value with IgA/IgG – value in “non-infected”) / (value in “no IgA/IgG” – value in “non-infected”)). IC_50_ values were calculated using Prism software (v.9.3.1, GraphPad Prism Inc.) by fitting replicate values using the four-parameters dose–response model (variable slope).

### In vitro SARS-CoV-2 pseudoneutralization assay

The SARS-CoV-2 pseudoneutralization assay was performed as previously described (Anna et al., 2021; Grzelak et al., 2020). Briefly, 2×10^4^ 293T-ACE2-TMPRSS2 were plated in 96-well plates. Purified serum IgA and IgG antibodies were tested at 250 µg/ml and 7 consecutive 1:2 dilutions in PBS (or in Penicillin/Streptomycin-containing 10%-FCS DMEM), and incubated with spike-pseudotyped lentiviral particles for 15-30 minutes at room temperature before addition to the cells. Recombinant IgG1, IgA1 or Fab-IgA mAbs were also tested at 70 or 350 nM, and 11 consecutive 1:3 dilutions in PBS. After a 48h incubation at 37°C in 5% CO2, the revelation was performed using the ONE-Glo™ Luciferase Assay System (Promega), and the luciferase signal was measured with EnSpire® Multimode Plate Reader (PerkinElmer). The percentage of neutralization was calculated as follow: 100 x (1 - mean (luciferase signal in sample duplicate) / mean (luciferase signal in virus alone)). Individual experiments were standardized using Cv2.3235 antibody. IC_50_ values were calculated as described above.

### Antibody-dependent cellular phagocytosis (ADCP) assay

PBMC were isolated from healthy donors’ blood (Etablissement Français du Sang) using Ficoll Plaque Plus (GE Healthcare). Primary human monocytes were purified from PBMC by MACS using Whole Blood CD14 MicroBeads (Miltenyi Biotech). Biotinylated-SARS-CoV-2 tri-S proteins were mixed with FITC-labelled NeutrAvidin beads (1 µm, Thermo Fisher Scientific) (1 µg of tri-S for 1 µl of beads), and incubated for 30 min at room temperature. After PBS washings, tri-S coupled-beads 1:500-diluted in DMEM were incubated for 1 h at 37°C with human IgG1 mAbs (at 3 µg/ml). tri-S-beads-antibody mixtures were then incubated with 7.5 × 10^4^ human monocytes for 2 h at 37°C. Following washings with 0.5% BSA, 2 mM EDTA-PBS, cells were fixed with 4% PFA-PBS and analyzed using a CytoFLEX flow cytometer (Beckman Coulter). ADCP assays were performed in two independent experiments, and analyzed using the FlowJo software (v10.6, FlowJo LLC). Phagocytic scores were calculated by dividing the fluorescence signals (% FITC-positive cells x geometric MFI FITC-positive cells) given by anti-SARS-CoV-2 spike antibodies by the one of the negative control antibody mGO53.

### Antibody-dependent cellular cytotoxicity (ADCC) assay

The ADCC activity of anti-SARS-CoV2 S IgG antibodies was determined using the ADCC Reporter Bioassay (Promega) as previously described (Dufloo et al., 2021). Briefly, 5×10^4^ Raji-Spike cells were co-cultured with 5×10^4^ Jurkat-CD16-NFAT-rLuc cells in presence or absence of SARS-CoV2 S-specific or control mGO53 IgG antibody at 10 µg/ml or 50 µg/ml and 10 consecutive 1:2 dilutions in PBS. Luciferase was measured after 18 h of incubation using an EnSpire plate reader (PerkinElmer). ADCC was measured as the fold induction of Luciferase activity compared to the control antibody. Experiments were performed in duplicate in two independent experiments.

### Complement-dependent cytotoxicity (CDC) assay

The CDC activity of anti-SARS-CoV2 S IgG antibodies was measured using SARS-CoV-2 Spike-expressing Raji cells as previously described (Pelleau et al., 2020). Briefly, 5×10^4^ Raji-Spike cells were cultivated in the presence of 50% normal or heat-inactivated human serum, and with or without IgG antibodies (at 10 µg/ml or 50 µg/ml and 10 consecutive 1:2 dilutions in PBS). After 24h, cells were washed with PBS, and incubated for 30 min at 4°C the live/dead fixable aqua dead cell marker (1:1,000 in PBS; Life Technologies) before fixation. Data were acquired on an Attune NxT instrument (Life Technologies). CDC was calculated using the following formula: 100 × (% of dead cells with serum − % of dead cells without serum) / (100 − % of dead cells without serum). Experiments were performed in duplicate in two independent experiments.

### Crystallization and structure determinations

The Fab of anti-SARS-CoV-2 S antibody CR3022 (Ter Meulen et al., 2006), served as a crystallization chaperone molecule, and was produced and purified as described above (section with heading *Single B-cell FACS sorting and expression-cloning of antibodies*) (Koide, 2009). The purified RBD protein was incubated overnight at 4 ºC with the Fabs with an RBD-Fab molar ratio of 2:1 (2:1:1 for the ternary complex RBD-Cv2.1169-CR3022). Each binding reaction was loaded onto a Superdex200 column (Cytiva) equilibrated in 10 mM Tris-HCl (pH 8.0), 100 mM NaCl. The fractions corresponding to the complexes were pooled, concentrated to 9-10 mg/ml and used in crystallization trials at 18 ºC using the sitting-drop vapor diffusion method. The RBD-Cv2.2325 Fab complex crystalized with 0.1 M ammonium citrate (pH 7.0), 12% PEG 3350, while crystals for RBD-Cv2.6264 Fab were obtained with 0.1 M NaAc, 7% PEG 6000, 30% ethanol. The RBD-Cv2.1169-CR3022 crystals grew in the presence of 6% PEG 8000, 0.5 M Li_2_SO4. Crystals were flash-frozen by immersion into a cryo-protectant containing the crystallization solution supplemented with 30% (v/v) glycerol (RBD-Cv2.2325; RBD-Cv2.1169-CR3022) or 30% (v/v) ethylenglycol (RBD-Cv2.6264), followed by flash-freezing in liquid nitrogen. Data collection was carried out at SOLEIL synchrotron (St Aubin, France). Data were processed, scaled and reduced with XDS and AIMLESS, and the structures were determined by molecular replacement using Phaser from the suite PHENIX (Liebschner et al., 2019) and search ensembles obtained from the PBDs 6M0J (RBD), 5I1E (Cv2.2325), 5VAG (Cv2.6264), 7K3Q (Cv2.1169) and 6YLA (CR3022). The final models were built by combining real space model building in Coot (Emsley et al., 2010) with reciprocal space refinement with phenix.refine. The final models were validated with Molprobity (Williams et al., 2018). Epitope and paratope residues, as well as their interactions, were identified by accessing PISA at the European Bioinformatics Institute (www.ebi.ac.uk/pdbe/prot_int/pistart.html) (Krissinel and Henrick, 2007). Superpositions and figures were rendered using Pymol and UCSF Chimera (Pettersen et al., 2004).

### Cryo-electron microscopy

The S_6P protein was incubated with the Cv2.1169 IgA Fab at a 1:3.6 (trimer:Fab) ratio and a final trimer concentration of 0.8 µM for 1h at room temperature. 3 µl aliquots of the sample were applied to freshly glow discharged R 1.2/1.3 Quantifoil grids prior to plunge freezing using a Vitrobot Mk IV (Thermo Fischer Scientific) at 8 ºC and 100% humidity (blot 4s, blot force 0). Data for the complex were acquired on a Titan Krios transmission electron microscope (Thermo Fischer Scientific) operating at 300 kV, using the EPU automated image acquisition software (Thermo Fisher Scientific). Movies were collected on a Gatan K3 direct electron detector operating in counting mode at a nominal magnification of 105,000x (0.85 Å/pixel) using defocus range of -1.0 µm to -3.0 µm. Movies were collected over a 2 s exposure and a total dose of ∼45 e-/Å^2^.

### Image processing

All movies were motion-corrected and dose-weighted with MotionCorr2 (Zheng et al., 2017) and the aligned micrographs were used to estimate the defocus values with patchCTF within cryosparc (Punjani et al., 2017). CryoSPARC blob picker was used for automated particle picking and the resulting particles used to obtain initial 2D references, which were then used to auto-pick the micrographs. An initial 3D model was obtained in cryosparc and used to perform a 3D classification without imposing any symmetry in Relion (Zivanov et al., 2018). The best class was selected and subjected to 3D, non-uniform refinement in cryosparc (Punjani et al., 2020).

### SARS-CoV-2 infection and treatment in K18-hACE2 mice

B6.Cg-Tg(K18-ACE2)2Prlmn/J mice (stock #034860) were imported from The Jackson Laboratory (Bar Harbor, ME, USA) and bred at the Institut Pasteur under strict SPF conditions. Infection studies were performed on 6 to 16 wk-old male and female mice, in animal biosafety level 3 (BSL-3) facilities at the Institut Pasteur, in Paris. All animals were handled in strict accordance with good animal practice. Animal work was approved by the Animal Experimentation Ethics Committee (CETEA 89) of the Institut Pasteur (project dap 200008 and 200023) and authorized by the French legislation (under project 24613) in compliance with the European Communities Council Directives (2010/63/UE, French Law 2013–118, February 6, 2013) and according to the regulations of Pasteur Institute Animal Care Committees before experiments were initiated. Anesthetized (ketamine/xylazine) mice were inoculated intranasally (i.n.) with 1 ×10^4^ or 1 ×10^5^ PFU of SARS-CoV-2 (20 µl/nostril). Six or 22 h post-inoculation, mice received an intraperitoneal (i.p.) injection of 5, 10, 20 or 40 mg/kg of Cv2.1169 IgG or IgA antibody, and of mGO53 control IgG or IgA antibody. Clinical signs of disease (ruffled fur, hunched posture, reduced mobility and breathing difficulties) and weight loss were monitored daily during 20 days. Mice were euthanized when they reached pre-defined end-point criteria. Sera were extracted from blood collected by puncture of the retromandibular vein.

### SARS-CoV-2 infection and treatment in golden hamsters

Golden Syrian hamsters (*Mesocricetus auratus;* RjHan:AURA) of 5-6 weeks of age (average weight 60-80 grams) were purchased from Janvier Laboratories (Le Genest-Saint-Isle, France), and handled under specific pathogen-free conditions. Golden hamsters were housed and manipulated in class III safety cabinets in the Pasteur Institute animal facilities accredited by the French Ministry of Agriculture for performing experiments on live rodents, with *ad libitum* access to water and food. Animal work was approved by the Animal Experimentation Ethics Committee (CETEA 89) of the Institut Pasteur (project dap 200023) and authorized by the French legislation (project #25326) in compliance with the European Communities Council Directives (2010/63/UE, French Law 2013–118, February 6, 2013) and according to the regulations of Pasteur Institute Animal Care Committees before experiments were initiated. Animal infection was performed as previously described (de Melo et al., 2021). Briefly, anesthetized animals were intranasally infected with 6×10^4^ plaque-forming units (PFU) of SARS-CoV-2 (BetaCoV/France/IDF00372/2020) (50 µl/nostril). Mock-infected animals received the physiological solution only. Four or 24 h post-intranasal inoculation, hamsters received an intraperitoneal (i.p.) injection of 10 or 5 mg/kg of Cv2.1169 IgG or IgA antibody, as well as the mGO53 control antibody or PBS. All hamsters were followed-up daily when the body weight and the clinical score were noted. At day 5 post-inoculation, animals were euthanized with an excess of anesthetics (ketamine and xylazine) and exsanguination (AVMA Guidelines 2020). Blood samples were collected by cardiac puncture; after coagulation, tubes were centrifuged at 1,500 x g during 10 min at 4°C, and sera were collected and frozen at -80°C until further analyses. The lungs were weighted and frozen at -80°C until further analyses. Frozen lungs fragments were weighted and homogenized with 1 ml of ice-cold DMEM (31966021, Gibco) supplemented with 1% penicillin/streptomycin (15140148, Thermo Fisher) in Lysing Matrix M 2 ml tubes (116923050-CF, MP Biomedicals) using the FastPrep-24™ system (MP Biomedicals), and the following scheme: homogenization at 4.0 m/s during 20 sec, incubation at 4°C during 2 min, and new homogenization at 4.0 m/s during 20 sec. The tubes were centrifuged at 10,000 x g during 1 min at 4°C. The supernatants were titrated on Vero-E6 cells by classical plaque assays using semisolid overlays (Avicel, RC581-NFDR080I, DuPont) and expressed and PFU/100 mg of tissue (Baer and Kehn-Hall, 2014). Frozen lungs fragments were homogenized with Trizol (15596026, Invitrogen) in Lysing Matrix D 2 ml tubes (116913100, MP Biomedicals) using the FastPrep-24™ system (MP Biomedicals), and the following scheme: homogenization at 6.5 m/s during 60 sec, and centrifugation at 12,000 x g during 2 min at 4°C. The supernatants were collected and the total RNA was then extracted using the Direct-zol RNA MiniPrep Kit (R2052, Zymo Research) and quantified using NanoDrop 2000. The presence of genomic SARS-CoV-2 RNA in these samples was evaluated by one-step RT-qPCR in a final volume of 25 μl *per* reaction in 96-well PCR plates using a thermocycler (7500t Real-time PCR system, Applied Biosystems) as previously described (Melo et al., 2021). Viral load quantification (expressed as RNA copy number/µg of RNA) was assessed by linear regression using a standard curve of six known quantities of RNA transcripts containing the *RdRp* sequence (ranging from 10^7^ to 10^2^ copies).

### Quantification and statistical analysis

The numbers of V_H_, Vκ and Vλ mutations were compared across groups of antibodies using unpaired Student’s t test with Welch’s correction. Bivariate correlations were assayed using two-tailed Pearson correlation test. Statistical and analyses were performed using GraphPad Prism software (v.8.2, GraphPad Prism Inc.). Volcano plot comparing gene features (n=206 parameters) of tri-S^+^ B cells and normal memory B-cells (mB) was also performed using GraphPad Prism software (v.8.4, GraphPad Prism Inc.). The y axis indicates the statistics expressed as -log_10_ (p-values) and the x axis represents the differences between the group means for each parameter. The Barnes-Hut implementation of *t*-distributed stochastic neighbor embedding (t-SNE) was computed using FlowJo software (v.10.3, FlowJo LLC, Ashland, OR) with 2000 iterations and a perplexity parameter of 200. Colors represent density of surface expression markers or cell-populations varying from low (blue) to high (red). Circos plot linking antibody sequences with at least 75% identity within their CDR_H_3 was performed using online software at http://mkweb.bcgsc.ca/circos. Phylogenetic tree was built using CLC Main Workbench (Qiagen) on aligned V_H_ sequences using the Neighbor-Joining method with a bootstrap analysis on 100 replicates. Mouse survival were compared across groups using a Kaplan-Meier analysis and Log-rank Mantel-Cox test (GraphPad Prism, v8.2, GraphPad Prism Inc.). Groups of golden Syrian hamsters were compared across analyses using two-tailed Mann-Whitney test (GraphPad Prism, v.8.2, GraphPad Prism Inc.). Principal component analysis (PCA) was performed using the prcomp() function in R Studio Server (v1.4.1103). PCA plots of individuals [fviz_pca_ind()], variables [fviz_pca_var()], and biplots [fviz_pca_biplot()], were generated using the factoextra package (v1.0.7, https://CRAN.R-project.org/package=factoextra). Spearman rank correlations were used to establish multiparameter associations. All correlograms and scatterplots were created using the corrplot and plot R functions, respectively. Correlation plots were generated using GraphPad Prism (v6.4, GraphPad Prism Inc.).

## Acknowledgements

We are grateful to all participants who consented to be part of this study. We thank the members of the Crystallography core facility (Institut Pasteur) for carrying out robot-driven crystallization screeenings, and of the beamlines Proxima 1 and Proxima 2 at the French national synchrotron facility (SOLEIL, St Aubin, France). We also thank the NanoImaging core facility (Institut Pasteur) for support with sample preparation and image acquisition. The NanoImaging Core was created with the help of a grant from the French Government’s Investissements d’Avenir program (EQUIPEX CACSICE, ANR-11-EQPX-0008). This work was supported by grants from the ANR REACTing Covid19 (#20RR028-00), the European Commission Horizon 2020 program (RECoVER project, #101003589), the Institut Pasteur Task Force COVID-19 (2019-NCOV THERAMAB project), the Fondation de France (#00106077), and partly by a SpikImm-Institut Pasteur R&D program. H.M. also received core funding from the Institut Pasteur, and the INSERM. M. Backovic (2020-TooLab project) received support from the “URGENCE COVID-19” fundraising campaign (Institut Pasteur). I.F. was a recipient of an ANRS post-doctoral fellowship. We thank the members of the SpikImm team for their support and helpful discussions.

## Author Contributions

H.M. conceived and supervised the study. J.D.D, F.A., H.B., E. S-L., X.M., F.A.R., O.S. and H.M. supervised the experiments. C.P., I.F., T.B., G.DDM, M.P., M.B, J.D., L. M-A., J. C., E.G., B.V., L.C., L.G., D.P., I.S., F.G-B, and H.M. designed, performed and analyzed the experiments. M. Backovic, P. G-C. collected and/or processed XRC and cryo-EM data. M. C-G., French COVID Cohort Study Group, CORSER Study Group and M-N.U. provided human samples and personal data. M. Boullé, P.C., and S.VDW. contributed with key reagents/assays and expertise. C.P. and H.M. wrote the manuscript with contributions from all the authors.

