## Supplementary Material for "Potent Human Broadly SARS-CoV-2 Neutralizing IgA and IgG Antibodies Effective Against Omicron BA.1 and BA.2"

#### Supplementary Figure Legends

**Figure S1. SARS-CoV-2 reactivity of sera, purified polyclonal and monoclonal antibodies from COVID-19 convalescents. Related to Figure 1. (A)** Graph comparing the single-dilution OD measurements (1:400) (X axis) and AUC values (Y axis) measured with serially-diluted sera from convalescent COVID-19 individuals in the CORSER (n=212) and French COVID-19 cohorts (n=159), and pre-epidemic donors (n=100) for the ELISA IgG antibody binding to SARS-CoV-2 tri-S as previously reported (Grzelak et al., 2020). **(B)** ELISA graphs showing the reactivity of serum IgG (blue) and IgA (red) antibodies from selected convalescent COVID-19 individuals in the CORSER (n=8) and French COVID-19 (n=34) cohorts against SARS-CoV-2 tri-S and RBD proteins. Samples were also tested against MERS tri-S to assay for cross-reactivity against another β-coronavirus. Means of duplicate values are shown. DF, dilution factor. **(C)** Correlation plots comparing the AUC binding values of serum IgG and IgA antibodies to SARS-CoV-2 tri-S, MERS-CoV tri-S and RBD proteins as determined in (A). p values were calculated using two-tailed Pearson correlation test. **(D)** ELISA graphs showing the reactivity of purified IgG (blue) and IgA (red) serum antibodies from selected donors (n=10) against SARS-CoV-2 protein and protein sub-units. Means of duplicate values are shown. **(E)** Same as in (D) but for trimeric spike proteins from other coronaviruses. **(F)** ELISA graphs showing the reactivity of antibodies cloned from SARS-CoV-2 S-captured memory B cells (n=133) against the SARS-CoV-2 tri-S protein. Means of duplicate values are shown.

**Figure S2. Humoral immune features of COVID-19 convalescents and SARS-CoV-2 S-specific memory B cells. Related to Figure 2. (A)** Correlograms showing the correlation analyses of the humoral immune parameters measured in COVID-19 convalescents including antibody titers, neutralizing activity, and memory B-cell subset frequencies. For each pair of compared parameters, Spearman correlation coefficients (color coded) with their corresponding p-value are shown. ***$p<0.0001$, **$p<0.001$, *$p<0.05$. **(B)** Heatmap showing the correlation analyses between the frequency of memory B-cell and cTfh subsets (%) measured in COVID-19 convalescents. Cells are color-coded according to the value of Spearman correlation coefficients with the corresponding p-values indicated in the center. **$p<0.001$, *$p<0.05$. **(C)** Pie charts comparing the distribution of $V_H$ / $J_H$ gene usage of blood SARS-CoV-2 spike-specific $IgG^+$/$IgA^+$ memory B cells and $IgG^+$ memory B cells from SARS-CoV-2-unexposed healthy individuals (mB). The number of antibody sequences analyzed is indicated in the center of each pie chart. Groups were compared using 2 × 5 Fisher's Exact test. **(D)** Bar graph comparing the distribution of $CDR_H3$ lengths (top) and positive charge numbers (bottom) between blood SARS-CoV-2 spike-specific $IgG^+$/$IgA^+$ memory B cells and $IgG^+$ memory B cells from unexposed individuals (mB). Groups were compared using 2 × 5 Fisher's Exact test. **(E)** Same as in (C) but according to the anti-spike antibody specificity (S1, S2 or RBD). **(F)** Bar graph comparing the distribution of IgG subtypes between blood SARS-CoV-2 spike-specific $IgG^+$/$IgA^+$ memory B cells and $IgG^+$ memory B cells from unexposed individuals (mB). Groups were compared using 2 × 5 Fisher's Exact test. **(G)** Pie charts showing the κ- vs λ-Ig chain usage of blood SARS-CoV-2 spike-specific $IgG^+$/$IgA^+$ memory B cells and $IgG^+$ memory B cells from unexposed individuals (mB). Groups were compared using 2 × 2 Fisher's Exact test. **(H)** Violin plots comparing the number of mutations in VH, Vκ and Vλ genes in SARS-CoV-2 spike-, S1-, S2-, and RBD-specific and control memory B cells. Numbers of mutations were compared across groups of antibodies using the unpaired Student t test with Welch's correction. **(I)** Same as in (C) but for Vκ / Jκ and Vλ / Jλ gene usages. **(J)** Bar graphs comparing the distribution of single immunoglobulin genes, $V_H$ (top) and $V_L$ (bottom), expressed by SARS-CoV-2 spike-specific and control $IgG^+$ memory B cells. Groups were compared using 2 × 2 Fisher's Exact test. **(K)** Same as in (D) but for CDRκ3 and CDRλ3 lengths. **(L)** Circos plots comparing the $V_H(D_H)J_H$ and $V_LJ_L$ rearrangement frequencies between SARS-CoV-2 spike-specific $IgA^+$/$IgG^+$ memory B cells and $IgG^+$ memory B cells from unexposed individuals (mB). Groups were compared using 2 × 5 Fisher's Exact test.

**Figure S3. Binding characteristics of potent anti-RBD antibody neutralizers. Related to Figures 3 and 4. (A)** Infrared immunoblot showing the reactivity of SARS-CoV-2 S-specific IgG antibodies (n=101) to denatured SARS-CoV-2 tri-S protein. Immunoreactive green bands correspond to denatured SARS-CoV-2 tri-S protein revealed with an anti-6xHis tag antibody. The red band (yellow when merged) indicates the SARS-CoV-2 antibody Cv2.3132 recognizing denatured tri-S protein. **(B)** Infrared dot blot showing the reactivity of Cv2.3132 antibody to denatured SARS-CoV-2 tri-S protein at various concentrations. mGO53 is a non-SARS-CoV-2 isotype control. Cv2.1169 was included for comparison. **(C)** Graphs showing the reactivity of Cv2.3132 IgG antibody against 15-mer S2 overlapping 5-amino acid peptides (n=52). mGO53 is a non-SARS-CoV-2 isotype control. Means ± SD of duplicate values are shown. **(D)** Competition ELISA graphs showing the IgG binding to SARS-CoV-2 tri-S (top) and RBD (bottom) of selected biotinylated SARS-CoV-2 S-specific antibodies in presence of the corresponding non-biotinylated IgG antibodies as potential competitors. Means ± SD of duplicate values are shown. **(E)** ELISA graphs showing the reactivity of SARS-CoV-2 RBD-specific IgG antibodies to RBD proteins from SARS-CoV-2 viral variants α, β and γ. Means ± SD of duplicate values are shown. **(F)** Competition ELISA graphs showing the binding of biotinylated RBD proteins from SARS-CoV-2 and viral variants (α, β and γ) to soluble ACE2 ectodomain in presence of SARS-CoV-2 S-specific IgG antibodies as potential competitors. Framed graphs show selected IgG competitors tested at a higher concentration against α and β RBD proteins. Means ± SD of duplicate values are shown. **(G)** Same as in (E) but for RBD proteins from SARS-CoV-2 viral variants κ, δ and $\delta^+$. **(H)** Same as in (F) but for RBD proteins from SARS-CoV-2 viral variants κ, δ and $\delta^+$. **(I)** ELISA graphs comparing the reactivity of the monomeric IgG / IgA and dimeric IgA (dIgA) antibody forms of Cv2.1169 to SARS-CoV-2 tri-S, S1 and RBD, and to RBD proteins from SARS-CoV-2 viral variants (α, β, γ, δ, $\delta^+$ and κ). Means ± SD of duplicate values are shown.

**Figure S4. Poly- and self-reactivity assessment of potent SARS-CoV-2 neutralizing antibodies. Related to Figure 4. (A)** ELISA graphs showing the reactivity of selected SARS-CoV-2 neutralizing antibodies against dsDNA (DNA), flagellin (Fla), YU2 HIV-1 Env (gp140), insulin (INS), keyhole limpet hemocyanin (KLH), lipopolysaccharide (LPS), lysozyme (LZ), MAPK-14 (MAPK), proteoglycan (PG), and thyroglobulin (Tg), mGO53 (Wardemann, 2003) and ED38 (Meffre et al., 2004) are negative and positive control antibody, respectively. Anti-SARS-CoV-2 S antibody Cv2.3132 showing HEp-2 reactivity in (C) was included for comparison. Mean of duplicate values are shown. **(B)** Heatmap comparing the area under the curve (AUC) values determined from the ELISA binding analyses shown in (A). Darker blue colors indicate high binding while light colors show moderate binding (white = no binding). **(C)** Microscopic images showing the reactivity of selected SARS-CoV-2 antibodies to HEp2-expressing self-antigens assayed by indirect immunofluorescence assay. The negative (mGO53), low-positive (ED38), and kit's positive (Ctr+) controls were included in the experiment. HEp-2-reactive anti-SARS-CoV-2 S antibody Cv2.3132 was also included for comparison. The scale bars represent 40 µm. **(D)** Bar graph showing the HEp-2 reactivity of selected SARS-CoV-2 antibodies as measured by ELISA. Means ± SD of duplicate values are shown. Ctr+ and Ctr- are the positive and negative control of the kit, respectively. **(E)** Representative microarray plots showing the reactivity of selected SARS-CoV-2 antibodies to human proteins. Plots (top) showing the mean fluorescence intensity (MFI) values given on each protein spot by the reference (Ref: mGO53) and test antibody on the *y* and *x* axis, respectively. Each dot represents the average of duplicate array proteins. Plots (bottom) showing the z-scores given on a single protein by the reference (Ref: mGO53, y axis) and test antibody (x axis). **(F)** Frequency histograms showing the $\log_{10}$ protein displacement (σ) of the MFI signals for the selected SARS-CoV-2 antibodies compared to non-reactive antibody mGO53 obtained from two independent experiments (Array #1 and #2). The polyreactivity index (PI) corresponds to the Gaussian mean of all array protein displacements. Blue and red histograms indicate non-polyreactive and polyreactive antibodies, respectively.

**Figure S5. Cross-neutralization and Fc-dependent effectors functions of potent SARS-CoV2 neutralizers. Related to Figure 4. (A)** Graphs comparing the NK-mediated ADCC activity of selected neutralizing (nAbs) and non-neutralizing (non-nAbs) SARS-CoV-2 S-specific antibodies. Means ± SD of duplicate values are shown. **(B)** Same as in (A) but for the CDC activity. **(C)** ADCP activity of Cv2.1169. The dot plot (left) shows the monocyte-mediated ADCP activity of Cv2.1169 IgG at a concentration of 1 and 10 µg/ml. Each dot corresponds to a donor of primary monocytes (n=6). Graph comparing the ADCP activity of Cv2.1169 expressed as recombinant IgG1, $IgG1^{NA}$, $IgG1^{LALA}$, monomeric IgA1 (mIgA1) and dimeric IgA1 (dIgA1) antibodies. mGO53 is the negative isotype control, and ADCP-induced S309 IgG1 was included for comparison. PS, phagocytic score. Means of duplicate values are shown. **(D)** Graphs showing the neutralization curves of SARS-CoV-2 and selected VOCs by potent anti-RBD IgG antibodies as determined with the S-Fuse neutralization assay. Error bars indicate the SD of duplicate values. $IC_{50}$ values are indicated in the top left-hand corner (in blue). ND, not determined. **(E)** Graphs showing the neutralization curves of SARS-CoV-2 and selected VOCs by Cv2.1169 and Cv2.3194 IgG antibodies as determined with the pseudo-neutralization assay. Error bars indicate the SD of duplicate values. $IC_{50}$ values are indicated in the top left-hand corner (in blue).

**Figure S6. Comparative analyses of Cv2.1169 with benchmarked antibodies. Related to Figures 5 and 6. (A)** Heatmap comparing the ELISA binding to the selected SARS-CoV-2 proteins of Cv2.1169, Cv2.3194 and benchmarked neutralizing antibodies in clinical use or in development. Darker blue colors indicate high binding while light colors show moderate binding or competition (white = 0, no binding). Means of duplicate AUC values are shown in each cell. **(B)** Heatmap comparing the tri-S- and RBD-ACE2 blocking capacity of Cv2.1169, Cv2.3194 and benchmarked neutralizing antibodies. Darker blue colors indicate high competition while light colors show moderate competition (white = 0, no competition). Means of duplicate values (% binding inhibition) are shown in each cell. NT, not tested. **(C)** Heatmap comparing the *in vitro* neutralizing activity of Cv2.1169 and benchmarked neutralizing antibodies against the selected SARS-CoV-2 viral variants. Means of triplicate $IC_{50}$ values in pM are shown in each cell. White color indicates that 50% neutralization was not reached at the maximum antibody concentration of 25 nM. **(D)** Heatmaps showing the competition potential of Cv2.1169, Cv2.3194 and benchmarked neutralizing antibodies for the ELISA binding to tri-S and RBD proteins. Darker blue colors indicate high competition while light colors show moderate competition (white = 0, no competition). Means of duplicate values (% binding inhibition) are shown in each cell.

**Figure S7. $V_H$1-58-encoded SARS-CoV-2 spike-captured memory B-cell antibodies. Related to Figure 4. (A)** Amino acid alignment of the heavy chains (IgH, left) and light chains (IgL, right) of the $V_H$1-58-encoded human antibodies produced from SARS-CoV-2 spike-captured memory B-cell antibodies. Dendrograms showing the relationship between $V_H$1-58-encoded human antibodies generated from the IgH and IgL sequence alignments are shown at the bottom. **(B)** ELISA graphs showing the reactivity of $V_H$1-58-encoded antibodies against the SARS-CoV-2 tri-S protein. Means ± SD of duplicate values are shown. **(C)** ELISA graphs comparing the binding of Cv2.5179 and Cv2.1169 antibodies to RBD proteins. Means ± SD of duplicate values are shown. **(D)** Competition ELISA graphs showing the binding of biotinylated SARS-CoV-2 tri-S and RBD proteins to the immobilized soluble ACE2 ectodomain in presence of Cv2.5179 or Cv2.1169 antibody as a competitor. Means ± SD of duplicate values are shown. **(E)** Graphs showing the neutralization curves of SARS-CoV-2 and VOCs by Cv2.5179 IgG antibody as determined with the S-Fuse neutralization assay. Means ± SD of duplicate values are shown. $IC_{50}$ values are indicated in the top left-hand corner (in blue).

**Figure S8. Structural comparison of $V_H$1-58-encoded antibodies in complex with the SARS-CoV-2 RBD. Related to Figure 6. (A)** Superposition of the RBD-Cv2.1169 crystal structure with the complexes formed by other $V_H$1-58-encoded antibodies [S2E12 (PDB: 7R6X), COVOX-253 (PDB: 7BEN) and A23-58.1 (PDB: 7LRS)].

**Figure S9. Structural analyses of the RBD-Cv2.3235 and RBD-Cv2.6264 complexes. Related to Figure 6. (A)** Crystal structure of the complex formed by the Receptor Binding Domain (RBD) and Cv2.3235 Fab. The RBD is represented in cartoon with a transparent grey surface, highlighting the Receptor Binding Motif (RBM, yellow) and residues that are mutated in the Variants of Concern (VOCs, red). **(B)** Same as in (A) but for the RBD-Cv2.6264 complex. **(C)** Polar interactions (dashed lines) formed at the interface of the RBD-Cv2.3235 complex. For simplicity, only the interactions that involve side chains on both proteins are represented. **(D)** Same as in (C) but for the RBD-Cv2.6264 complex.

**Figure S10. Cryo-EM data collection and processing of the Cv2.1169-S$^{HexaPro}$ complex. Related to Figure 6.** A micrograph with particles, selected two-dimensional class averages, a local resolution graphic, and a scheme with the steps followed to process the collected data (along with GSFSC resolution plot) are shown for the Cv2.1169-S$^{HexaPro}$ complex. Mic., micrographs; part., particles.

**Figure S11. Cv2.1169 antibody treatment in SARS-CoV-2-infected mice and hamsters. Related to Figure 7. (A)** Dot plot comparing the SARS-CoV-2 RNA levels in the oral swabs (OS) of SARS-CoV-2-infected K18-hACE2 mice (at 4 dpi) treated with 5 mg/kg i.p. of Cv2.1169 IgG or IgA (n= 8 / group) or mGO53 control (ctr) IgA antibody (n=6) as shown in Figure 7C. Each dot corresponds to a mouse. Means of duplicate values are shown. **(B)** Dot plot comparing the human IgG concentrations in the serum of SARS-CoV-2-infected K18-hACE2 mice (at 20 dpi) receiving once 5, 10 or 20 mg/kg i.p. of antibody Cv2.1169 (n = 7 / group) as shown in Figures 7A and 7C. Each dot corresponds to a mouse. Means of duplicate values are shown. **(C)** Dot plot comparing the human IgG and IgA concentrations in the serum of K18-hACE2 mice infected with the SARS-CoV-2 β variant, and pre-treated (IgA, n=8) or treated (IgG, n=6) with Cv2.1169, or with mGO53 IgG control (ctr, n=7) as shown in Figure 7F. Each dot corresponds to a mouse. Means of duplicate values are shown. **(D)** Dot plot showing the ELISA SARS-CoV-2 tri-S binding of serum murine IgG antibodies in K18-hACE2 mice infected with the SARS-CoV-2 β variant, and pre-treated (IgA, n=8) or treated (IgG, n=6) with Cv2.1169 as shown in Figure 7F. AUC, area under the curve. Each dot corresponds to a mouse. Means of duplicate values are shown. **(E)** Dot plot comparing the human IgG and IgA concentrations in the serum of SARS-CoV-2-infected golden Syrian hamsters (at 5 dpi) treated once with Cv2.1169 or mGO53 control (5 mg/kg i.p., n=8; or 10 mg/kg i.p., n=7) as shown in Figures 7D and 7E (left and right, respectively). Each symbol corresponds to a mouse. Means of duplicate values are shown. **(F)** Dot plot showing the ELISA SARS-CoV-2 tri-S binding of serum hamster IgG antibodies in SARS-CoV-2-infected hamsters (at 5 dpi) treated once with Cv2.1169 or mGO53 control (5 mg/kg i.p., n=8; or 10 mg/kg i.p., n=7) as shown in Figures 7D and 7E (left and right, respectively). AUC, area under the curve. Each symbol corresponds to a mouse. Means of duplicate values are shown.

### Figure S1

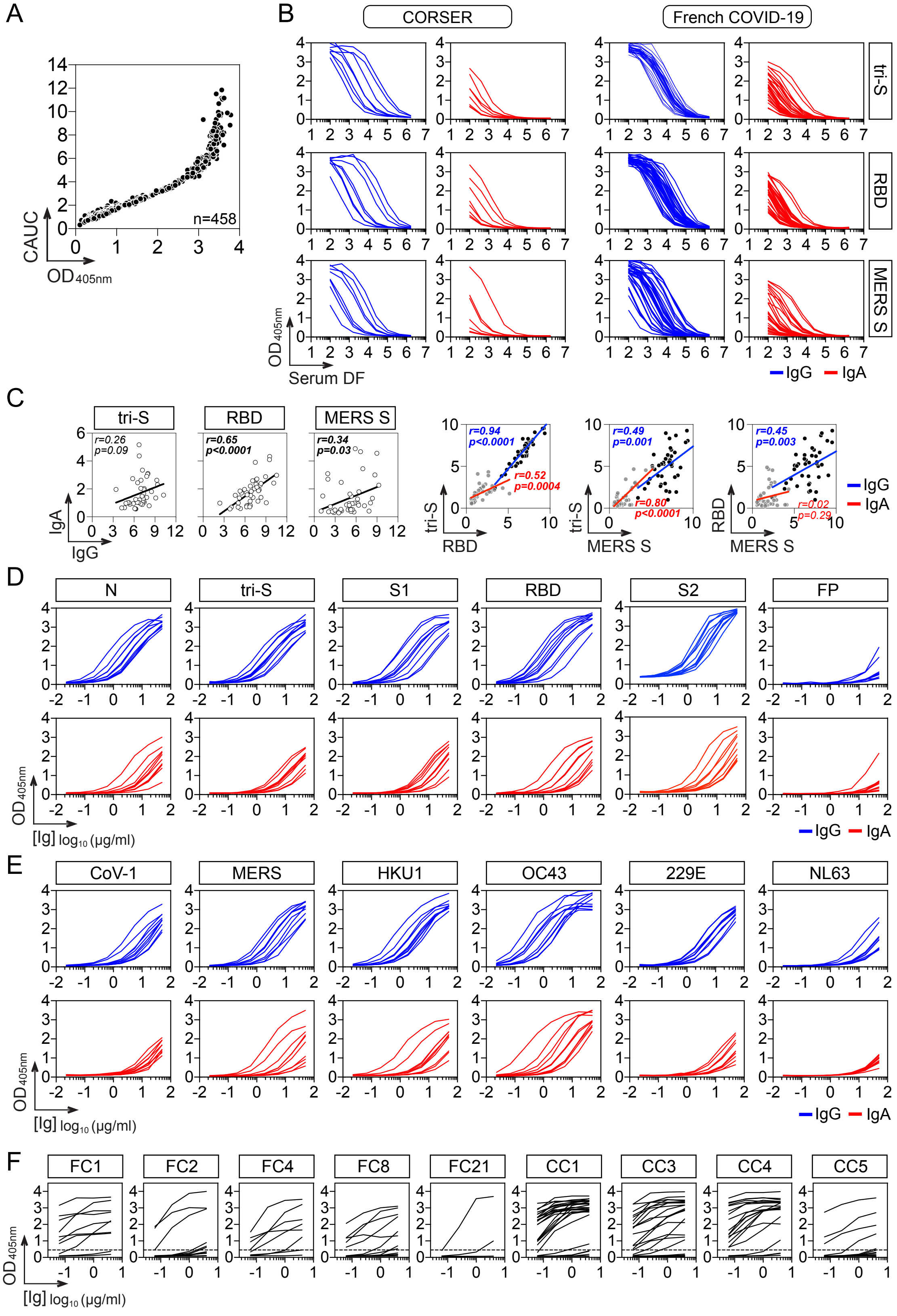

A
B
C
D
E
F
CAUC
OD405nm
n=458
CORSER
French COVID-19
tri-S
RBD
MERS S
Serum DF
IgG
IgA
r=0.26
p=0.09
r=0.65
p<0.0001
r=0.34
p=0.03
r=0.94
p<0.0001
r=0.52
p=0.0004
r=0.49
p=0.001
r=0.80
p<0.0001
r=0.45
p=0.003
r=0.02
p=0.29
N
S1
S2
FP
CoV-1
MERS
HKU1
OC43
229E
NL63
FC1
FC2
FC4
FC8
FC21
CC1
CC3
CC4
CC5
[Ig] log10 (µg/ml)

### Figure S2

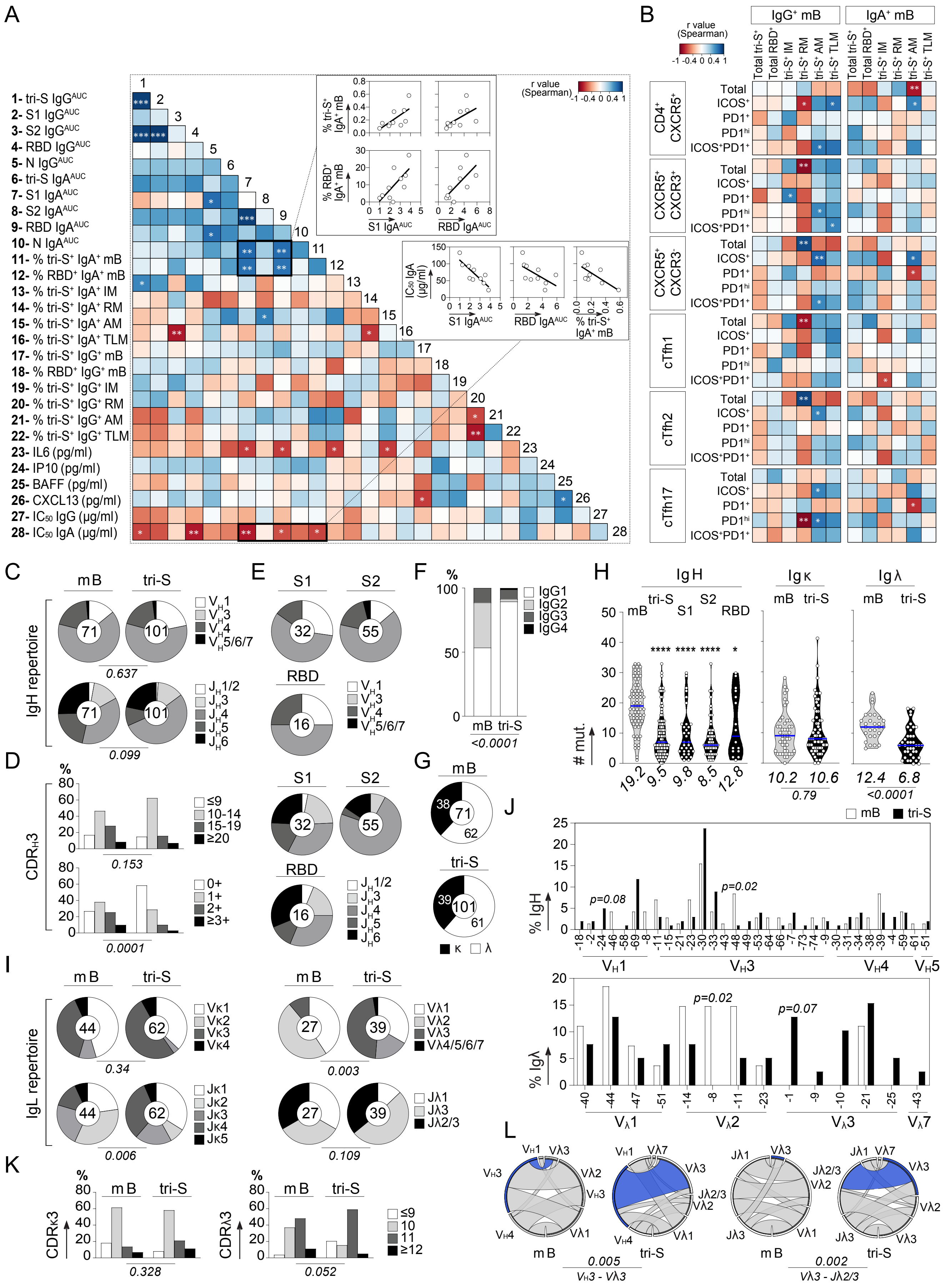

A
1- tri-S IgG^AUC
2- S1 IgG^AUC
3- S2 IgG^AUC
4- RBD IgG^AUC
5- N IgG^AUC
6- tri-S IgA^AUC
7- S1 IgA^AUC
8- S2 IgA^AUC
9- RBD IgA^AUC
10- N IgA^AUC
11- % tri-S+ IgA+ mB
12- % RBD+ IgA+ mB
13- % tri-S+ IgA+ IM
14- % tri-S+ IgA+ RM
15- % tri-S+ IgA+ AM
16- % tri-S+ IgA+ TLM
17- % tri-S+ IgG+ mB
18- % RBD+ IgG+ mB
19- % tri-S+ IgG+ IM
20- % tri-S+ IgG+ RM
21- % tri-S+ IgG+ AM
22- % tri-S+ IgG+ TLM
23- IL6 (pg/ml)
24- IP10 (pg/ml)
25- BAFF (pg/ml)
26- CXCL13 (pg/ml)
27- IC50 IgG (µg/ml)
28- IC50 IgA (µg/ml)
r value (Spearman)
S1 IgA^AUC
RBD IgA^AUC
% tri-S+ IgA+ mB
% RBD+ IgA+ mB
IC50 IgA (µg/ml)
B
IgG+ mB
IgA+ mB
Total tri-S+
Total RBD+
tri-S+ IM
tri-S+ RM
tri-S+ AM
tri-S+ TLM
CD4+ CXCR5+
CXCR5+ CXCR3+
CXCR5+ CXCR3-
cTfh1
cTfh2
cTfh17
Total
ICOS+
PD1+
PD1hi
ICOS+PD1+
C
mB
tri-S
IgH repertoire
0.637
0.099
D
CDRH3
0.153
0.0001
E
S1
S2
RBD
F
IgG1
IgG2
IgG3
IgG4
<0.0001
G
κ
λ
H
IgH
Igκ
Igλ
RBD
### mut.
19.2
9.5
9.8
8.5
12.8
10.2
10.6
0.79
12.4
6.8
<0.0001
J
% IgH
p=0.08
p=0.02
VH1
VH3
VH4
VH5
% Igλ
p=0.02
p=0.07
Vλ1
Vλ2
Vλ3
Vλ7
I
IgL repertoire
0.34
0.003
0.006
0.109
K
CDRκ3
CDRλ3
0.328
0.052
L
0.005
VH3 - Vλ3
0.002
Vλ3 - Jλ2/3

### Figure S3

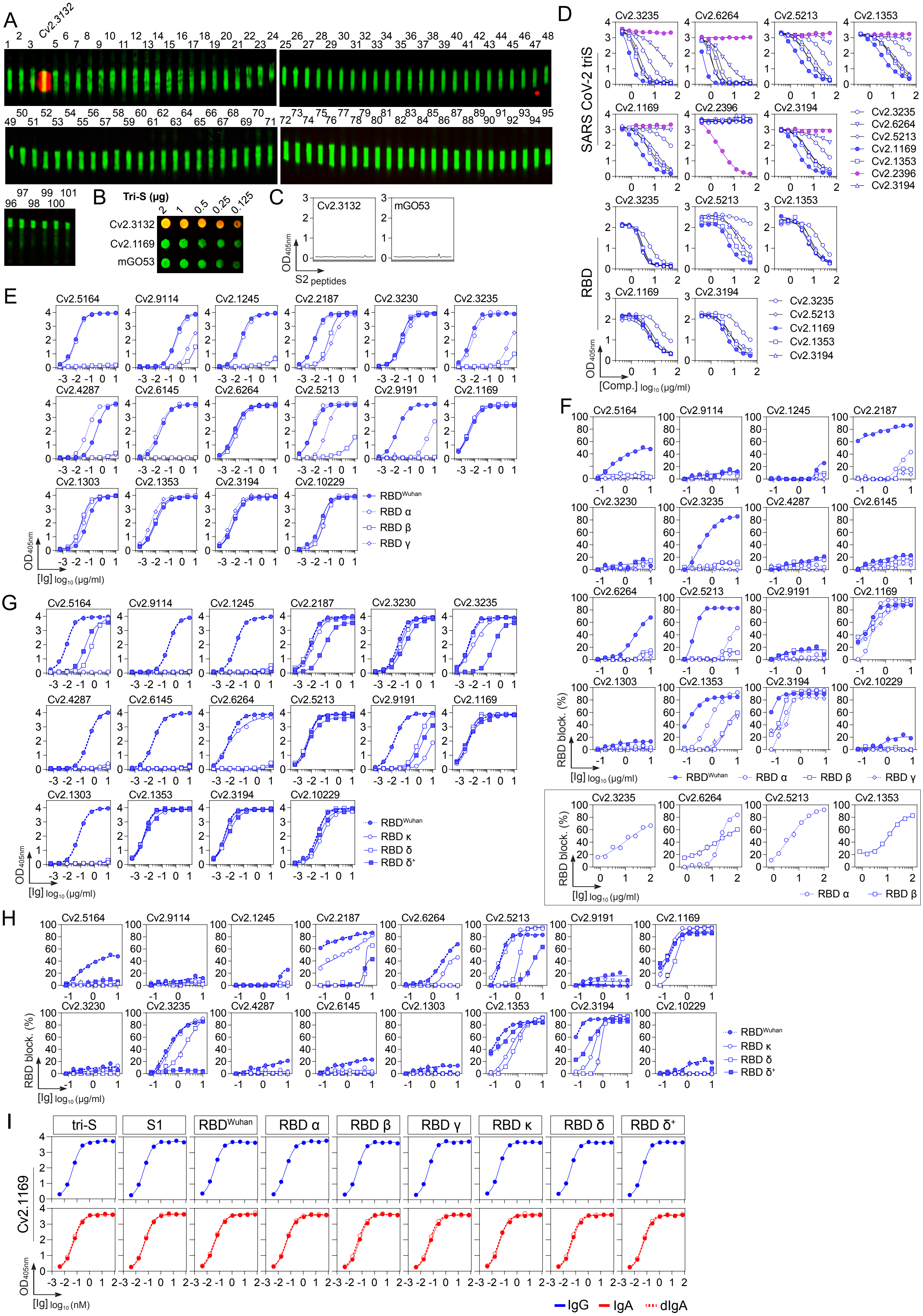

A
Cv2.3132
B
Tri-S (µg)
2 1 0.5 0.25 0.125
Cv2.3132
Cv2.1169
mGO53
C
Cv2.3132
mGO53
OD405nm
S2 peptides
D
SARS CoV-2 triS
RBD
Cv2.3235
Cv2.6264
Cv2.5213
Cv2.1353
Cv2.1169
Cv2.2396
Cv2.3194
OD405nm
[Comp.] log10 (µg/ml)
E
Cv2.5164
Cv2.9114
Cv2.1245
Cv2.2187
Cv2.3230
Cv2.3235
Cv2.4287
Cv2.6145
Cv2.6264
Cv2.5213
Cv2.9191
Cv2.1169
Cv2.1303
Cv2.1353
Cv2.3194
Cv2.10229
RBDWuhan
RBD α
RBD β
RBD γ
OD405nm
[Ig] log10 (µg/ml)
F
RBD block. (%)
[Ig] log10 (µg/ml)
RBDWuhan
RBD α
RBD β
RBD γ
G
RBDWuhan
RBD κ
RBD δ
RBD δ+
OD405nm
[Ig] log10 (µg/ml)
H
RBD block. (%)
[Ig] log10 (µg/ml)
I
tri-S
S1
RBDWuhan
RBD α
RBD β
RBD γ
RBD κ
RBD δ
RBD δ+
Cv2.1169
OD405nm
[Ig] log10 (nM)
IgG
IgA
dIgA

### Figure S4

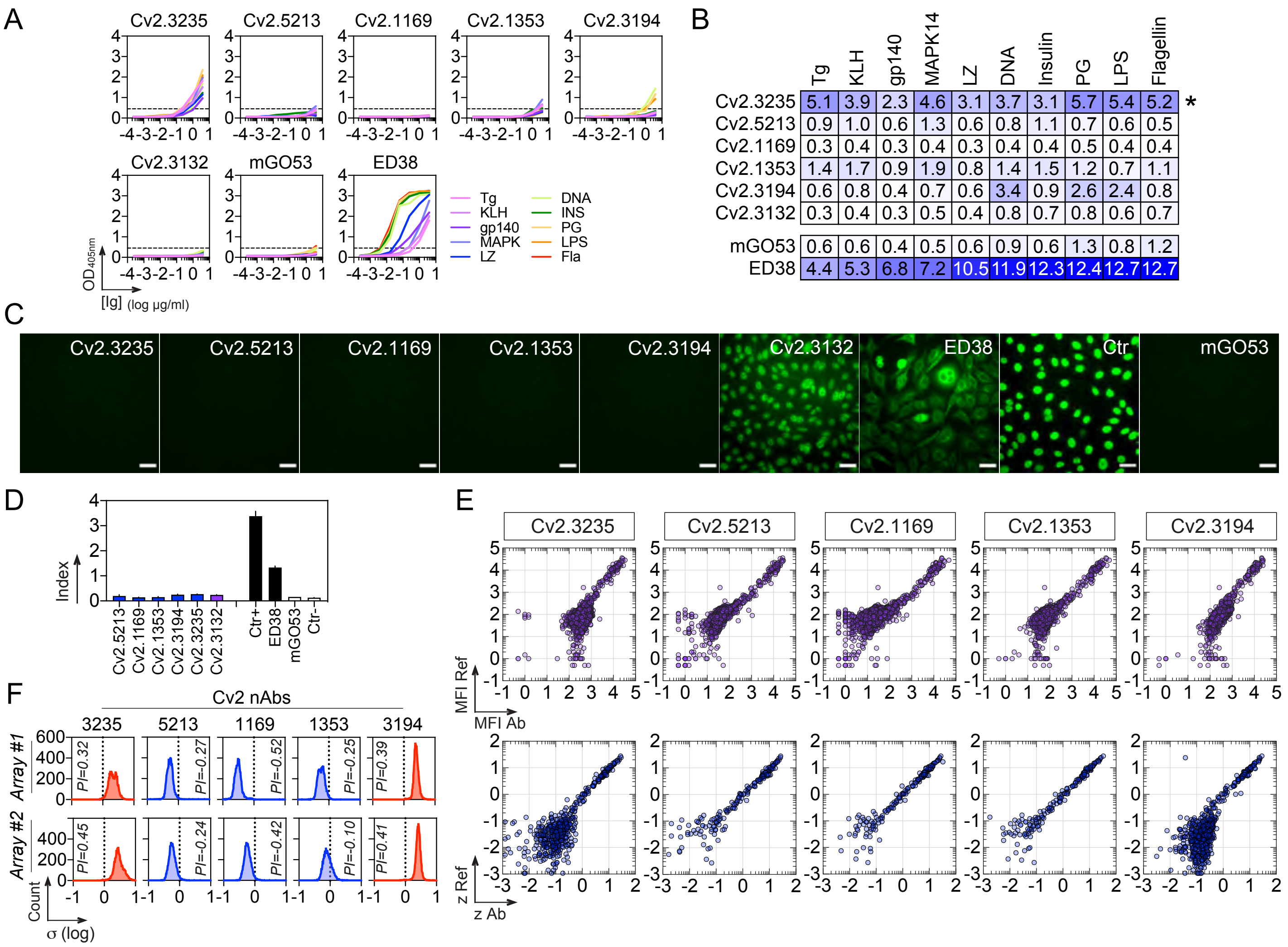


| | Tg | KLH | gp140 | MAPK14 | LZ | DNA | Insulin | PG | LPS | Flagellin | |
|---|---|---|---|---|---|---|---|---|---|---|---|
| Cv2.3235 | 5.1 | 3.9 | 2.3 | 4.6 | 3.1 | 3.7 | 3.1 | 5.7 | 5.4 | 5.2 | * |
| Cv2.5213 | 0.9 | 1.0 | 0.6 | 1.3 | 0.6 | 0.8 | 1.1 | 0.7 | 0.6 | 0.5 | |
| Cv2.1169 | 0.3 | 0.4 | 0.3 | 0.4 | 0.3 | 0.4 | 0.4 | 0.5 | 0.4 | 0.4 | |
| Cv2.1353 | 1.4 | 1.7 | 0.9 | 1.9 | 0.8 | 1.4 | 1.5 | 1.2 | 0.7 | 1.1 | |
| Cv2.3194 | 0.6 | 0.8 | 0.4 | 0.7 | 0.6 | 3.4 | 0.9 | 2.6 | 2.4 | 0.8 | |
| Cv2.3132 | 0.3 | 0.4 | 0.3 | 0.5 | 0.4 | 0.8 | 0.7 | 0.8 | 0.6 | 0.7 | |
| mGO53 | 0.6 | 0.6 | 0.4 | 0.5 | 0.6 | 0.9 | 0.6 | 1.3 | 0.8 | 1.2 | |
| ED38 | 4.4 | 5.3 | 6.8 | 7.2 | 10.5 | 11.9 | 12.3 | 12.4 | 12.7 | 12.7 | |

### Figure S5

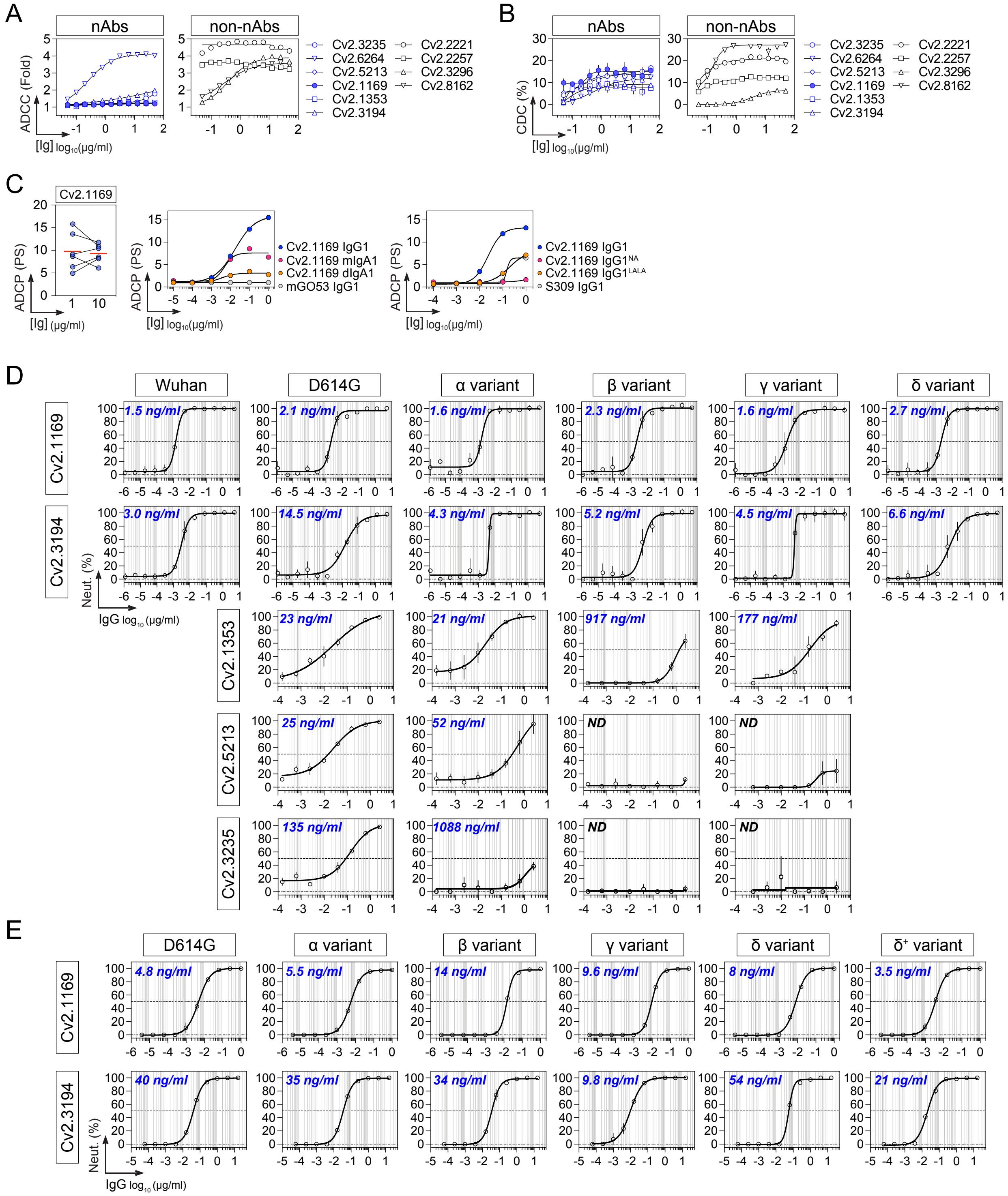
A
nAbs
non-nAbs
ADCC (Fold)
[Ig] log10(µg/ml)
Cv2.3235
Cv2.6264
Cv2.5213
Cv2.1169
Cv2.1353
Cv2.3194
Cv2.2221
Cv2.2257
Cv2.3296
Cv2.8162
B
CDC (%)
C
Cv2.1169
ADCP (PS)
[Ig] (µg/ml)
Cv2.1169 IgG1
Cv2.1169 mIgA1
Cv2.1169 dIgA1
mGO53 IgG1
Cv2.1169 IgG1NA
Cv2.1169 IgG1LALA
S309 IgG1
D
Wuhan
D614G
α variant
β variant
γ variant
δ variant
1.5 ng/ml
2.1 ng/ml
1.6 ng/ml
2.3 ng/ml
1.6 ng/ml
2.7 ng/ml
3.0 ng/ml
14.5 ng/ml
4.3 ng/ml
5.2 ng/ml
4.5 ng/ml
6.6 ng/ml
Neut. (%)
IgG log10 (µg/ml)
Cv2.1353
23 ng/ml
21 ng/ml
917 ng/ml
177 ng/ml
Cv2.5213
25 ng/ml
52 ng/ml
ND
ND
Cv2.3235
135 ng/ml
1088 ng/ml
ND
ND
E
D614G
α variant
β variant
γ variant
δ variant
δ+ variant
4.8 ng/ml
5.5 ng/ml
14 ng/ml
9.6 ng/ml
8 ng/ml
3.5 ng/ml
40 ng/ml
35 ng/ml
34 ng/ml
9.8 ng/ml
54 ng/ml
21 ng/ml

### Figure S6

A

| | Cv2.1169 | Cv2.3194 | REGN10987 | COV2-2196 | CT-P59 | S309 | COV2-2130 | REGN10933 | ADG-2 | LY-CoV555 | CB6 |
|---|---|---|---|---|---|---|---|---|---|---|---|
| tri-S | 3 | 3 | 3 | 3 | 3 | 3 | 3 | 3 | 3 | 3 | 3 |
| S1 | 3 | 3 | 3 | 3 | 3 | 3 | 3 | 3 | 3 | 3 | 3 |
| Wuhan | 3 | 3 | 3 | 3 | 3 | 3 | 3 | 3 | 3 | 3 | 3 |
| α | 3 | 3 | 3 | 3 | 3 | 3 | 3 | 3 | 3 | 3 | 3 |
| β | 3 | 4 | 3 | 3 | 3 | 3 | 3 | 3 | 3 | | |
| γ | 4 | 3 | 4 | 4 | 4 | 4 | 3 | 3 | 3 | | |
| δ | 3 | 3 | 3 | 3 | 3 | 3 | 3 | 3 | 3 | 3 | 3 |
| δ⁺ | 4 | 3 | 4 | 3 | 3 | 3 | 3 | 3 | 3 | 3 | |
| κ | 4 | 4 | 4 | 4 | 3 | 3 | 4 | 4 | 4 | | 4 |

B

| | Cv2.1169 | Cv2.3194 | ADG-2 | COV2-2130 | REGN10987 | COV2-2196 | CT-P59 | REGN10933 | CB6 | LY-CoV555 | S309 | COV2-2130 +COV2-2196 | REGN10933 +REGN10987 | LY-CoV555 +CB6 |
|---|---|---|---|---|---|---|---|---|---|---|---|---|---|---|
| tri-S | 96 | 97 | 96 | 95 | 93 | 96 | 95 | 96 | 88 | 94 | 5 | NT | 97 | 96 |
| Wuhan | 94 | 96 | 94 | 96 | 91 | 97 | 96 | 97 | 92 | 94 | | 95 | 96 | 93 |
| α | 86 | 97 | 92 | 85 | 51 | 83 | 86 | 91 | 9 | 82 | 5 | 96 | 98 | 67 |
| β | 92 | 95 | 95 | 96 | 71 | 58 | 49 | 28 | 2 | 1 | | 95 | 95 | |
| γ | 86 | 94 | 89 | 90 | 53 | 44 | 19 | | | | | 94 | 90 | 1 |
| δ | 93 | 94 | 96 | 77 | 89 | 92 | 91 | 97 | 89 | 20 | | 95 | 97 | 81 |
| δ⁺ | 95 | 94 | 80 | 88 | 93 | 96 | 91 | 93 | 3 | 4 | | 95 | 96 | 4 |
| κ | 95 | 92 | 95 | 80 | 87 | 91 | 94 | 94 | 89 | 17 | | 94 | 97 | 71 |

C

| | Cv2.1169 | REGN10987 | COV2-2196 | ADG-2 | COV2-2130 | S309 | CT-P59 | REGN10933 | CB6 | LY-CoV555 | REGN10933 +REGN10987 | COV2-2130 +COV2-2196 | LY-CoV555 +CB6 |
|---|---|---|---|---|---|---|---|---|---|---|---|---|---|
| D614G | 0.06 | 0.16 | 0.27 | 0.60 | 0.11 | 1.44 | 0.17 | 0.08 | 0.59 | 1.62 | 0.07 | 0.23 | 0.92 |
| α | 0.04 | 0.03 | 0.08 | 0.08 | 0.10 | 1.30 | 0.33 | 0.06 | 2.06 | 0.07 | 0.04 | 0.05 | 0.20 |
| β | 0.10 | 0.04 | 0.11 | 0.15 | 0.28 | 1.43 | 2.10 | 1.81 | | | 0.14 | 0.28 | |
| γ | 0.36 | 0.09 | 0.71 | 0.67 | 0.62 | 1.23 | 1.93 | 4.12 | | | 0.32 | 0.27 | |
| δ | 0.63 | 0.90 | 0.64 | 1.09 | 2.88 | 1.45 | 2.16 | 0.45 | 2.48 | | 0.68 | 0.76 | 2.18 |

D

*Competitors* ↓ / *Biot. mAbs* →

tri-S Wuhan

| | CT-P59 | Cv2.1169 | Cv2.3194 | COV2-2196 | REGN10933 | CB6 | LY-CoV555 | COV2-2130 | REGN10987 | S309 | ADG-2 |
|---|---|---|---|---|---|---|---|---|---|---|---|
| CT-P59 | 91 | 98 | 96 | 97 | 98 | 98 | 90 | 97 | 6 | | 93 |
| Cv2.1169 | 42 | 93 | 90 | 95 | 94 | 97 | 83 | | | | 2 |
| Cv2.3194 | 50 | 89 | 90 | 87 | 91 | 97 | 85 | | | | 81 |
| COV2-2196 | 35 | 93 | 89 | 94 | 94 | 97 | 80 | | | | 5 |
| REGN10933 | 30 | 86 | 84 | 91 | 94 | 97 | 69 | | | | 25 |
| CB6 | 2 | 86 | 35 | 28 | 38 | 93 | 11 | | | | 37 |
| LY-CoV555 | 47 | 92 | 84 | 94 | 94 | 98 | 91 | 95 | 29 | 1 | |
| COV2-2130 | 20 | 2 | | 2 | | | 63 | 93 | 96 | 81 | |
| REGN10987 | | | | | | | 1 | 77 | 94 | 68 | 47 |
| S309 | | | | | | 4 | | 85 | 94 | 90 | 1 |
| ADG-2 | 37 | 2 | 71 | 3 | 79 | 96 | | | 95 | 1 | 93 |

RBD Wuhan

| | CT-P59 | Cv2.1169 | Cv2.3194 | COV2-2196 | REGN10933 | CB6 | LY-CoV555 | COV2-2130 | REGN10987 | S309 | ADG-2 |
|---|---|---|---|---|---|---|---|---|---|---|---|
| CT-P59 | 79 | 94 | 91 | 95 | 93 | 95 | 78 | 93 | | 10 | 65 |
| Cv2.1169 | 38 | 90 | 91 | 92 | 89 | 96 | 60 | | | 10 | |
| Cv2.3194 | 30 | 82 | 88 | 85 | 86 | 94 | 25 | | | 10 | 42 |
| COV2-2196 | 3 | 82 | 82 | 88 | 87 | 93 | 52 | | | 7 | |
| REGN10933 | 21 | 64 | 84 | 81 | 87 | 95 | 33 | | 2 | 9 | 1 |
| CB6 | 2 | 16 | 67 | 15 | 27 | 84 | 11 | | | 16 | 11 |
| LY-CoV555 | 40 | 65 | 76 | 81 | 71 | 93 | 86 | 88 | 4 | 12 | |
| COV2-2130 | | 3 | 11 | 4 | | | 36 | 87 | 88 | 73 | |
| REGN10987 | 22 | 12 | 8 | 4 | 1 | | 15 | 77 | 86 | 80 | 13 |
| S309 | | 6 | 13 | | | | | 34 | 16 | 88 | |
| ADG-2 | | 20 | 60 | 1 | 6 | 77 | 15 | 1 | 72 | 19 | 84 |

RBD δ

| | Cv2.1169 | Cv2.3194 | COV2-2196 | REGN10933 | CB6 | CT-P59 | LY-CoV555 | COV2-2130 | REGN10987 | S309 | ADG-2 |
|---|---|---|---|---|---|---|---|---|---|---|---|
| Cv2.1169 | 86 | 83 | 91 | 67 | 92 | 90 | 92 | | | 6 | |
| Cv2.3194 | 85 | 87 | 88 | 81 | 91 | 90 | 92 | | | | 41 |
| COV2-2196 | 77 | 79 | 87 | 54 | 90 | 88 | 91 | | | 2 | |
| REGN10933 | 88 | 83 | 93 | 86 | 93 | 93 | 92 | 12 | | | |
| CB6 | 18 | 43 | 37 | 13 | 81 | 62 | 91 | | | 9 | 6 |
| CT-P59 | 37 | 68 | 49 | 18 | 80 | 87 | 90 | 75 | | | 1 |
| LY-CoV555 | | | | | | | 79 | 4 | | | |
| COV2-2130 | 4 | | 4 | | | 23 | 91 | 88 | 55 | 47 | |
| REGN10987 | 1 | | 4 | | | 4 | 34 | 90 | 86 | 64 | 22 |
| S309 | | | | | | | 2 | 67 | 27 | 82 | 5 |
| ADG-2 | | 33 | | | 82 | 86 | 6 | | 82 | 23 | 85 |

### Figure S7

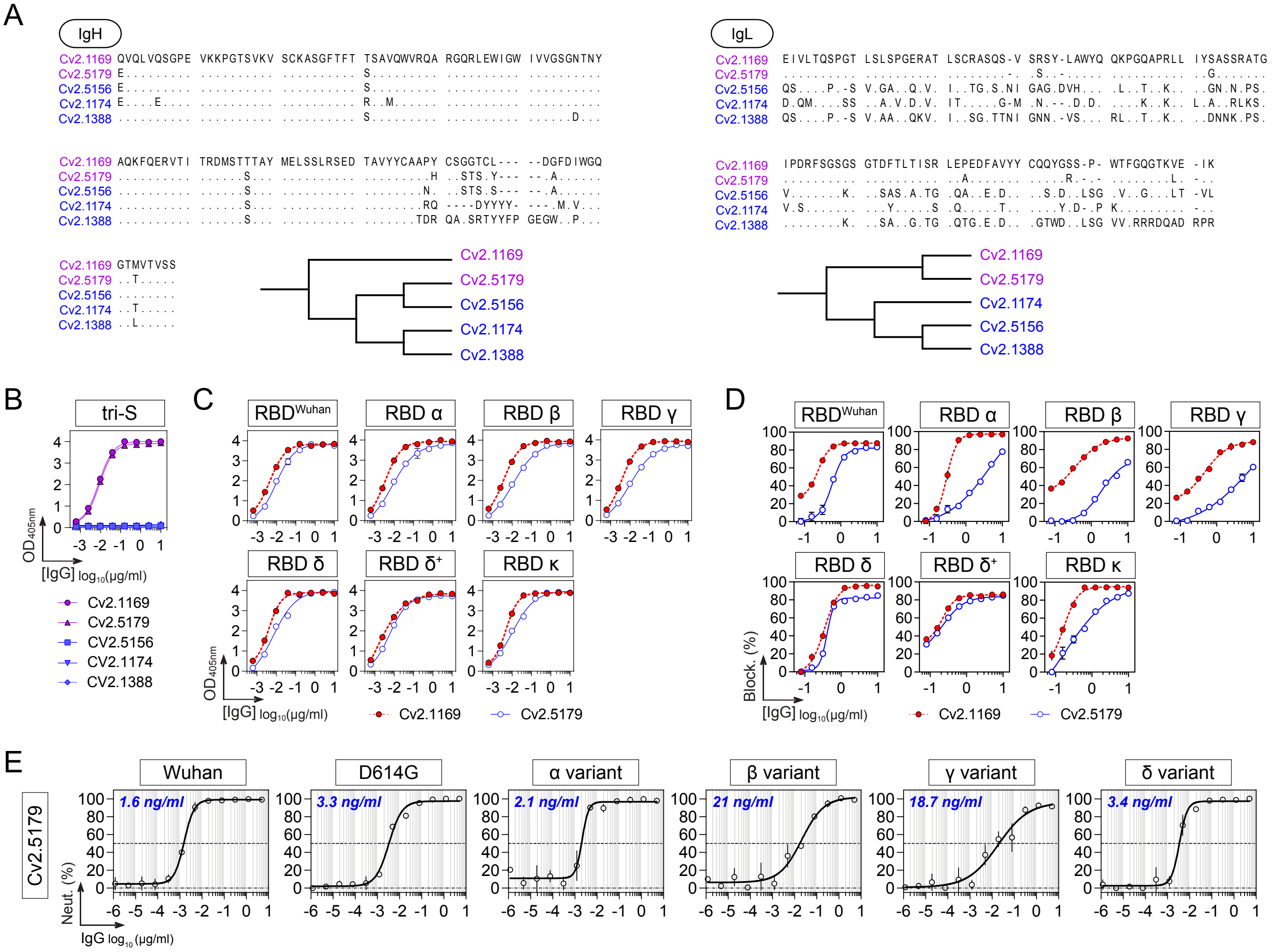

A
IgH
Cv2.1169 QVQLVQSGPE VKKPGTSVKV SCKASGFTFT TSAVQWVRQA RGQRLEWIGW IVVGSGNTNY
Cv2.5179 E......... .......... .......... S......... .......... ..........
Cv2.5156 E......... .......... .......... S......... .......... ..........
Cv2.1174 E....E.... .......... .......... R..M...... .......... ..........
Cv2.1388 .......... .......... .......... S......... .......... ......D...
Cv2.1169 AQKFQERVTI TRDMSTTTAY MELSSLRSED TAVYYCAAPY CSGGTCL--- --DGFDIWGQ
Cv2.5179 .......... ......S... .......... .........H ..STS.Y--- --.A......
Cv2.5156 .......... ......S... .......... ........N. ..STS.S--- --.A......
Cv2.1174 .......... ......S... .......... ........RQ ----DYYYY- ---.M.V...
Cv2.1388 .......... ......S... .......... .......TDR QA.SRTYYFP GEGW..P...
Cv2.1169 GTMVTVSS
Cv2.5179 ..T.....
Cv2.5156 ........
Cv2.1174 ..T.....
Cv2.1388 ..L.....
Cv2.1169
Cv2.5179
Cv2.5156
Cv2.1174
Cv2.1388
IgL
Cv2.1169 EIVLTQSPGT LSLSPGERAT LSCRASQS-V SRSY-LAWYQ QKPGQAPRLL IYSASSRATG
Cv2.5179 .......... .......... ........-. .S..-..... .......... ..G.......
Cv2.5156 QS....P.-S V.GA..Q.V. I..TG.S.NI GAG.DVH... .L..T..K.. ..GN.N.PS.
Cv2.1174 D.QM....SS ..A.V.D.V. IT.....G-M .N.--.D.D. ....K..K.. L.A..RLKS.
Cv2.1388 QS....P.-S V.AA..QKV. I..SG.TTNI GNN.-VS... RL..T..K.. ..DNNK.PS.
Cv2.1169 IPDRFSGSGS GTDFTLTISR LEPEDFAVYY CQQYGSS-P- WTFGQGTKVE -IK
Cv2.5179 .......... .......... ..A....... .....R.-.- ........L. -..
Cv2.5156 V.......K. ..SAS.A.TG .QA..E.D.. ..S.D..LSG .V..G...LT -VL
Cv2.1174 V.S....... ...Y.....S .Q.....T.. ....Y.D-.P K......... -..
Cv2.1388 ........K. ..SA..G.TG .QTG.E.D.. .GTWD..LSG VV.RRRDQAD RPR
Cv2.1169
Cv2.5179
Cv2.1174
Cv2.5156
Cv2.1388
B
tri-S
OD 405nm
[IgG] log10(µg/ml)
Cv2.1169
Cv2.5179
CV2.5156
CV2.1174
CV2.1388
C
RBD Wuhan
RBD α
RBD β
RBD γ
RBD δ
RBD δ+
RBD κ
OD 405nm
[IgG] log10(µg/ml)
Cv2.1169
Cv2.5179
D
RBD Wuhan
RBD α
RBD β
RBD γ
RBD δ
RBD δ+
RBD κ
Block. (%)
[IgG] log10(µg/ml)
Cv2.1169
Cv2.5179
E
Cv2.5179
Wuhan
1.6 ng/ml
D614G
3.3 ng/ml
α variant
2.1 ng/ml
β variant
21 ng/ml
γ variant
18.7 ng/ml
δ variant
3.4 ng/ml
Neut. (%)
IgG log10 (µg/ml)

### Figure S8

A

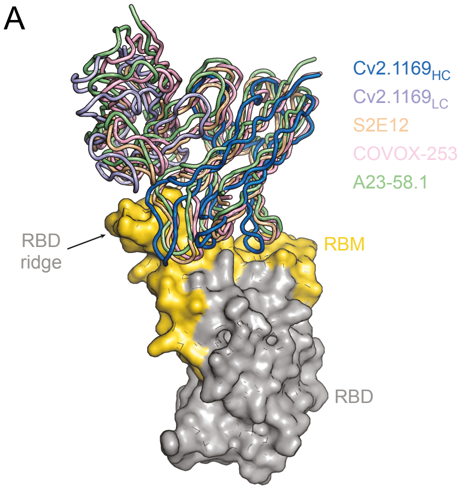
Cv2.1169HC
Cv2.1169LC
S2E12
COVOX-253
A23-58.1
RBD
ridge
RBM
RBD

### Figure S9

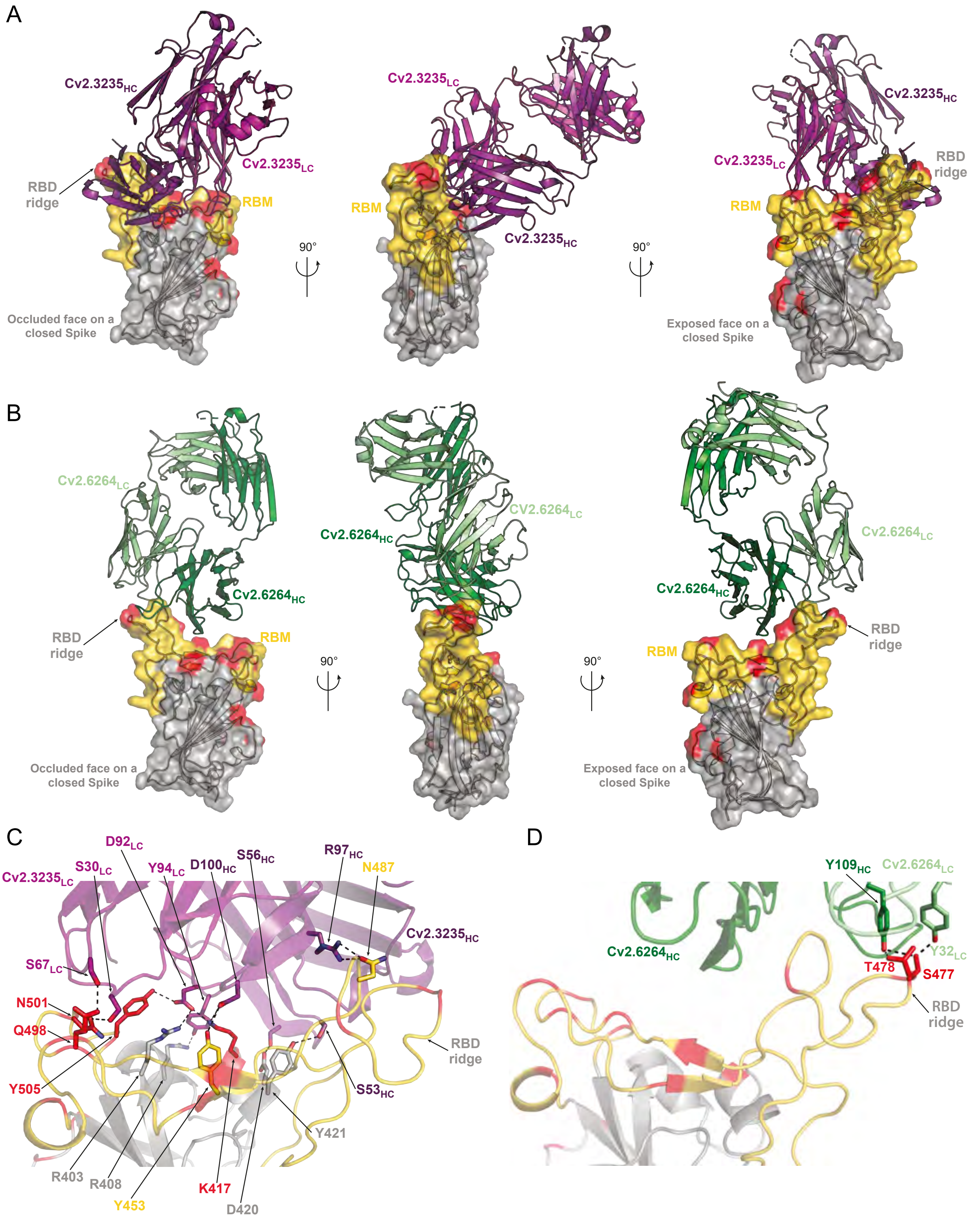

A
Cv2.3235HC
Cv2.3235LC
RBD ridge
RBM
Occluded face on a closed Spike
90°
Cv2.3235LC
RBM
Cv2.3235HC
90°
Cv2.3235HC
Cv2.3235LC
RBD ridge
RBM
Exposed face on a closed Spike
B
Cv2.6264LC
Cv2.6264HC
RBD ridge
RBM
Occluded face on a closed Spike
90°
Cv2.6264HC
Cv2.6264LC
90°
Cv2.6264LC
Cv2.6264HC
RBM
RBD ridge
Exposed face on a closed Spike
C
Cv2.3235LC
S30LC
D92LC
Y94LC
D100HC
S56HC
R97HC
N487
Cv2.3235HC
S67LC
N501
Q498
Y505
RBD ridge
S53HC
Y421
R403
R408
Y453
K417
D420
D
Y109HC
Cv2.6264LC
Cv2.6264HC
Y32LC
T478
S477
RBD ridge

#### Figure S10

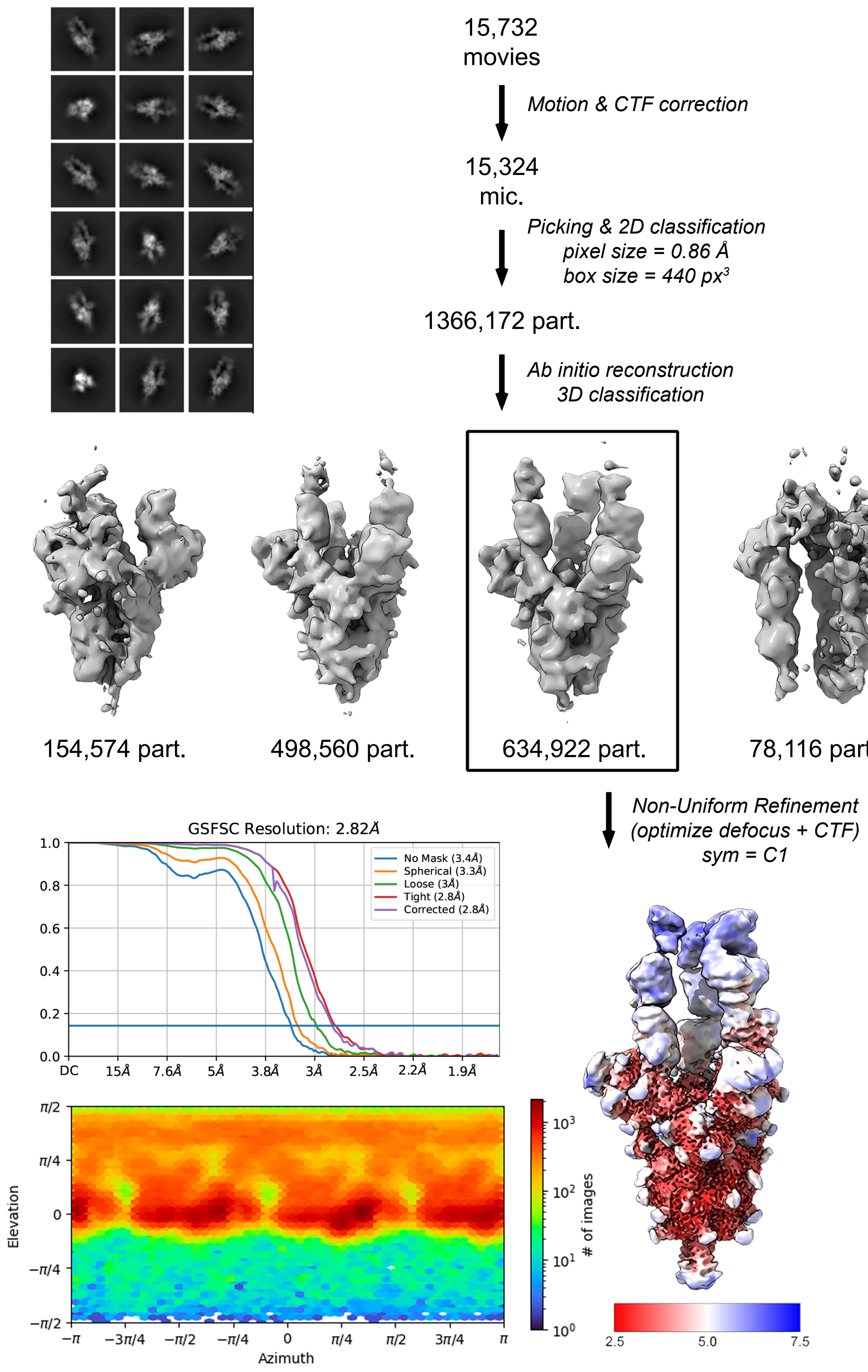

15,732
movies
Motion & CTF correction
15,324
mic.
Picking & 2D classification
pixel size = 0.86 Å
box size = 440 px³
1366,172 part.
Ab initio reconstruction
3D classification
154,574 part.
498,560 part.
634,922 part.
78,116 part.
Non-Uniform Refinement
(optimize defocus + CTF)
sym = C1
GSFSC Resolution: 2.82Å
No Mask (3.4Å)
Spherical (3.3Å)
Loose (3Å)
Tight (2.8Å)
Corrected (2.8Å)
DC
15Å
7.6Å
5Å
3.8Å
3Å
2.5Å
2.2Å
1.9Å
Elevation
Azimuth
### of images
2.5
5.0
7.5

### Figure S11

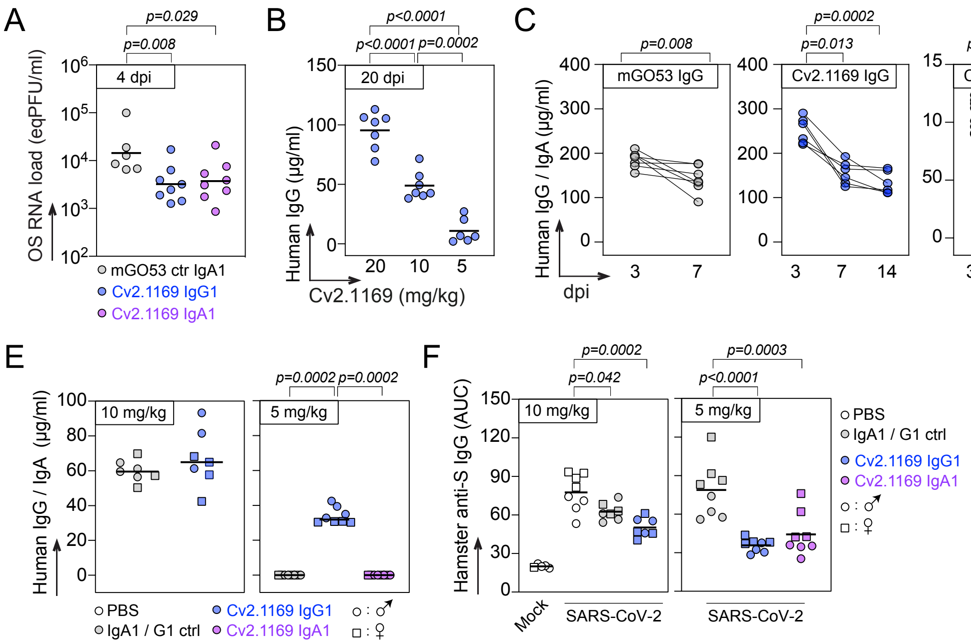
A
p=0.029
p=0.008
4 dpi
OS RNA load (eqPFU/ml)
mGO53 ctr IgA1
Cv2.1169 IgG1
Cv2.1169 IgA1
B
p<0.0001
p<0.0001
p=0.0002
20 dpi
Human IgG (µg/ml)
Cv2.1169 (mg/kg)
C
p=0.008
mGO53 IgG
p=0.0002
p=0.013
Cv2.1169 IgG
p<0.0001
p<0.0001
Cv2.1169 IgA
Human IgG / IgA (µg/ml)
dpi
D
p=0.002
14 dpi
Mouse anti-S IgG (AUC)
Cv2.1169 IgG1
Cv2.1169 IgA1
E
10 mg/kg
p=0.0002
p=0.0002
5 mg/kg
Human IgG / IgA (µg/ml)
PBS
IgA1 / G1 ctrl
Cv2.1169 IgG1
Cv2.1169 IgA1
F
p=0.0002
p=0.042
p=0.0003
p<0.0001
10 mg/kg
5 mg/kg
Hamster anti-S IgG (AUC)
Mock
SARS-CoV-2
SARS-CoV-2
PBS
IgA1 / G1 ctrl
Cv2.1169 IgG1
Cv2.1169 IgA1

**Table S1. Immunoglobulin gene repertoire, reactivity and antiviral functions of human SARS-CoV-2 spike memory B-cell antibodies. Related to Figures 1-3.**

| | | IgH | | | | | | | | | IgL | | | | | | | S-Flow | | | | Reactivity ($OD_{450nm}$) | | | | | | | | | | | | | | | | | |
|---|---|---|---|---|---|---|---|---|---|---|---|---|---|---|---|---|---|---|---|---|---|---|---|---|---|---|---|---|---|---|---|---|---|---|---|---|---|---|---|
| | | | | | | $CDR_H3$ | | | | | | | | $CDR_L3$ | | | | | | | | | | | | | | | | | | | | | | | | | |
| mAb | Donnor | $V_H$ | $D_H$ | $J_H$ | Subtype | Sequence (AA) | Length | (+) | (-) | $V_H$ MUT | IgL | $V_L$ | $J_L$ | Sequence (AA) | Length | $V_L$ MUT | Target | Pos. Cells (%) | log10MFI | tri-S (AUC) | Capt. tri-S | tri-S | S1 | RBD | NTD | CD | S2 | FP | SARS-CoV-1 | MERS-CoV | HKU1 | OC43 | 229E | tri-S Block. (%) | PseudoN (%) | S-Fuse (%) | ADCC (Fold) | CDC (%) | ADCP (PS) |
| **Cv2.1102** | CC1 | 3-33 | 3-10 | 4 | γ1 | DGGFGEANFDY | 11 | 0 | 3 | 18 | λ | 3-9 | 1 | QVWDSSPYV | 9 | 5 | S2 | 94.9 | 4.7 | 7.4 | 2.70 | 2.58 | 0.15 | 0.17 | 0.09 | 0.16 | 2.63 | 0.00 | 0.51 | 0.09 | 0.16 | 0.09 | 0.20 | 5 | 12 | 12 | 2.7 | 3.1 | 3.6 |
| **Cv2.1150** | CC1 | 4-31 | 2-2/2-2 | 4 | γ1 | TLGYCSSTSCYVDY | 14 | 0 | 1 | 10 | κ | 1-5 | 1 | QQYNTYFQT | 9 | 19 | S2 | 94.6 | 4.5 | 10.7 | 0.94 | 3.31 | 0.12 | 0.08 | 0.08 | 0.16 | 3.02 | 0.00 | 1.94 | 0.13 | 0.12 | 0.16 | 0.17 | 2 | 5 | 15 | 2.0 | 2.6 | 1.6 |
| **Cv2.1151** | CC1 | 1-69/1-69D | 6-19 | 4 | γ1 | DPSGWYLGYFDY | 12 | 0 | 2 | 9 | κ | 3-20 | 1 | QQYGSSPPT | 9 | 6 | S2 | 96.0 | 4.8 | 11.2 | 2.66 | 3.31 | 0.08 | 0.08 | 0.06 | 0.08 | 3.19 | 0.01 | 3.11 | 0.06 | 0.08 | 0.07 | 0.08 | 0 | 0 | 11 | 2.3 | 3.0 | 5.2 |
| **Cv2.1153** | CC1 | 3-7 | 3-3 | 6 | γ1 | VALSGVIRLLDRGYYYYGMDV | 21 | 2 | 2 | 4 | κ | 1-39/1D-39 | 3 | QQSYTTPHT | 9 | 10 | NTD | 96.9 | 4.6 | 11.1 | 0.84 | 3.40 | 2.80 | 0.10 | 3.24 | 0.15 | 0.13 | 0.00 | 0.16 | 0.16 | 0.11 | 0.14 | 0.17 | 8 | 22 | 60 | 2.7 | 11.9 | 3.8 |
| **Cv2.1169** | CC1 | 1-58 | 2-15 | 3 | α1 | PYCSGGTCLDGFDI | 14 | 0 | 2 | 4 | κ | 3-20 | 1 | QQYGSSPWT | 9 | 5 | RBD | 95.8 | 5.3 | 13.1 | 2.23 | 3.78 | 3.52 | 3.51 | 0.09 | 0.08 | 0.08 | 0.06 | 0.08 | 0.06 | 0.07 | 0.15 | 0.07 | 90 | 100 | 95 | 1.2 | 14.6 | 6.2 |
| **Cv2.1177** | CC1 | 1-24 | 3-3 | 4 | γ1 | QYAIFGVGYLDY | 12 | 0 | 1 | 13 | κ | 4-1 | 4 | QQYYTTPLT | 9 | 6 | NTD | 98.4 | 5.5 | 13.3 | 1.70 | 3.83 | 3.47 | 0.08 | 3.31 | 0.11 | 0.11 | 0.01 | 0.12 | 0.11 | 0.10 | 0.12 | 0.12 | 3 | 0 | 68 | 3.1 | 14.0 | 4.0 |
| **Cv2.1191** | CC1 | 3-23 | 3-22 | 4 | γ1 | GPMIKVVITWEGEGYFFDY | 19 | 1 | 3 | 13 | κ | 3-20 | 4 | QQYGSSPLT | 9 | 7 | NTD | 96.8 | 4.5 | 15.7 | 1.93 | 3.57 | 3.49 | 0.08 | 3.28 | 0.08 | 0.07 | 0.01 | 0.08 | 0.07 | 0.08 | 0.07 | 0.09 | 53 | 32 | 36 | 1.1 | 0.9 | 2.5 |
| **Cv2.1223** | CC1 | 1-69/1-69D | 3-9 | 4 | γ1 | MYGYDNIPLLDY | 12 | 0 | 2 | 7 | κ | 3-11 | 4 | QQRSNWPPKLT | 11 | 5 | S2 | 97.4 | 4.9 | 11.3 | 2.59 | 3.32 | 0.12 | 0.08 | 0.10 | 0.13 | 3.25 | 0.01 | 3.24 | 0.18 | 0.13 | 0.14 | 0.17 | 0 | 3 | 15 | 1.4 | 0.8 | 5.4 |
| **Cv2.1245** | CC1 | 4-34 | 3-22 | 4 | γ1 | VNSNYYDSSGSRLGHYFDY | 19 | 2 | 2 | 22 | κ | 3-15 | 1 | QQYNNWRKT | 9 | 14 | RBD | 96.1 | 4.6 | 0.6 | 0.08 | 0.53 | 1.72 | 3.86 | 0.10 | 0.18 | 0.14 | 0.01 | 0.15 | 0.10 | 0.29 | 0.10 | 0.27 | 5 | 0 | 35 | 1.0 | 0.0 | 2.8 |
| **Cv2.1303** | CC1 | 3-49 | 2-2/2-2/2-2 | 5 | γ1 | GASQLLAGNWFAS | 13 | 0 | 3 | 24 | κ | 1-39/1D-39 | 3 | QQSYSTLVIT | 10 | 11 | RBD | 0.2 | 2.3 | 2.3 | 0.13 | 1.76 | 2.97 | 3.62 | 0.11 | 0.15 | 0.14 | 0.07 | 0.15 | 0.14 | 0.14 | 0.12 | 0.20 | 2 | 2 | 11 | 1.0 | 0.0 | 0.8 |
| **Cv2.1309** | CC1 | 3-7 | 6-6 | 4 | γ1 | DLDTYSSSPNDY | 12 | 2 | 1 | 9 | κ | 1-5 | 4 | QQYNSYPLT | 9 | 0 | NTD | 94.6 | 5.6 | 11.2 | 0.76 | 3.73 | 0.44 | 0.07 | 2.79 | 0.08 | 0.08 | 0.06 | 0.08 | 0.06 | 0.07 | 0.08 | 0.07 | 0 | 0 | 0 | 1.1 | 0.6 | 2.4 |
| **Cv2.1328** | CC1 | 3-30 | 5-24 | 4 | γ1 | GRGGYKTFFDY | 11 | 0 | 2 | 5 | κ | 3-20 | 4 | QQYGSLPLT | 9 | 7 | S2 | 91.2 | 5.1 | 12.9 | 2.81 | 3.40 | 0.09 | 0.08 | 0.07 | 0.08 | 3.42 | 0.06 | 2.67 | 0.06 | 0.07 | 0.20 | 0.08 | 0 | 32 | 9 | 1.2 | 0.0 | 3.9 |
| **Cv2.1330** | CC1 | 3-30 | 6-19 | 4 | γ1 | DFYSSGSSIDY | 11 | 0 | 2 | 2 | λ | 3-1 | 2/3 | QAWDSRTVV | 9 | 12 | S2 | 91.2 | 4.9 | 7.9 | 2.69 | 2.84 | 0.86 | 0.52 | 0.22 | 1.00 | 2.67 | 0.54 | 0.89 | 0.21 | 0.69 | 0.08 | 0.95 | 0 | 17 | 7 | 1.3 | 0.0 | 2.4 |
| **Cv2.1333** | CC1 | 3-33 | 2-15 | 4 | γ1 | ESQWSGGSQIDY | 12 | 0 | 2 | 8 | λ | 3-1 | 2/3 | QAWDSSVVV | 9 | 5 | S2 | 89.7 | 4.9 | 10.8 | 2.67 | 3.11 | 0.08 | 0.07 | 0.07 | 0.07 | 2.77 | 0.06 | 0.08 | 0.06 | 0.07 | 0.14 | 0.07 | 0 | 25 | 10 | 1.6 | 0.8 | 3.7 |
| **Cv2.1353** | CC1 | 3-66 | 3-3 | 3 | γ1 | DLGPMGFDI | 9 | 1 | 1 | 7 | κ | 1-9 | 3 | QQLNSDPPVT | 10 | 3 | RBD | 94.6 | 5.4 | 14.0 | 2.22 | 3.50 | 3.46 | 3.55 | 0.12 | 0.08 | 0.12 | 0.06 | 0.09 | 0.09 | 0.09 | 0.79 | 0.11 | 92 | 100 | 100 | 1.2 | 13.6 | 4.7 |
| **Cv2.1354** | CC1 | 3-30/3-30-3 | 1-1/1-20/1-7 | 6 | γ1 | ARGGNYYYGMDV | 12 | 1 | 1 | 11 | κ | 1-9 | 3 | QQLNSYPPFT | 10 | 9 | S2 | 90.4 | 5.0 | 13.4 | 2.74 | 3.36 | 0.08 | 0.08 | 0.07 | 0.07 | 3.35 | 0.06 | 2.66 | 0.07 | 0.08 | 0.16 | 0.08 | 0 | 5 | 8 | 1.1 | 0.0 | 3.8 |
| **Cv2.1357** | CC1 | 3-30 | 3-3/3-3 | 6 | γ1 | AHGGAYYYGMDV | 12 | 1 | 1 | 11 | κ | 1-39/1D-39 | 5 | QQSYSTFT | 8 | 8 | S2 | 91.9 | 5.0 | 11.9 | 2.75 | 2.97 | 0.08 | 0.07 | 0.07 | 0.07 | 2.81 | 0.06 | 1.06 | 0.06 | 0.07 | 0.14 | 0.07 | 0 | 34 | 11 | 1.3 | 0.0 | 4.6 |
| **Cv2.1380** | CC1 | 3-30 | 4-17 | 6 | γ1 | DYGDYGGMDV | 10 | 0 | 3 | 12 | κ | 4-1 | 1 | QHYYTTPT | 8 | 6 | CD | 87.1 | 4.6 | 14.1 | 2.22 | 3.53 | 3.51 | 0.07 | 0.10 | 3.57 | 0.07 | 0.06 | 0.07 | 0.06 | 0.07 | 0.48 | 0.07 | 20 | 45 | 20 | 1.0 | 0.0 | 2.2 |
| **Cv2.2154** | CC2 | 3-30-3 | 1-26 | 4 | γ1 | AYSGNYYGSDY | 11 | 0 | 1 | 9 | κ | 1-39/1D-39 | 4 | QQSYSAPWLT | 10 | 8 | S2 | 91.9 | 5.0 | 15.0 | 2.82 | 3.30 | 0.08 | 0.07 | 0.07 | 0.08 | 3.47 | 0.06 | 2.66 | 0.06 | 0.07 | 0.17 | 0.08 | 0 | 24 | 12 | 1.1 | 1.6 | 3.7 |
| **Cv2.2157** | CC2 | 3-7 | 6-19 | 3 | γ1 | DSLGPPGYSSGWRDDAFDI | 19 | 1 | 4 | 1 | λ | 3-21 | 3 | QVWDSSSDHWV | 11 | 1 | NTD | 85.8 | 5.0 | 12.0 | 0.45 | 3.35 | 0.08 | 0.07 | 3.30 | 0.07 | 0.11 | 0.06 | 0.08 | 0.05 | 0.07 | 0.07 | 0.06 | 2 | 1 | 20 | 3.1 | 11.0 | 3.3 |
| **Cv2.2180** | CC2 | 1-69/1-69D | 6-19 | 5 | γ1 | EAMAGFSWFDP | 11 | 0 | 2 | 6 | κ | 1-5 | 1 | QQYNTYST | 8 | 13 | S2 | 96.2 | 4.8 | 11.9 | 2.63 | 3.50 | 0.17 | 0.09 | 0.09 | 0.16 | 3.53 | 0.01 | 3.43 | 0.17 | 0.12 | 0.13 | 0.19 | 0 | 5 | 4 | 1.6 | 1.3 | 5.2 |
| **Cv2.2187** | CC2 | 3-9 | 6-19 | 4 | γ1 | DIAGGYSSGPCDY | 13 | 0 | 2 | 2 | λ | 3-21 | 2/3 | QVWDSSSDPHVV | 12 | 3 | RBD | 96.2 | 4.9 | 12.7 | 1.25 | 3.28 | 3.48 | 3.79 | 0.06 | 0.08 | 0.07 | 0.01 | 0.31 | 0.07 | 0.08 | 0.07 | 0.08 | 75 | 35 | 69 | 1.7 | 6.1 | 4.2 |
| **Cv2.2190** | CC2 | 3-30-3 | 1-7 | 4 | γ1 | SPGGNYYYFDS | 11 | 0 | 1 | 7 | κ | 3-11 | 3 | QQRSNWPPT | 9 | 21 | S2 | 96.4 | 4.8 | 11.8 | 2.36 | 3.30 | 0.23 | 0.13 | 0.12 | 0.24 | 3.40 | 0.01 | 1.17 | 0.32 | 0.20 | 0.24 | 0.30 | 0 | 4 | 12 | 1.1 | 0.5 | 3.8 |
| **Cv2.2216** | CC2 | 3-33 | 3-10/3-10 | 4 | γ1 | EGVFGESGIDY | 11 | 0 | 3 | 1 | λ | 3-10 | 2/3 | YSTGSSGNHRV | 11 | 6 | S2 | 95.8 | 4.7 | 10.2 | 2.43 | 2.99 | 0.08 | 0.08 | 0.07 | 0.10 | 3.40 | 0.01 | 0.10 | 0.32 | 0.20 | 0.24 | 0.30 | 2 | 11 | 15 | 1.6 | 0.9 | 4.1 |
| **Cv2.2217** | CC2 | 5-51 | 6-19 | 4 | γ1 | PRFAVAGPQGEFDY | 14 | 1 | 2 | 6 | κ | 3-20 | 1 | QHYGSSSWT | 9 | 21 | S2 | 97.4 | 4.9 | 13.6 | 2.86 | 3.36 | 0.07 | 0.07 | 0.08 | 0.09 | 4.14 | 0.01 | 1.56 | 0.08 | 0.08 | 0.08 | 0.09 | 0 | 3 | 13 | 1.7 | 3.6 | 3.3 |
| **Cv2.2221** | CC2 | 1-24 | 6-13/6-19/6-25 | 5 | γ1 | ASIAGNPTWFDP | 12 | 0 | 1 | 12 | λ | 1-51 | 3 | GTWDSNLTAVV | 11 | 9 | NTD | 96.9 | 5.2 | 13.4 | 2.17 | 3.48 | 3.53 | 0.07 | 3.50 | 0.08 | 0.08 | 0.01 | 0.09 | 0.07 | 0.08 | 0.08 | 0.10 | 0 | 0 | 53 | 3.2 | 18.4 | 4.1 |
| **Cv2.2223** | CC2 | 3-30-3 | 3-22 | 4 | γ3 | GQGGGYYSSFDY | 12 | 0 | 1 | 2 | κ | 3-20 | 2 | QQYGSSPMYT | 10 | 23 | S2 | 97.3 | 4.9 | 11.3 | 2.82 | 3.22 | 0.08 | 0.08 | 0.07 | 0.08 | 3.74 | 0.01 | 2.18 | 0.07 | 0.07 | 0.07 | 0.09 | 0 | 9 | 16 | 1.2 | 0.4 | 1.2 |
| **Cv2.2243** | CC2 | 4-39 | 2-15/2-21/2-21 | 4 | γ1 | LLWWHFDY | 8 | 1 | 1 | 3 | κ | 3-11 | 4 | QHRSNWPLT | 9 | 6 | S2 | 91.3 | 5.0 | 11.3 | 2.72 | 2.94 | 0.16 | 0.10 | 0.09 | 0.19 | 3.17 | 0.16 | 1.95 | 0.08 | 0.16 | 0.09 | 0.18 | 0 | 19 | 3 | 2.2 | 1.7 | 3.3 |
| **Cv2.2251** | CC2 | 3-33 | 1-14/3-3/3-3 | 3 | γ1 | ARNFFDAFDI | 10 | 1 | 2 | 5 | κ | 1-5 | 1 | QQYNSYPWT | 9 | 18 | S2 | 95.4 | 4.8 | 11.5 | 2.89 | 3.21 | 0.17 | 0.09 | 0.09 | 0.14 | 4.04 | 0.01 | 3.15 | 0.23 | 0.13 | 0.13 | 0.19 | 0 | 9 | 11 | 1.5 | 1.5 | 3.3 |
| **Cv2.2257** | CC2 | 4-59 | | 4 | γ1 | GFDY | 4 | 0 | 1 | 5 | κ | 3-20 | 1 | QQYGTSLWT | 9 | 17 | S2 | 95.9 | 4.7 | 13.0 | 2.64 | 3.36 | 0.26 | 0.14 | 0.11 | 0.24 | 3.89 | 0.01 | 3.33 | 0.35 | 0.21 | 0.21 | 0.32 | 0 | 14 | 20 | 3.1 | 7.8 | 4.2 |
| **Cv2.2293** | CC2 | 3-33 | 3-10 | 6 | γ1 | EMSLVRGVILPYYGMDV | 17 | 1 | 2 | 6 | κ | 1-5 | 1 | QQYNSYSWT | 9 | 14 | S2 | 95.9 | 4.8 | 12.8 | 2.73 | 3.10 | 0.11 | 0.08 | 0.08 | 0.11 | 4.10 | 0.00 | 3.17 | 0.11 | 0.11 | 0.11 | 0.13 | 3 | 6 | 18 | 2.3 | 4.5 | 1.4 |
| **Cv2.2313** | CC2 | 3-30 | 2-2 | 4 | γ1 | GMDIVVVPAALFDY | 14 | 0 | 2 | 2 | λ | 2-14 | 1 | SSYTRSSTPFV | 11 | 6 | S2 | 90.7 | 5.0 | 13.1 | 2.81 | 3.07 | 0.20 | 0.09 | 0.11 | 0.13 | 3.02 | 0.07 | 1.95 | 0.21 | 0.25 | 0.14 | 0.30 | 0 | 26 | 14 | 2.2 | 3.1 | 3.8 |
| **Cv2.2318** | CC2 | 3-30-3 | 1-26 | 4 | γ1 | PNSGSYFSYFDY | 12 | 0 | 1 | 3 | κ | 3-20 | 3 | QQYGSSSRVT | 10 | 5 | S2 | 91.0 | 5.0 | 12.7 | 2.78 | 3.08 | 0.10 | 0.10 | 0.08 | 0.10 | 3.00 | 0.07 | 1.79 | 0.06 | 0.07 | 0.13 | 0.07 | 0 | 37 | 2 | 1.1 | 0.0 | 4.1 |
| **Cv2.2333** | CC2 | 3-21 | 3-22 | 3 | γ1 | WNGGSDYYPGAFDI | 14 | 0 | 2 | 4 | κ | 3D-15 | 1 | QQYNNWPPWT | 10 | 1 | NTD | 84.3 | 4.4 | 9.8 | 0.75 | 3.42 | 1.42 | 0.09 | 3.34 | 0.09 | 0.08 | 0.07 | 0.08 | 0.07 | 0.08 | 0.07 | 0.07 | 14 | 32 | 18 | 1.1 | 1.4 | 1.6 |
| **Cv2.2346** | CC2 | 3-33 | 1-26 | 4 | γ1 | EGMVGATTGLDY | 12 | 0 | 2 | 9 | λ | 3-10 | 2/3 | HSTDSSGNHRV | 12 | 4 | S2 | 88.3 | 4.7 | 11.1 | 2.72 | 3.15 | 0.87 | 0.61 | 0.19 | 1.30 | 3.07 | 0.68 | 1.25 | 0.36 | 0.95 | 0.21 | 1.34 | 0 | 27 | 6 | 1.6 | 1.9 | 2.8 |
| **Cv2.2365** | CC2 | 3-30-3 | 1-1/1-20/1-7 | 3 | γ1 | ARRGNYQNAFDI | 12 | 2 | 1 | 13 | κ | 3-20 | 2 | QQYGSAYT | 8 | 4 | S2 | 92.4 | 5.1 | 12.5 | 2.88 | 3.17 | 0.08 | 0.07 | 0.07 | 0.08 | 3.25 | 0.06 | 1.87 | 0.06 | 0.07 | 0.17 | 0.07 | 0 | 28 | 19 | 1.1 | 0.0 | 4.0 |
| **Cv2.2376** | CC2 | 3-73 | 4-17 | 4 | γ1 | LGQEVSTVTTVLDY | 14 | 0 | 2 | 24 | κ | 1-17 | 4 | LQHSGFPLT | 9 | 14 | S2 | 90.7 | 5.0 | 5.3 | 3.27 | 1.87 | 0.10 | 0.10 | 0.07 | 0.10 | 2.72 | 0.07 | 0.76 | 2.86 | 0.13 | 3.10 | 0.07 | 0 | 31 | 14 | 2.4 | 10.0 | 4.2 |
| **Cv2.2392** | CC2 | 3-30-3/3-30-3 | 3-10 | 4 | γ1 | TGSGSYWDFDY | 11 | 0 | 2 | 8 | λ | 7-43 | 3 | LLYYGGAWV | 9 | 6 | S2 | 89.5 | 5.0 | 10.8 | 2.74 | 3.02 | 0.49 | 0.26 | 0.15 | 0.72 | 2.88 | 0.28 | 2.02 | 0.21 | 0.52 | 0.21 | 0.66 | 0 | 18 | 11 | 1.1 | 4.0 | 2.3 |
| **Cv2.2393** | CC2 | 3-9 | 6-19 | 4 | γ1 | DIGGLIAVAGYFDY | 14 | 0 | 2 | 3 | κ | 1-16 | 1 | QQYNSYPLT | 9 | 0 | S2 | 81.5 | 4.5 | 2.3 | 2.85 | 0.59 | 0.12 | 0.09 | 0.09 | 0.12 | 1.79 | 0.07 | 0.11 | 0.08 | 0.10 | 0.07 | 0.10 | 0 | 0 | 5 | 1.3 | 7.1 | 2.6 |
| **Cv2.2396** | CC2 | 3-30 | 5-12 | 6 | γ1 | GLRPLHYYYYGLDV | 14 | 2 | 1 | 10 | κ | 3-11 | 3 | QQRNNWPPGIT | 11 | 7 | NTD | 89.8 | 4.9 | 13.5 | 1.56 | 3.43 | 3.42 | 0.09 | 3.51 | 0.12 | 0.15 | 0.07 | 2.69 | 0.09 | 0.15 | 0.12 | 0.15 | 32 | 54 | 31 | 1.3 | 4.7 | 1.9 |
| **Cv2.3110** | CC3 | 4-59 | | 4 | γ1 | GFDY | 4 | 0 | 1 | 6 | κ | 3-20 | 1 | QQYGSSPWT | 9 | 4 | S2 | 95.5 | 4.7 | 11.4 | 2.63 | 2.97 | 0.08 | 0.09 | 0.09 | 0.13 | 3.57 | 0.07 | 2.62 | 0.07 | 0.09 | 0.07 | 0.07 | 9 | 17 | 7 | 1.5 | 3.2 | 3.9 |
| **Cv2.3132** | CC3 | 3-11 | 3-22 | 4 | α1 | YYYDSSGYWKRLDY | 14 | 2 | 2 | 6 | κ | 1-39/1D-39 | 4 | QQSYSTPLT | 9 | 6 | S2 | 98.7 | 4.9 | 14.1 | 2.70 | 3.18 | 0.08 | 0.09 | 0.07 | 0.08 | 3.47 | 0.01 | 2.94 | 0.07 | 0.08 | 0.08 | 0.08 | 0 | 48 | 36 | 0.9 | 0.1 | 2.6 |
| **Cv2.3144** | CC3 | 1-69 | 3-22 | 5 | γ1 | EIGYYDSSGYSGWFDP | 16 | 0 | 3 | 6 | κ | 3-20 | 4 | QQYGSSPLT | 9 | 5 | NTD | 95.8 | 4.3 | 13.5 | 1.83 | 3.16 | 3.35 | 0.08 | 3.48 | 0.08 | 0.07 | 0.01 | 0.08 | 0.07 | 0.08 | 0.08 | 0.08 | 37 | 21 | 35 | 1.2 | 1.9 | 2.0 |
| **Cv2.3155** | CC3 | 1-69/1-69D | 3-22 | 4 | γ1 | VGAYDSGGYYPEY | 13 | 0 | 2 | 7 | κ | 3-11 | 3 | QHRSNWPPLFT | 11 | 5 | S2 | 96.2 | 4.8 | 12.2 | 2.82 | 3.12 | 0.10 | 0.09 | 0.08 | 0.09 | 3.45 | 0.01 | 3.12 | 0.10 | 0.09 | 0.08 | 0.11 | 6 | 6 | 8 | 1.3 | 0.7 | 5.1 |
| **Cv2.3175** | CC3 | 3-33 | 2-8 | 6 | γ1 | EGQWSEVDGMDV | 12 | 0 | 4 | 3 | λ | 3-25 | 2/3 | QSTDSSNTYRV | 11 | 6 | S2 | 89.1 | 4.9 | 12.6 | 2.82 | 3.08 | 0.08 | 0.11 | 0.07 | 0.08 | 3.03 | 0.06 | 0.07 | 0.06 | 0.07 | 0.16 | 0.06 | 5 | 36 | 5 | 1.2 | 4.4 | 2.8 |
| **Cv2.3181** | CC3 | 3-23 | 3-3/3-3 | 4 | α2 | GRLTYDGDSHYYTVFDY | 17 | 0 | 2 | 14 | λ | 1-51 | 2/3 | GTWDNNLSVVV | 11 | 7 | Und. | 22.6 | 2.6 | 2.3 | 0.07 | 2.31 | 0.29 | 0.30 | 0.14 | 0.23 | 0.20 | 0.09 | 0.13 | 0.09 | 0.12 | 0.10 | 0.12 | 5 | 0 | 0 | 1.0 | 0.0 | 1.1 |
| **Cv2.3194** | CC3 | 3-53 | 2-2/2-2/2-2 | 6 | γ3 | DLVVYGMDV | 9 | 0 | 2 | 5 | κ | 3-20/3-NL5 | 4 | QQGVT | 5 | 7 | RBD | 94.0 | 5.4 | 15.8 | 2.25 | 3.63 | 3.66 | 3.63 | 0.12 | 0.08 | 0.09 | 0.06 | 0.10 | 0.07 | 0.07 | 0.79 | 0.08 | 91 | 100 | 100 | 1.6 | 15.0 | 4.1 |
| **Cv2.3215** | CC3 | 3-30-3 | 6-6 | 3 | γ1 | ASSMAARNDAFDI | 13 | 1 | 2 | 10 | λ | 3-21 | 3 | QLWDSSRDHWV | 11 | 9 | NTD | 97.4 | 5.0 | 13.4 | 1.60 | 3.25 | 1.90 | 0.14 | 3.59 | 0.20 | 0.17 | 0.01 | 0.26 | 0.17 | 0.16 | 0.13 | 0.21 | 0 | 14 | 78 | 1.6 | 3.4 | 3.6 |
| **Cv2.3216** | CC3 | 3-30/3-30 | 6-19 | 3 | γ1 | GAVAGTDSFDI | 11 | 0 | 2 | 25 | κ | 2-30 | 5 | MQGTHWPIT | 9 | 9 | S2 | 96.5 | 4.9 | 0.6 | 2.81 | 2.51 | 0.13 | 0.15 | 0.11 | 0.14 | 1.95 | 0.08 | 0.95 | 0.08 | 0.84 | 3.96 | 0.09 | 3 | 0 | 22 | 2.2 | 4.9 | 3.2 |
| **Cv2.3221** | CC3 | 1-69/1-69D | 3-22 | 4 | γ1 | VWAYDSRGYYLDY | 13 | 1 | 2 | 3 | κ | 3-11 | 3 | QQRSNWPPLIT | 11 | 10 | S2 | 89.9 | 5.0 | 12.8 | 2.85 | 3.28 | 0.14 | 0.08 | 0.10 | 0.12 | 3.63 | 0.08 | 1.90 | 0.10 | 0.14 | 0.22 | 0.14 | 4 | 12 | 11 | 1.2 | 5.5 | 1.0 |
| **Cv2.3229** | CC3 | 3-74 | 6-13 | 4 | γ1 | SSWPNRIDY | 9 | 1 | 1 | 4 | λ | 3-1 | 2/3 | QAWDSSTVV | 9 | 1 | S2 | 89.0 | 4.9 | 5.2 | 2.97 | 1.90 | 0.08 | 0.07 | 0.07 | 0.07 | 2.61 | 0.07 | 0.08 | 0.06 | 0.07 | 0.08 | 0.07 | 5 | 32 | 5 | 3.7 | 5.7 | 3.1 |
| **Cv2.3230** | CC3 | 1-18 | 6-6 | 1 | γ2 | EGPNSSFLY | 9 | 0 | 1 | 30 | κ | 1-12 | 5 | QQGYSFPRIT | 10 | 32 | RBD | 93.1 | 4.6 | 6.2 | 0.82 | 2.58 | 3.79 | 3.69 | 0.06 | 0.08 | 0.07 | 0.00 | 0.08 | 0.08 | 0.10 | 0.08 | 0.09 | 44 | 32 | 65 | 1.1 | 3.3 | 1.7 |
| **Cv2.3235** | CC3 | 3-53 | 3-16/3-16/7-27 | 6 | γ1 | DGDYYGMDV | 9 | 0 | 3 | 4 | κ | 1-9 | 5 | QRLDSYPPIT | 10 | 8 | RBD | 97.3 | 5.2 | 13.2 | 1.01 | 3.65 | 3.85 | 3.86 | 0.15 | 0.19 | 0.26 | 0.00 | 0.24 | 0.21 | 0.22 | 0.26 | 0.13 | 77 | 100 | 100 | 1.2 | 13.5 | 4.6 |
| **Cv2.3251** | CC3 | 3-7 | 1-1/1-20/1-7 | 4 | γ1 | LSPADWSFDY | 10 | 0 | 2 | 5 | λ | 3-1 | 1 | QAWDSSTGGV | 10 | 2 | S2 | 95.8 | 4.7 | 10.2 | 2.42 | 3.38 | 0.10 | 0.07 | 0.07 | 0.09 | 3.32 | 0.00 | 3.02 | 0.11 | 0.09 | 0.12 | 0.08 | 10 | 7 | 7 | 2.2 | 0.0 | 3.7 |
| **Cv2.3257** | CC3 | 4-39 | 3-10 | 3 | γ1 | HILLVWFGESGDSDAFDI | 18 | 1 | 4 | 4 | λ | 2-11 | 2/3 | CSYADSYTWV | 10 | 5 | NTD | 91.8 | 4.7 | 2.8 | 0.27 | 3.73 | 0.21 | 0.17 | 3.35 | 0.19 | 0.20 | 0.09 | 0.17 | 0.11 | 0.11 | 0.14 | 0.12 | 8 | 0 | 61 | 3.4 | 9.3 | 4.3 |
| **Cv2.3262** | CC3 | 3-23 | 1-1/1-20/7-27 | 4 | γ1 | ERAPLDDNDDHWALFLDFEGFDY | 23 | 2 | 8 | 9 | λ | 1-44 | 3 | AAWDDSLNGWV | 11 | 4 | S2 | 92.8 | 4.7 | 4.5 | 2.99 | 1.97 | 0.08 | 0.08 | 0.06 | 0.08 | 2.24 | 0.00 | 1.06 | 0.08 | 0.10 | 0.08 | 0.10 | 23 | 13 | 17 | 1.8 | 2.9 | 4.3 |
| **Cv2.3265** | CC3 | 3-30-3 | 1-26/1-7 | 4 | γ1 | VKGGNYQGYFDY | 12 | 1 | 1 | 6 | κ | 3-15 | 1 | QQYNNWPVT | 9 | 4 | S2 | 95.6 | 4.8 | 8.1 | 2.77 | 3.20 | 0.20 | 0.08 | 0.08 | 0.08 | 3.15 | 0.00 | 1.51 | 0.08 | 0.10 | 0.08 | 0.10 | 17 | 6 | 12 | 1.8 | 3.4 | 4.0 |
| **Cv2.3287** | CC3 | 5-51 | 6-19 | 6 | γ1 | HVMEYSSGWYYYYGMDV | 17 | 1 | 2 | 6 | κ | 3-20 | 1 | QQYGSSQGT | 9 | 4 | S2 | 93.2 | 4.2 | 9.8 | 2.23 | 2.83 | 0.06 | 0.07 | 0.07 | 0.07 | 3.15 | 0.07 | 0.15 | 0.06 | 0.07 | 0.06 | 0.06 | 2 | 23 | 8 | 1.2 | 4.6 | 3.0 |
| **Cv2.3296** | CC3 | 1-24 | | 3 | γ1 | EGI | 3 | 0 | 1 | 7 | λ | 3-10 | 3 | YSTDSSGNRWV | 11 | 2 | NTD | 96.5 | 5.2 | 9.7 | 1.07 | 3.13 | 0.14 | 0.07 | 3.58 | 0.08 | 0.08 | 0.00 | 0.08 | 0.08 | 0.09 | 0.07 | 0.09 | 5 | 0 | 33 | 3.5 | 2.4 | 3.3 |
| **Cv2.4102** | CC4 | 3-11/3-11 | 3-16/3-16/3-3 | 4 | γ1 | DSTFTDIGNFDN | 12 | 0 | 3 | 33 | κ | 1-39/1D-39 | 2 | QQSYNTPFYT | 10 | 41 | S2 | 95.3 | 4.8 | 7.2 | 2.99 | 2.56 | 0.09 | 0.08 | 0.06 | 0.08 | 3.38 | 0.00 | 3.00 | 3.92 | 3.63 | 3.71 | 0.10 | 22 | 4 | 14 | 1.6 | 0.1 | 5.2 |
| **Cv2.4117** | CC4 | 4-39 | 3-3/3-3 | 4 | γ1 | VNLQFLVDY | 9 | 0 | 1 | 22 | κ | 3-11 | 3 | QQRGNWPLFS | 10 | 22 | S2 | 95.4 | 4.7 | 7.9 | 2.74 | 1.85 | 0.08 | 0.07 | 0.11 | 0.14 | 3.28 | 0.08 | 1.52 | 3.68 | 2.44 | 3.24 | 0.07 | 12 | 19 | 6 | 1.4 | 1.8 | 3.2 |
| **Cv2.4163** | CC4 | 4-34 | 3-10 | 4 | γ1 | VGDYGSGSYYNRRSEIDY | 18 | 2 | 3 | 10 | κ | 4-1 | 1 | QQYYSTPMT | 9 | 9 | S2 | 95.4 | 4.8 | 13.1 | 2.86 | 3.49 | 0.07 | 0.08 | 0.07 | 0.08 | 3.26 | 0.00 | 2.80 | 0.06 | 0.08 | 0.07 | 0.08 | 0 | 5 | 22 | 2.2 | 3.6 | 4.8 |
| **Cv2.4171** | CC4 | 4-59 | | 5 | γ1 | ESPNNWFDS | 9 | 0 | 2 | 16 | κ | 4-1 | 4 | QQCHTRGLT | 9 | 17 | S2 | 92.4 | 4.6 | 1.7 | 3.11 | 1.04 | 0.08 | 0.07 | 0.07 | 0.08 | 2.06 | 0.00 | 0.89 | 3.66 | 3.42 | 3.69 | 0.09 | 0 | 0 | 21 | 1.4 | 1.5 | 3.3 |
| **Cv2.4209** | CC4 | 4-4 | 2-15 | 3 | γ1 | PFVGVGGAFDI | 11 | 0 | 1 | 17 | λ | 1-44 | 2/3 | AAWDDSLNGVV | 11 | 7 | S2 | 93.1 | 4.5 | 0.4 | 2.75 | 0.86 | 0.16 | 0.15 | 0.10 | 0.12 | 0.71 | 0.08 | 0.47 | 1.56 | 3.55 | 3.95 | 0.12 | 0 | 0 | 0 | 2.2 | 0.1 | 1.9 |
| **Cv2.4287** | CC4 | 3-21 | 3-3 | 4 | γ1 | VSPQIFGVVTTFDY | 14 | 0 | 1 | 20 | λ | 3-21 | 2/3 | QLWDSSSDHVV | 11 | 16 | RBD | 26.6 | 2.4 | 1.2 | 0.10 | 0.23 | 0.61 | 3.13 | 0.13 | 0.18 | 0.21 | 0.00 | 0.23 | 0.20 | 0.19 | 0.22 | 0.17 | 0 | 1 | 27 | 1.0 | 0.0 | 1.1 |
| **Cv2.5105** | FC1 | 4-4 | 6-25 | 5 | γ1 | RQSGKHNWFDP | 11 | 3 | 1 | 6 | λ | 2-23/2-23 | 3 | GSYAGSSTWV | 10 | 8 | NTD | 91.3 | 3.9 | 12.8 | 0.80 | 3.22 | 3.71 | 0.09 | 3.91 | 0.09 | 0.09 | 0.01 | 0.09 | 0.08 | 0.09 | 0.08 | 0.10 | 34 | 15 | 27 | 1.0 | 0.2 | 1.7 |
| **Cv2.5145** | FC1 | 3-30-3 | 1-26 | 6 | γ1 | AFGGSYYYGMDV | 12 | 0 | 1 | 5 | λ | 2-14 | 1 | SSYTISSTPYV | 11 | 7 | S2 | 95.9 | 4.8 | 10.5 | 2.54 | 3.35 | 0.10 | 0.11 | 0.09 | 0.11 | 3.44 | 0.01 | 3.05 | 0.09 | 0.10 | 0.08 | 0.11 | 0 | 17 | 6 | 1.1 | 1.2 | 3.6 |
| **Cv2.5164** | FC1 | 4-31 | 5-12 | 6 | γ1 | NGYRGYAPLYSYGMDV | 16 | 1 | 1 | 25 | κ | 2-28/2D-28 | 1 | MQALETPPT | 9 | 28 | RBD | 82.1 | 3.2 | 6.5 | 0.90 | 2.48 | 3.71 | 3.75 | 0.07 | 0.08 | 0.08 | 0.00 | 0.09 | 0.07 | 0.09 | 0.08 | 0.10 | 27 | 13 | 4 | 1.1 | 0.6 | 1.9 |
| **Cv2.5168** | FC1 | 1-69/1-69D | 3-22 | 4 | γ1 | TSPYDSSGYYKDY | 13 | 1 | 2 | 5 | κ | 3-11 | 4 | QQRSNWPPRLT | 11 | 23 | S2 | 97.0 | 4.8 | 11.0 | 2.75 | 3.41 | 0.18 | 0.10 | 0.10 | 0.20 | 3.43 | 0.00 | 3.42 | 0.95 | 0.18 | 0.18 | 0.19 | 13 | 17 | 9 | 1.4 | 1.7 | 5.9 |
| **Cv2.5179** | FC1 | 1-58 | 2-2/2-2/2-2 | 3 | γ3 | PHCSSTSCYDAFDI | 14 | 1 | 2 | 4 | κ | 3-20 | 1 | QQYGRSPWT | 9 | 3 | RBD | ND | ND | ND | 3.45 | 3.93 | 3.59 | 3.95 | 0.12 | 0.07 | 0.12 | 0.07 | 0.12 | 0.08 | 0.10 | 0.11 | 0.09 | ND | ND | ND | ND | ND | ND |
| **Cv2.5187** | FC1 | 3-30 | 3-10 | 4 | γ3 | AGIGRWFGFDY | 11 | 1 | 1 | 11 | λ | 3-1 | 2/3 | QAWDSSTVV | 9 | 1 | S2 | 96.7 | 4.7 | 7.9 | 2.59 | 3.01 | 0.10 | 0.09 | 0.09 | 0.10 | 2.97 | 0.00 | 3.08 | 0.08 | 0.09 | 0.09 | 0.10 | 6 | 17 | 12 | 2.3 | 0.6 | 3.0 |
| **Cv2.5213** | FC1 | 3-53 | | 6 | γ1 | DLNYYGMDV | 9 | 0 | 2 | 4 | κ | 1-9 | 3 | QQLDSYSPFT | 10 | 15 | RBD | 94.6 | 5.4 | 14.2 | 2.13 | 3.80 | 3.49 | 3.55 | 0.12 | 0.09 | 0.12 | 0.07 | 0.09 | 0.08 | 0.08 | 0.72 | 0.09 | 90 | 100 | 96 | 1.1 | 12.0 | 4.5 |
| **Cv2.5230** | FC1 | 3-30-3 | 1-26/4-23 | 6 | γ1 | GNGGSYYYGMDV | 12 | 0 | 1 | 0 | λ | 1-40 | 2/3 | QSYDSSLSGVV | 11 | 0 | S2 | 87.5 | 4.1 | 0.5 | 2.61 | 0.90 | 0.14 | 0.18 | 0.11 | 0.13 | 0.43 | 0.09 | 0.14 | 0.07 | 0.08 | 0.07 | 0.08 | 0 | 0 | 0 | 1.4 | 1.2 | 2.3 |
| **Cv2.5243** | FC1 | 1-69/1-69D | 3-22 | 4 | γ1 | VPGYDSSGYTFDY | 13 | 0 | 2 | 2 | κ | 3-11 | 2 | QQRSNWPPRYT | 11 | 11 | S2 | 96.2 | 4.8 | 5.5 | 2.95 | 2.32 | 0.11 | 0.10 | 0.09 | 0.11 | 2.93 | 0.00 | 2.60 | 0.11 | 0.12 | 0.16 | 0.12 | 13 | 3 | 2 | 2.3 | 6.3 | 4.1 |
| **Cv2.5262** | FC1 | 1-18 | 3-10 | 4 | γ3 | IVGYYYGWGSYYKEKRRFED | 20 | 4 | 3 | 19 | λ | 1-40 | 2/3 | QSSDSSLSGVV | 11 | 4 | S2 | 90.5 | 3.8 | 1.0 | 0.35 | 0.33 | 0.11 | 0.11 | 0.08 | 0.14 | 0.57 | 0.01 | 0.17 | 0.10 | 0.13 | 0.12 | 0.14 | 0 | 10 | 32 | 1.3 | 1.3 | 1.3 |
| **Cv2.5286** | FC1 | 3-53 | 4-17 | 6 | γ1 | DTGDLPYYYYGMDV | 14 | 0 | 3 | 6 | λ | 1-44 | 2/3 | AAWDDSLNGLV | 11 | 7 | S2 | 94.7 | 4.6 | 5.8 | 2.85 | 1.54 | 0.06 | 0.06 | 0.06 | 0.07 | 2.83 | 0.06 | 0.41 | 0.06 | 0.07 | 0.06 | 0.06 | 2 | 12 | 0 | 1.2 | 2.4 | 3.1 |
| **Cv2.6117** | FC2 | 1-24 | 2-21 | 4 | γ1 | WWAYCGGDCPSMFDY | 15 | 0 | 2 | 2 | λ | 1-47 | 3 | AAWDDSLSGWV | 11 | 0 | Und. | 85.2 | 4.0 | 0.3 | 0.16 | 1.50 | 0.13 | 0.15 | 0.14 | 0.11 | 0.12 | 0.08 | 0.09 | 0.07 | 0.08 | 0.12 | 0.08 | 0 | 7 | 28 | 1.1 | 0.0 | 2.6 |
| **Cv2.6136** | FC2 | 3-33 | 3-22 | 4 | γ3 | DPYDSSAYYFDY | 12 | 0 | 3 | 16 | λ | 3-21 | 3 | QVWDSSSDHWV | 11 | 12 | S2 | 89.2 | 3.8 | 4.5 | 1.84 | 2.56 | 0.32 | 0.23 | 0.10 | 0.17 | 3.05 | 0.01 | 0.30 | 0.18 | 0.37 | 0.11 | 0.41 | 5 | 5 | 8 | 1.3 | 0.1 | 1.0 |
| **Cv2.6145** | FC2 | 3-30/3-33 | 6-13 | 4 | γ1 | TYSSIWYFDY | 10 | 0 | 1 | 11 | κ | 3-11 | 4 | QQRSNWPLT | 9 | 6 | RBD | 87.0 | 3.8 | 1.3 | 0.18 | 0.98 | 3.36 | 3.57 | 0.08 | 0.07 | 0.09 | 0.01 | 0.08 | 0.07 | 0.08 | 0.08 | 0.09 | 0 | 1 | 5 | 1.0 | 0.0 | 1.4 |
| **Cv2.6182** | FC2 | 3-15 | 3-22/3-3/3-3 | 4 | γ1 | GGQWFFYFDD | 10 | 0 | 2 | 14 | λ | 3-25 | 2/3 | QSGDSGATNVV | 11 | 18 | S2 | 90.1 | 4.3 | 0.8 | 2.85 | 1.58 | 0.14 | 0.16 | 0.10 | 0.11 | 1.22 | 0.08 | 0.09 | 0.07 | 0.09 | 0.11 | 0.08 | 0 | 0 | 2 | 1.3 | 0.0 | 2.4 |
| **Cv2.6185** | FC2 | 1-69 | 3-3 | 6 | γ4 | GGAREGDFWSGYTPFYYYMDV | 21 | 1 | 3 | 12 | κ | 3-20 | 4 | QQYGNSALT | 9 | 8 | Und. | 96.7 | 4.9 | 0.7 | 0.08 | 2.92 | 1.69 | 0.30 | 2.25 | 1.08 | 1.32 | 0.11 | 0.80 | 0.90 | 1.05 | 0.87 | 0.87 | 2 | 13 | 0 | 1.1 | 0.0 | 1.0 |
| **Cv2.6234** | FC2 | 1-69 | 3-3 | 6 | γ3 | GGAREGDFWSGYTPYFYYMDV | 21 | 1 | 3 | 11 | κ | 3-20 | 4 | QHYGSSALT | 9 | 8 | Und. | 97.5 | 5.1 | 2.1 | 0.08 | 2.76 | 2.35 | 0.70 | 1.62 | 1.94 | 1.94 | 0.18 | 1.29 | 1.25 | 1.54 | 1.20 | 1.26 | 0 | 6 | 0 | 1.0 | 0.0 | 0.9 |
| **Cv2.6238** | FC2 | 3-48 | 3-3 | 4 | γ3 | LTDYDFWSGLYSYVDY | 16 | 0 | 3 | 5 | κ | 1-6 | 1 | LQDYNYPRT | 9 | 3 | NTD | 13.8 | 2.3 | 1.6 | 0.16 | 0.78 | 0.27 | 0.09 | 2.94 | 0.17 | 0.25 | 0.00 | 0.26 | 0.27 | 0.21 | 0.16 | 0.24 | 29 | 3 | 15 | 1.0 | 0.0 | 0.8 |
| **Cv2.6247** | FC2 | 3-33 | 6-13 | 4 | γ1 | GSKAGDY | 7 | 1 | 1 | 14 | κ | 1-5 | 4 | QQYNSYALT | 9 | 5 | S2 | 96.3 | 4.8 | 11.6 | 2.77 | 3.16 | 0.07 | 0.07 | 0.06 | 0.07 | 3.44 | 0.01 | 3.37 | 3.59 | 3.44 | 3.44 | 0.08 | 7 | 12 | 21 | 2.2 | 1.9 | 0.9 |
| **Cv2.6264** | FC2 | 1-69 | 3-22 | 4 | α1 | LPYYYDSSGYYYFDY | 15 | 0 | 2 | 2 | κ | 1-39/1D-39 | 1 | QQSYSNPWT | 9 | 2 | RBD | 93.2 | 5.0 | 15.2 | 1.17 | 3.44 | 3.50 | 3.67 | 0.09 | 0.14 | 0.10 | 0.00 | 0.12 | 0.09 | 0.11 | 0.08 | 0.11 | 55 | 73 | 98 | 2.9 | 10.4 | 3.7 |
| **Cv2.8109** | FC4 | 3-30-3 | 3-16/3-16/3-3 | 6 | γ1 | ALGGHYYYGMDV | 12 | 1 | 1 | 3 | κ | 3-20 | 4 | QQYGSSPALT | 10 | 13 | S2 | 96.8 | 4.8 | 8.3 | 2.88 | 2.89 | 0.09 | 0.09 | 0.07 | 0.08 | 3.43 | 0.00 | 1.85 | 0.07 | 0.10 | 0.08 | 0.10 | 0 | 7 | 1 | 1.2 | 0.0 | 2.8 |
| **Cv2.8127** | FC4 | 4-34 | 4-23 | 5 | γ2 | LDSDGVNYHWFDP | 13 | 1 | 3 | 25 | λ | 3-10 | 3 | YSTDSSGLHRV | 11 | 11 | S2 | 79.2 | 2.9 | 2.7 | 0.11 | 0.61 | 0.15 | 0.09 | 0.11 | 0.15 | 1.06 | 0.00 | 0.22 | 0.16 | 0.15 | 0.16 | 0.16 | 0 | 0 | 14 | 1.0 | 0.0 | 0.9 |
| **Cv2.8145** | FC4 | 1-24 | 6-19 | 5 | γ1 | SAVAGRKYWFDP | 12 | 2 | 1 | 3 | λ | 1-44 | 2/3 | AAWDDSLNVV | 10 | 1 | NTD | 95.3 | 5.2 | 6.3 | 0.76 | 1.94 | 0.27 | 0.07 | 2.36 | 0.08 | 0.08 | 0.01 | 0.12 | 0.07 | 0.09 | 0.07 | 0.08 | 7 | 0 | 53 | 3.5 | 6.3 | 3.2 |
| **Cv2.8162** | FC4 | 3-49 | 3-22 | 4 | γ1 | DEDYYDSSGYYPSRVDY | 17 | 1 | 5 | 3 | λ | 1-51 | 2/3 | GTWDSSLSVVV | 11 | 4 | NTD | 97.3 | 4.8 | 10.7 | 0.68 | 3.51 | 0.20 | 0.07 | 3.33 | 0.07 | 0.07 | 0.01 | 0.07 | 0.07 | 0.09 | 0.07 | 0.08 | 1 | 3 | 69 | 2.5 | 19.1 | 3.5 |
| **Cv2.8169** | FC4 | 4-30-4 | 4-11/4-4 | 6 | γ1 | DNGVQSYNYYGMDV | 14 | 0 | 2 | 14 | κ | 1-16 | 1 | QQYYSYPLT | 9 | 20 | NTD | 97.1 | 4.9 | 1.4 | 0.47 | 2.45 | 0.19 | 0.17 | 0.58 | 0.13 | 0.15 | 0.09 | 0.10 | 0.11 | 0.10 | 0.11 | 0.10 | 1 | 0 | 0 | 1.2 | 3.7 | 2.3 |
| **Cv2.8173** | FC4 | 3-30-3 | 3-16/3-16 | 3 | γ1 | PRGGIYADAFDI | 12 | 1 | 2 | 1 | κ | 3-20 | 4 | QQYGTSPLT | 9 | 3 | S2 | 91.2 | 5.0 | 6.4 | 2.98 | 1.96 | 0.07 | 0.07 | 0.07 | 0.07 | 2.68 | 0.06 | 1.40 | 0.07 | 0.06 | 0.08 | 0.07 | 1 | 24 | 12 | 3.4 | 13.2 | 4.5 |
| **Cv2.8187** | FC4 | 4-59 | 3-10/3-10 | 4 | γ1 | GFDY | 4 | 0 | 1 | 1 | κ | 3-20 | 1 | QQYGSSPWT | 9 | 24 | S2 | 94.7 | 4.8 | 13.3 | 2.53 | 3.53 | 0.22 | 0.13 | 0.12 | 0.16 | 3.39 | 0.01 | 3.20 | 0.22 | 0.25 | 0.17 | 0.21 | 3 | 12 | 11 | 2.2 | 1.4 | 4.4 |
| **Cv2.9114** | FC8 | 4-34 | 2-8 | 5 | α2 | NGRGGYWFDP | 10 | 1 | 1 | 29 | λ | 2-11 | 1 | CSYAGGYYV | 9 | 17 | RBD | 84.2 | 3.9 | 0.4 | 0.06 | 0.14 | 0.30 | 3.52 | 0.06 | 0.09 | 0.10 | 0.00 | 0.10 | 0.08 | 0.12 | 0.09 | 0.12 | 0 | 13 | 16 | 1.1 | 0.0 | 1.7 |
| **Cv2.9116** | FC8 | 3-30/3-30/3-30 | 2-2/2-2/2-2 | 3 | γ1 | ASYRSTWSLGDAFDI | 15 | 1 | 2 | 24 | κ | 1-27 | 3 | QKYNSAPFT | 9 | 12 | S2 | 95.9 | 4.8 | 5.0 | 2.53 | 2.59 | 0.09 | 0.09 | 0.06 | 0.07 | 3.03 | 0.00 | 2.66 | 3.58 | 3.34 | 3.56 | 3.37 | 15 | 5 | 16 | 2.3 | 5.1 | 6.8 |
| **Cv2.9165** | FC8 | 4-39 | 7-27 | 5 | γ1 | LKLGMGWFDP | 10 | 2 | 1 | 24 | λ | 2-23/2-23/2-23 | 3 | CSYAGSNTWV | 10 | 12 | Und. | 0.0 | 2.1 | 1.4 | 0.07 | 2.74 | 2.56 | 0.89 | 1.33 | 2.29 | 2.18 | 0.53 | 1.18 | 0.84 | 1.17 | 0.52 | 1.18 | 0 | 0 | 0 | 1.0 | 0.3 | 0.9 |
| **Cv2.9174** | FC8 | 1-69/1-69D | 3-9 | 6 | α1 | GKVIENYDILTGFYRREEYSNYYDMDV | 27 | 3 | 6 | 6 | λ | 2-14 | 3 | SSYTSSRNWV | 10 | 9 | Und. | 78.3 | 3.5 | 0.7 | 0.06 | 3.09 | 2.96 | 1.00 | 1.25 | 2.64 | 2.39 | 0.65 | 1.87 | 1.22 | 1.68 | 0.72 | 1.54 | 0 | 15 | 0 | 1.1 | 0.0 | 1.3 |
| **Cv2.9178** | FC8 | 3-30 | 6-19 | 4 | γ1 | AGSGSYQTYFDY | 12 | 0 | 1 | 3 | κ | 4-1 | 4 | QQYYSTMLT | 9 | 1 | S2 | 90.9 | 5.1 | 9.4 | 2.96 | 2.48 | 0.08 | 0.08 | 0.07 | 0.08 | 2.63 | 0.06 | 1.31 | 0.07 | 0.07 | 0.10 | 0.07 | 0 | 16 | 14 | 1.1 | 0.0 | 2.9 |
| **Cv2.9187** | FC8 | 4-4 | 6-19 | 4 | γ1 | PVSGARFDF | 9 | 1 | 1 | 19 | λ | 1-44 | 2/3 | AAWDDSLNGPI | 11 | 18 | S2 | 90.6 | 5.0 | 11.8 | 2.90 | 3.14 | 0.08 | 0.08 | 0.07 | 0.08 | 3.22 | 0.07 | 2.46 | 3.57 | 1.52 | 3.52 | 0.08 | 0 | 6 | 11 | 1.2 | 1.1 | 4.8 |
| **Cv2.9191** | FC8 | 1-2 | 1-26 | 6 | γ1 | ASYSGSAGVVWYAMDV | 16 | 0 | 1 | 13 | λ | 1-40/1-40 | 3 | QSYDNSLSGWV | 11 | 13 | RBD | 74.8 | 4.2 | 3.7 | 0.30 | 2.68 | 3.51 | 3.56 | 0.09 | 0.12 | 0.12 | 0.10 | 0.12 | 0.09 | 0.12 | 0.08 | 0.14 | 3 | 13 | 26 | 1.0 | 0.0 | 2.4 |
| **Cv2.9194** | FC8 | 1-2 | 4-23 | 6 | γ1 | YDYGGNSEKGYYYYYYGMDV | 20 | 1 | 3 | 0 | λ | 1-47 | 3 | AGWDDSLSGWV | 11 | 2 | S2 | 93.4 | 4.6 | 1.5 | 2.72 | 0.60 | 0.09 | 0.08 | 0.06 | 0.08 | 0.72 | 0.00 | 0.07 | 0.07 | 0.09 | 0.08 | 0.09 | 0 | 2 | 0 | 1.5 | 6.1 | 3.6 |
| **Cv2.10229** | FC21 | 4-31 | 1-26 | 3 | γ1 | LPGGGSYPGAFDI | 13 | 0 | 1 | 1 | λ | 3-21 | 2/3 | QVWDSNSDL | 9 | 6 | RBD | 58.6 | 3.3 | 5.4 | 0.29 | 3.27 | 3.60 | 3.65 | 0.07 | 0.07 | 0.08 | 0.06 | 0.08 | 0.06 | 0.06 | 0.07 | 0.07 | 9 | 19 | 0 | 1.0 | 4.7 | 1.1 |
| **Cv2.10234** | FC21 | 3-11 | 6-19 | 4 | γ1 | GFWDNNGWYDY | 11 | 0 | 2 | 14 | κ | 3-15 | 2 | QHYNNWPPMCS | 11 | 3 | Und. | 0.3 | 2.3 | 4.2 | 0.27 | 2.95 | 2.91 | 1.53 | 1.45 | 2.72 | 2.28 | 0.59 | 1.84 | 1.32 | 1.82 | 0.83 | 1.64 | 3 | 16 | 9 | 1.0 | 2.7 | 0.9 |

(-) and (+) indicate the numbers of negatively and positively charged amminoacids in the IgH complementary determining region (CDR3), respectively. AA, amino acid; MUT, number of somatic mutations; Und., unidentified; ND, not determined.

**Table S2. Affinity and neutralization activity of Cv2.1169 and Cv2.3194 antibodies. Related to Figures 4 and 5.**

| | | Cv2.1169 | | Cv2.3194 | |
|---|---|---|---|---|---|
| **Affinity (SPR)** | | | $K_D$ (M) | | $K_D$ (M) |
| Wuhan tri-S | ka ($M^{-1}s^{-1}$) | 6.36E+05 | | 2.45E+06 | |
| | kd ($s^{-1}$) | 1.80E-04 | 2.84E-10 | 1.12E-03 | 4.59E-10 |
| Wuhan S1 | ka ($M^{-1}s^{-1}$) | 2.46E+05 | | 4.39E+05 | |
| | kd ($s^{-1}$) | 3.54E-04 | 1.44E-09 | 5.25E-04 | 1.20E-09 |
| Wuhan RBD | ka ($M^{-1}s^{-1}$) | 1.12E+06 | | 4.87E+05 | |
| | kd ($s^{-1}$) | 7.52E-04 | 6.72E-10 | 3.06E-06 | 6.27E-12 |
| **triS/RBD-ACE2 blocking** | | $IC_{50}$ (ng/ml) | $IC_{90}$ (ng/ml) | $IC_{50}$ (ng/ml) | $IC_{90}$ (ng/ml) |
| Wuhan tri-S | | 53 | 115 | 93 | 212 |
| Omicron BA.1 tri-S | | 584 | 1314 | 876 | 1204 |
| Wuhan RBD | | 234 | 523 | 99 | 185 |
| Alpha RBD | | 295 | 601 | 212 | 737 |
| Beta RBD | | 351 | 2159 | 206 | 406 |
| Gamma RBD | | 479 | 2376 | 412 | 591 |
| Delta RBD | | 344 | 908 | 827 | 1310 |
| $Delta^+$ RBD | | 216 | 518 | 280 | 967 |
| Kappa RBD | | 168 | 446 | 469 | 1099 |
| Omicron BA.1 RBD | | 25491 | 242267 | 6530 | 37554 |
| Omicron BA.2 RBD | | 20427 | 136446 | 2713 | 29066 |
| **Pseudoneutralization** | | $IC_{50}$ (ng/ml) | $IC_{90}$ (ng/ml) | $IC_{50}$ (ng/ml) | $IC_{90}$ (ng/ml) |
| Wuhan | | 3.0 | 12.4 | 18.9 | 240.6 |
| D614G | | 4.8 | 24.0 | 40.5 | 159.8 |
| Alpha | | 5.5 | 24.0 | 34.7 | 131.9 |
| Beta | | 14.2 | 34.2 | 34.3 | 118.8 |
| Gamma | | 9.6 | 29.2 | 9.9 | 56.1 |
| Delta | | 8.0 | 34.1 | 54.4 | 114.5 |
| $Delta^+$ | | 3.6 | 14.9 | 20.6 | 89.0 |
| **S-Fuse neutralization** | | $IC_{50}$ (ng/ml) | $IC_{90}$ (ng/ml) | $IC_{50}$ (ng/ml) | $IC_{90}$ (ng/ml) |
| Wuhan | | 1.5 | 3.5 | 3.0 | 9.8 |
| D614G | | 2.1 | 5.7 | 14.5 | 145.4 |
| Alpha | | 1.6 | 4.0 | 4.3 | 5.4 |
| Beta | | 2.3 | 7.2 | 5.2 | 20.0 |
| Gamma | | 1.6 | 8.6 | 4.5 | 5.6 |
| Delta | | 2.1 | 6.4 | 6.6 | 54.7 |
| Omicron BA.1 | | 265.3 | 7592 | 24.2 | 709 |
| Omicron BA.2 | | 331.5 | 9694 | 40.5 | 582 |

**Table S3. Data collection and refinement statistics of the RBD-Fab crystallized complexes. Related to Figure 6.**

| | **RBD+Cv2.1169+CR3022** PBD: 7QEZ | **RBD+Cv2.3235** PDB: 7QF0 | **RBD+Cv2.6264** PDB: 7QF1 |
|---|---|---|---|
| **Data collection** | | | |
| Space group | $P\ 4_1$ | $C\ 2\ 2\ 2_1$ | $P\ 2_1\ 2_1\ 2_1$ |
| Cell dimensions | | | |
| *a, b, c* (Å) | 97.8, 97.8, 164.5 | 84.6, 148.9, 145.1 | 67.2, 191.3, 220.9 |
| $\alpha$, $\beta$, $\gamma$ (°) | 90, 90, 90 | 90, 90, 90 | 90, 90, 90 |
| Resolution (Å) | 48.9-2.9 (3.0-2.9) | 42.8-2.3 (2.4-2.3) | 48.2-2.8 (2.9-2.8) |
| Total reflections | 485,298 (48,609) | 561,508 (53,427) | 975,211 (98,050) |
| Unique reflections | 34,440 (3,444) | 40,941 (4,025) | 71,133 (7,013) |
| Completeness (%) | 99.8 (97.8) | 99.8 (99.6) | 99.8 (99.6) |
| Redundancy | 14.1 (14.1) | 13.7 (13.3) | 13.7 (14.0) |
| $R_{merge}$ | 0.24 (3.23) | 0.15 (1.2) | 0.73 (8.5) |
| $R_{pim}$ | 0.066 (0.887) | 0.042 (0.345) | 0.757 (8.86) |
| $I/\sigma$ ($I$) | 11.0 (0.79) | 19.7 (2.28) | 4.9 (0.25) |
| $CC_{1/2}$ | 0.996 (0.192) | 0.999 (0.767) | 0.984 (0.116) |
| **Refinement** | | | |
| Resolution (Å) | 48.9-2.9 (3.0-2.9) | 42.8-2.3 (2.4-2.3) | 48.2-2.8 (2.9-2.8) |
| No. reflections | 34,364 (3,371) | 40,936 (4.025) | 71,035 (6,988) |
| No. of reflections for Rfree | 1,785 (172) | 2,075 (193) | 3,591 (345) |
| $R_{work}/R_{free}$ | 0.24 (0.37) / 0.28 (0.37) | 0.19 (0.26) / 0.22 (0.29) | 0.23 (0.43) / 0.25 (0.44) |
| No. atoms | 6,633 | 5,076 | 9,689 |
| Protein and sugar | 6,613 | 4,810 | 9,650 |
| Ions/Buffer | 19 | 32 | 39 |
| Water | 1 | 234 | 1,251 |
| **Mean B value (Å²)** | | | |
| Protein and sugar | 91.2 | 50.6 | 91.6 |
| Ligand/Ion | 114.9 | 91.5 | 119.9 |
| Water | 51.5 | 48.2 | |
| **R.m.s. deviations** | | | |
| Bond lengths (Å) | 0.002 | 0.003 | 0.003 |
| Bond angles (°) | 0.54 | 0.62 | 0.65 |
| Ramachandran favored/outliers (%) | 95.0/0.12 | 97.2/0.0 | 97.1/0.0 |

Statistics for the highest-resolution shell are shown in parentheses.

**Table S4. Buried surface area (BSA) at the RBD-antibody interface. Related to Figure 6.**

| | Cv2.1169 | Cv2.3235 | Cv2.6264 |
|---|---|---|---|
| | BSA PARATOPE | | |
| **Heavy chain** | **537.4** | **829** | **523.5** |
| FWR1 | | 22.9 | |
| CDRH1 | 101.4 | 291 | 4.5 |
| FWR2 | 45.9 | | 48.6 |
| CDRH2 | 142.2 | 203.8 | 171 |
| FWR3 | 19 | 38.2 | 111.7 |
| CDRH3 | 228.9 | 272.9 | 187.7 |
| FWR4 | | | |
| **Light chain** | **150.5** | **520.6** | **282.4** |
| FWR1 | | 22.3 | |
| CDRH1 | 86.6 | 257.8 | 47 |
| FWR2 | | | |
| CDRH2 | 12.8 | 42.3 | |
| FWR3 | | 198.2 | |
| CDRH3 | 51.1 | | 235.4 |
| FWR4 | | | |
| **TOTAL** | **687.9** | **1349.6** | **805.9** |
| | BSA EPITOPE | | |
| **Heavy chain** | **560** | **800** | **544.5** |
| **Light chain** | **149.3** | **472** | **260.8** |
| **TOTAL** | **709.3** | **1272** | **805.3** |

**Table S5. Polar contacts at the RBD-Fab interface for the different crystallized complexes. Related to Figure 6.**

| RBD + Cv2.1169 (+CR3022) | | | RBD + Cv2.3235 | | | RBD + Cv2.6264 | | |
|---|---|---|---|---|---|---|---|---|
| PDB: 7QEZ | | | PDB: 7QF0 | | | PDB: 7QF1 | | |
| RBD residue | Fab residue | Distance (Å) | RBD residue | Fab residue | Distance (Å) | RBD residue | Fab residue | Distance (Å) |
| A: A475 (O) | H: C106 (N) | 3 | A: D420 (OD2) | H: S56 (OG) | 2.42 | A: S477 (OG) | C: Y109 (OH) | 3.58 |
| A: Q493 (NE2) | H: S55 (O) | 2.95 | A: Y421 (OH) | H: S53 (OG) | 3.26 | A: T478 (OG1) | C: Y109 (OH) | 3.11 |
| A: Y473 (OH) | H: G104 (O) | 3.13 | A: Y421 (OH) | H: S53 (N) | 3.5 | A: G485 (O) | C: N59 (ND2) | 2.56 |
| A: N487 (ND2) | H: L107 (O) | 3.07 | A: Y421 (OH) | H: G54 (N) | 3.01 | A: N487 (OD1) | C: Y108 (N) | 2.74 |
| A: T478 (N) | H: D108 (OD1) | 3.8 | A: L455 (O) | H: Y33 (OH) | 2.63 | A: S477 (N) | C: S105 (O) | 3.59 |
| A: T478 (OG1) | H: D108 (OD1) | 3.28 | A: R457 (O) | H: S53 (OG) | 2.46 | A: S477 (N) | C: Y109 (OH) | 2.95 |
| A: S477 (N) | H: D108 (OD1) | 3.72 | A: K458 (O) | H: S53 (OG) | 3.43 | A: S477 (OG) | S105 (OG) | 3.04 |
| A: S477 (OG) | H: D108 (OD2) | 3.6 | A: Q474 (O) | H: R31 (NE) | 2.7 | A: T478 (N) | C: Y109 (OH) | 3.57 |
| A: N487 (ND2) | H: D108 (OD1) | 3.73 | A: A475 (O) | H: T28 (N) | 3.11 | A: N487 (ND2) | C: Y108 (O) | 3.07 |
| A: T478 (OG1) | L: Y33 (OH) | 3.85 | A: A475 (O) | H: N32 (ND2) | 3.1 | A: Q493 (NE2) | C: G56 (O) | 3.27 |
| A: F486 (O) | L: Y33 (OH) | 3.66 | A: N487 (OD1) | H: R97 (NH1) | 2.99 | A: S477 (OG) | B: Y32 (OH) | 2.56 |
| | | | A: N487 (OD1) | H: R97 (NH2) | 2.89 | A: S477 (O) | B: Y32 (OH) | 3.88 |
| | | | A: N487 (ND2) | H: G26 (O) | 2.65 | A: F486 (O) | B: N94 (ND2) | 3.14 |
| | | | A: K458 (NZ) | H: S30 (O) | 2.83 | A: F486 (N) | B: N94 (OD1) | 2.93 |
| | | | A: Y473 (OH) | H: R31 (O) | 2.66 | | | |
| | | | A: K458 (NZ) | H: S53 (O) | 3.07 | | | |
| | | | A: Y453 (OH) | H: D100 (OD1) | 3.23 | | | |
| | | | A: K417 (NZ) | H: D100 (OD2) | 2.62 | | | |
| | | | A: K417 (N) | H: Y52 (OH) | 3.89 | | | |
| | | | A: N501 (OD1) | L: S30 (N) | 3.38 | | | |
| | | | A: G496 (O) | L: S30 (OG) | 2.86 | | | |
| | | | A: N501 (OD1) | L: S30 (OG) | 2.77 | | | |
| | | | A: G502 (N) | L: G28 (O) | 3 | | | |
| | | | A: Q498 (N) | L: S30 (OG) | 3.8 | | | |
| | | | A: Q498 (NE2) | L: S67 (OG) | 2.81 | | | |
| | | | A: Y505 (OH) | L: D92 (OD1) | 3.1 | | | |
| | | | A: R403 (NH2) | L: D92 (OD2) | 3.33 | | | |
| | | | A: K417 (NZ) | L: D92 (OD2) | 2.82 | | | |
| | | | A: R408 (NH1) | L: Y94 (OH) | 2.94 | | | |
